# Fast and efficient root phenotyping via pose estimation

**DOI:** 10.1101/2023.11.20.567949

**Authors:** Elizabeth M. Berrigan, Lin Wang, Hannah Carrillo, Kimberly Echegoyen, Mikayla Kappes, Jorge Torres, Angel Ai-Perreira, Erica McCoy, Emily Shane, Charles D. Copeland, Lauren Ragel, Charidimos Georgousakis, Sanghwa Lee, Dawn Reynolds, Avery Talgo, Juan Gonzalez, Ling Zhang, Ashish B. Rajurkar, Michel Ruiz, Erin Daniels, Liezl Maree, Shree Pariyar, Wolfgang Busch, Talmo D. Pereira

**Affiliations:** Salk Institute for Biological Studies, La Jolla, CA 92037 United States of America

## Abstract

Image segmentation is commonly used to estimate the location and shape of plants and their external structures. Segmentation masks are then used to localize landmarks of interest and compute other geometric features that correspond to the plant’s phenotype. Despite its prevalence, segmentation-based approaches are laborious (requiring extensive annotation to train), and error-prone (derived geometric features are sensitive to instance mask integrity). Here we present a segmentation-free approach which leverages deep learning-based landmark detection and grouping, also known as pose estimation. We use a tool originally developed for animal motion capture called SLEAP (Social LEAP Estimates Animal Poses) to automate the detection of distinct morphological landmarks on plant roots. Using a gel cylinder imaging system across multiple species, we show that our approach can reliably and efficiently recover root system topology at high accuracy, few annotated samples, and faster speed than segmentation-based approaches. In order to make use of this landmark-based representation for root phenotyping, we developed a Python library (*sleap-roots*) for trait extraction directly comparable to existing segmentation-based analysis software. We show that landmark-derived root traits are highly accurate and can be used for common downstream tasks including genotype classification and unsupervised trait mapping. Altogether, this work establishes the validity and advantages of pose estimation-based plant phenotyping. To facilitate adoption of this easy-to-use tool and to encourage further development, we make *sleap-roots*, all training data, models, and trait extraction code available at: https://github.com/talmolab/sleap-roots and https://osf.io/k7j9g/.

## 1. Introduction

Plant roots play a pivotal role in the survival and growth of nearly all terrestrial plants. They are responsible for water and nutrient uptake, anchoring the plant, and can serve as storage organs. Given these crucial functions, roots are key players in stress resilience and adaptation to varying environments. Their ability to transfer carbon into the soil [1], positions them at the nexus of climate change and its mitigation. It is thought that enhancing plant root traits that lead to more of their fixed carbon being stored in the soil for a longer time can contribute to substantial mitigation of climate change [2]. To identify genes and genetic variants that can be harnessed to develop plants with superior root systems, and to pinpoint existing plant varieties with these traits, large-scale phenotypic screens of plants are necessary [3].

High-throughput plant phenotyping systems allow for the acquisition of very large image datasets. For instance, the Root Architecture 3-D Imaging Cylinder (*RADICYL)* high-throughput gel system for root phenotyping allows us to capture images of the 3D root system architecture (RSA) of hundreds of plants daily using machine vision cameras and an automated rotating stage [4]. The system facilitates acquiring comprehensive and detailed images during the early stages of plant development (typically up to two weeks, depending on the crop/variety), while maintaining a relatively short overall experiment duration. Our primary objective is to identify and characterize young root phenotypes at both individual and root systems levels traits that can predict root architecture traits of mature plants. This technique can be used in tandem with lower-throughput methods, such as those involving soil or older plants, to validate findings [5]. It significantly empowers low-cost, high-throughput phenotyping in a controlled environment [6–8].

In order to automate the analysis of root images and subsequent RSA phenotypic trait extraction, most previous approaches have relied on image segmentation [9]. This technique first separates foreground plant root pixels from the background, requiring time-consuming manual adjustment of thresholding parameters and regions of interest in order to achieve reliable segmentation [10,11]. Once segmentation masks are obtained, ’skeletonization’ can be applied, an algorithm which recovers the center line of the plant roots which can be used to extract geometric (e.g., root lengths) and topological (e.g., branching angles) traits [12]. This process is highly sensitive to errors in the image segmentation step as even minor failures may result in fragmented or merged roots. This issue is tolerable in smaller-scale settings, it can become intractable in large-scale screens and across multiple species due to variability in imaging conditions and morphology.

The adoption of deep learning has drastically improved the accuracy of root segmentation [9,13–17]. Despite these advances, two issues remain: (i) annotation for segmentation is intrinsically laborious as pixels must be carefully annotated; and (ii) downstream skeletonization and instance segmentation have a very low tolerance to errors. Recent approaches have been developed to mitigate the annotation labor issue by using human-in-the-loop annotation (e.g., RootPainter [16]) or synthetic data [18]. These are still subject to the skeletonization and instance segmentation challenges, however. To address these, other methods have resorted to object detection [19], or hybrid architectures that combine segmentation and landmark detection [13,20]. These approaches either sacrifice the richness of inferring the full plant morphology in lieu of landmarks, or still require segmentation as a complementary step in processing.

Recently, it has been demonstrated that multi-instance pose estimation, a technique for landmark localization and grouping, can be used to directly extract plant morphology [21]. Unlike hybrid approaches [13], pose estimation is completely segmentation-free, thereby circumventing the issues of annotation labor and sensitivity to errors. Despite these advantages, pose estimation-based approaches have not been systematically evaluated in the context of plant RSA phenotyping.

Here we describe a pipeline for plant root pose estimation (segmentation-free landmark detection and grouping) and downstream RSA trait extraction. For pose estimation, we leveraged SLEAP, a deep learning-based framework for landmark detection and grouping originally developed for animal motion capture [22]. As most RSA trait estimation tools are designed for segmentation-based inputs, we also developed *sleap-roots*, a Python-based package for pose-based RSA trait extraction, which produces up to 1035 traits per plant. We applied our approach to a range of plant species, including crop plants such as soybean *(Glycine max*; **Supplementary Video 1**), rice *(Oryza sativa*; **Supplementary Video 2**), canola *(Brassica napus*; **Supplementary Video 3**), and the model plant Arabidopsis *(Arabidopsis thaliana*; **Supplementary Video 4**). Our results show that our approach is highly accurate (0.3 mm to 2.3 mm root landmark localization error) and efficient (peak accuracy at 10 to 200 labeled images), depending on the morphological complexity of the plant. We find that our method outperforms segmentation-based approaches in terms of annotation speed (∼1.5x faster), training (∼10x faster) and prediction (∼10x faster), at the same or better accuracy. We further validate the correctness of our trait extraction pipeline, demonstrating that it is highly accurate when compared to Fiji-based manual trait annotation (R^2^ = 0.980-0.998) or traits computed from manually proofread landmarks (up to 99.5% of data points within 1 s.d.). Finally, as a proof-of-principle, we show that the traits derived from our pipeline can be used for genotype classification and unsupervised phenotypic trait space visualization. We make all code, labeled data, and trained models available at: https://github.com/talmolab/sleap-roots and https://osf.io/k7j9g/.

## 2. Materials and Methods

### 2.1 Overview

We developed a root trait extraction pipeline specifically designed for images from the *RADICYL* phenotyping system, utilizing the SLEAP software for landmark detection (**Fig. 1**). The plants were cultivated in three-dimensional, controlled environments using transparent cylindrical containers, which are referred to as “cylinders”. 72 images for each plant were captured in 5° steps, for a comprehensive 360° view of the root system within the transparent medium. Following the imaging, the resultant images were compressed into Hierarchical Data Format, version 5 (HDF5) files [23]. This provided both portability and easy importation into the SLEAP software.

**Fig. 1.**
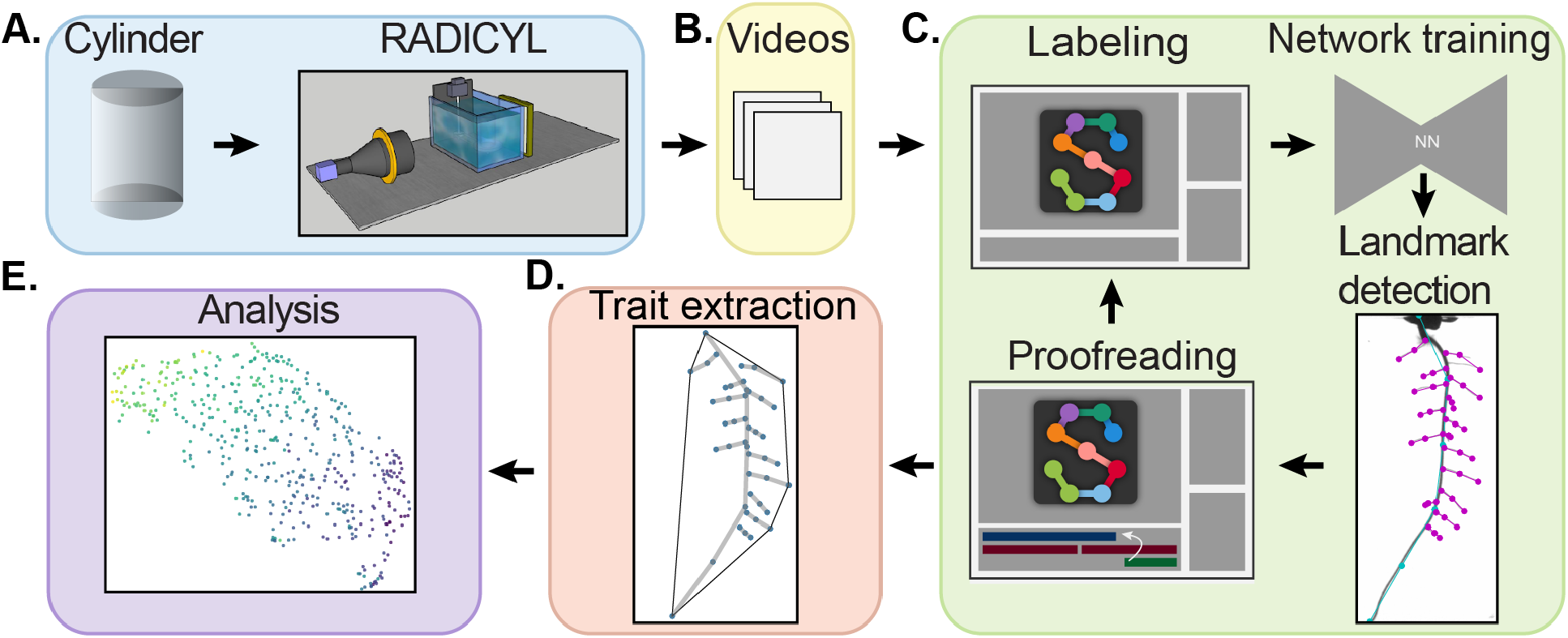
Schematic of the high-throughput phenotyping pipeline using SLEAP. This figure provides a step-by-step visual representation of our experimental design. (A) Beginning with plant cultivation in cylinders, it proceeds to the imaging process using the *RADICYL* imaging system. (B) Next, the images are compressed into HDF5 files and imported into the SLEAP software. (C) The user inputs labels of roots to train a model for root landmark detection. Predictions are made and the user refines the labels by correcting predictions (proofreading). These steps are completed in the SLEAP GUI. Corrected predictions can be used for (D) root trait extraction and included in the next training set for model improvement. (E) Efficient root trait extraction permits the computation of many traits per plant, over many plants, improving the performance of machine learning algorithms for data analysis.

The SLEAP workflow incorporated an iterative process of labeling, training, and correction (https://sleap.ai/tutorials/tutorial.html). This was designed to establish a diverse training set and to enable accurate identification of root instances and classes through landmark detection. Following the initial prediction process, the results were reviewed and refined within the SLEAP graphical user interface (GUI) which we call “proofreading”.

Two root trait extraction pipelines were made: one is designed for younger monocot plants with one primary and seminal roots, the other is for dicot plants with one primary and lateral roots. The younger monocot pipeline computes 102 traits per frame and a total of 918 derived traits per plant. The dicot pipeline computes a total of 115 traits per frame and a total of 1035 summary traits per plant. Outliers were identified by employing Principal Component Analysis (PCA) [24] and the Mahalanobis distance [25]. We utilized the high-dimensional dataset for genotype classification using the Random Forest algorithm [26] and SHAP [27] for feature selection, aimed at identifying heritable traits that could effectively differentiate genotypes. Unsupervised exploratory data analysis techniques, such as PCA, and Uniform Manifold Approximation and Projection (UMAP) [28] were implemented to visualize the phenotypic trait space.

### 2.2 Plant Cultivation

#### 2.2.1 Seed Collections

To train and evaluate the potential applications of SLEAP, we prepared datasets of cylinder images from several dicot species—soybean *(Glycine max)*, canola *(Brassica napus)*, and Arabidopsis *(Arabidopsis thaliana)*—and one monocot species, rice *(Oryza sativa* ssp. *japonica)*. These species were chosen due to their distinct early root system architectures.

For rice, we employed neutron-induced mutant Kitaake strains provided by the lab of Pamela Ronald at the University of California, Davis [29]. The soybean lines were sourced from a USDA strain collection maintained by the lab of Henry Nguyen at the University of Missouri [30]. Canola strains were obtained from the laboratory of Michael Stamm at Kansas State University [31]. Lastly, the Arabidopsis seeds came from the TRANSPLANTA collection [32].

#### 2.2.2 Environmental Conditions

The crop plants - soybean, rice, and canola-were cultivated in a greenhouse located in Encinitas, CA. The plants were organized using a completely randomized block design, and a barcode system was employed to monitor and track each individual plant throughout the study. Arabidopsis plants were grown in growth chambers at a temperature of 21°C during the daytime and 16°C during the nighttime, with a photoperiod of 16 daytime hours, and relative humidity of 60%.

#### 2.2.3 Soybean

Soybean seeds underwent vapor-phase sterilization in an airtight container for 16 hours. This was achieved using a sterilizing solution composed of 200 mL of 8.25% bleach and 3.5 mL of 1M HCl. The solution was agitated on a stir plate within the container to produce 0.0035 mol of Cl2 gas. The plants were cultivated in a medium consisting of ½ strength Murashige and Skoog (MS) medium (Sigma-Aldrich, St. Louis, USA) and 0.8% Phytagel (Sigma-Aldrich, St. Louis, USA) with a pH of 5.7. Gel cylinders were filled with 170 mL of media and were allowed to solidify at room temperature for 2 hours and could be stored at 4°C for up to 3 weeks. In sterile conditions, the sterilized seeds were sown approximately 1 mm beneath the gel media’s surface. A small hole was created in the gel to accommodate each seed. The hilum of the seed was oriented downward in the gel. The plants were then transferred to the greenhouse for growth. Imaging of the cylinders was conducted 6 days after germination (DAG).

#### 2.2.4 Rice

The seeds were first dehusked and then sterilized in a 2.475% bleach solution for 30 minutes. After sterilization, they were rinsed thrice with sterile water and soaked in sterile water for an additional 5 minutes. The water was then drained, and the sterilized seeds were carefully transferred to wet filter paper placed on sealed germination plates, ensuring all procedures were conducted under sterile conditions. These seeds were incubated on the wet filter paper at 28°C for 2 days. Seedlings of a similar developmental stage were selected and cultivated in cylinders using the same protocol as described for soybean, with the radicle oriented downwards. Rice plants were cultivated in the greenhouse and grown for 10 DAG with the day of planting in the cylinders considered as the starting point. Imaging of the cylinders was conducted at both 3 DAG and 10 DAG.

#### 2.2.5 Canola

Canola seeds underwent vapor-phase sterilization following the same protocol as soybean. Subsequently, the seeds were liquid-phase sterilized following the same protocol as rice. The water was then replaced, and the seeds were stored at 4°C for 72 hours. Next, seeds were sown on plates filled with 45 mL of 0.01% plant preservative mixture (PPM) (Caisson Labs, Smithfield, USA) in Hoagland growth media. The medium consisted of ½ strength Hoagland (MP Biomedicals, Solon, USA) and 1.0% Phytagel, adjusted to a pH of 5.8. Pregermination took place on these plates in the greenhouse over a span of 24 to 48 hours. Once the root tip emerged from the seed coat, the germination date was noted, and the seedling was transferred to a cylinder under sterile conditions. This time-point was marked as 0 (DAG). For planting, a small hole was created in the gel medium, allowing for half of the seed to be submerged. While ensuring sterility, the seedlings were planted with their root tips oriented downward in the cylinders. Canola plants imaged from 5-13 DAG were included in the training set.

#### 2.2.6 Arabidopsis

Arabidopsis seeds underwent a stratification process for 48 hours at 4°C. For sterilization, we employed vapor-phase sterilization using a mixture of 200 mL bleach and 5 mL concentrated HCl in a sealed container for one hour. Seeds were pre-germinated on plates and subsequently grown in cylinders in a medium containing ¼ strength MS, 1% sucrose and 1% Phytagel, adjusted to a pH of 5.7. Imaging of the cylinders took place 7 days after planting in cylinders.

### 2.3 Data Acquisition

The cylinder imaging system has a Basler acA2000-50gm GigE camera which produces gray-scale images with a resolution of 2048 px X 1088 px. The camera is equipped with a 0.093X, 2/3” C-Mount TitanTL® telecentric lens and placed so that the plant is in the field of view and directly in front of the camera. The cylinder is placed on a rotating stage inside of an aquarium that has a constant backlight with a red filter. The stage rotates 5°, 72 times. An image is captured at every new angle, resulting in 72 frames per plant. The camera, aquarium, and backlight are mounted on an optical breadboard for consistent positions. The rate of image acquisition is 72 frames / 7 s. Each image is 2165 KB. The scale is 10.6 px /mm.

Cylinders are 110 mm high with a diameter of 68 mm (VWR International, Radnor, USA) and filled with 170 mL of media. Cylinders are made of clear polystyrene.

### 2.4 Data Annotation

For each plant, a set of 72 images was archived in the HDF5 file format. These files were subsequently compressed utilizing lossless GZIP compression, set at a compression level of 1. To ensure unbiased annotation, each HDF5 file was named based on the unique barcode identifying each cylinder. This naming convention not only facilitated the alphabetical arrangement of plants but also shielded the annotator from potential biases that might arise from prior knowledge of the plant’s identity or genotype.

For each crop and root type, specific models were trained as detailed in **Table 2**. To bolster the training set’s diversity, plants were sourced at random from broader screenings of the pertinent seed collections.

**Table 1.** Summary of experimental conditions for each crop.

| Crop | Experimental duration | Photo-period (daytime hrs) | Average daytime temp. (°C) | Average nighttime temp. (°C) | Average daytime relative humidity (%) | Average nighttime relative humidity (%) |
| --- | --- | --- | --- | --- | --- | --- |
| Soybean | May-August, 2021 | 13 | 26.59 | 22.26 | 57.80 | 68.21 |
| Rice | April-May, 2022 | 13 | 33.81 | 25.52 | 44.65 | 65.54 |
| Canola | September-November, 2022 | 16 | 22.65 | 18.56 | 63.02 | 76.34 |

**Table 2.** Datasets used for training each model.

| Model name | Plant species | Root type | Age (DAG) | Number of plants | Number of labeled frames | Skeleton (number of nodes) |
| --- | --- | --- | --- | --- | --- | --- |
| Soybean primary roots (5-8 DAG) | *Glycine max* | Primary | 5-8 | 226 | 1389 | 6 |
| Soybean lateral roots (5-8 DAG) | *Glycine max* | Lateral | 5-8 | 220 | 482 | 4 |
| Canola primary roots (5-13 DAG) | *Brassica napus* | Primary | 5-13 | 69 | 335 | 6 |
| Canola lateral roots (5-13 DAG) | *Brassica napus* | Lateral | 5-13 | 65 | 390 | 3 |
| Arabidopsis primary roots (7 DAG) | *Arabidopsis thaliana* | Primary | 7 | 25 | 50 | 6 |
| Arabidopsis lateral roots (7 DAG) | *Arabidopsis thaliana* | Lateral | 7 | 131 | 276 | 4 |
| Rice primary root (3 DAG) | *Oryza sativa* | Primary | 3 | 134 | 748 | 6 |
| Rice seminal roots (3 DAG) | *Oryza sativa* | Seminal, Primary | 3 | 193 | 868 | 6 |
| Rice seminal roots (10 DAG) | *Oryza sativa* | Seminal, Primary | 10 | 313 | 586 | 6 |

**Table 3.** Summary of proofread data subsets.

| Dataset | Number of plants | Number of genotypes | Timepoints (DAG) |
| --- | --- | --- | --- |
| Soybean | 78 | 10 | 6 |
| Canola | 108 | 10 | 7 |
| Rice | 298 | 17 | 3 |

Root annotation within images was conducted using the SLEAP labeling workflow, in accordance with the guidelines provided in SLEAP’s online documentation (https://sleap.ai). For every video, labelers were presented with uniformly distributed labeling suggestions, 20 per video. They were directed to annotate two frames for each plant: an initial frame and a mid-time series frame, providing different plant viewpoints. Given that cylinders are symmetrically placed on the stage devoid of any bias, the selection of the first frame was effectively randomized.

Each visible root was categorized as an individual instance, symbolized by a tree structure with nodes dictated by the skeleton. This skeleton is a path, where every node is linked to precisely two others, barring the two terminal nodes which had a singular connection (**Fig. 2**). The nodes are named {r1, r2, …, rN}, with ’N’ signifying the number of nodes in the root’s skeleton. The determination of ’N’ for each crop and root type was approached with care, balancing the trade-off between accuracy and efficiency. Specifically, ’N’ was identified as the minimum number of nodes necessary to aptly depict the root’s curvature. While a higher number of nodes allows for a more precise match to the root, excessive nodes in the skeleton can inadvertently introduce errors and prolong the labeling process. We opted for a consistent number of nodes: 6 for primary and seminal roots, and 3-4 for lateral roots. The starting point, labeled as r1, represents either the root’s commencement or the highest visible point ascending the root. Nodes were evenly spaced along each root, with root tips defining their extremities. If the root tips or bases weren’t visible within the image, their visibility markers were deactivated. It’s important to note that no breaks in root visibility were permitted.

**Fig. 2.**
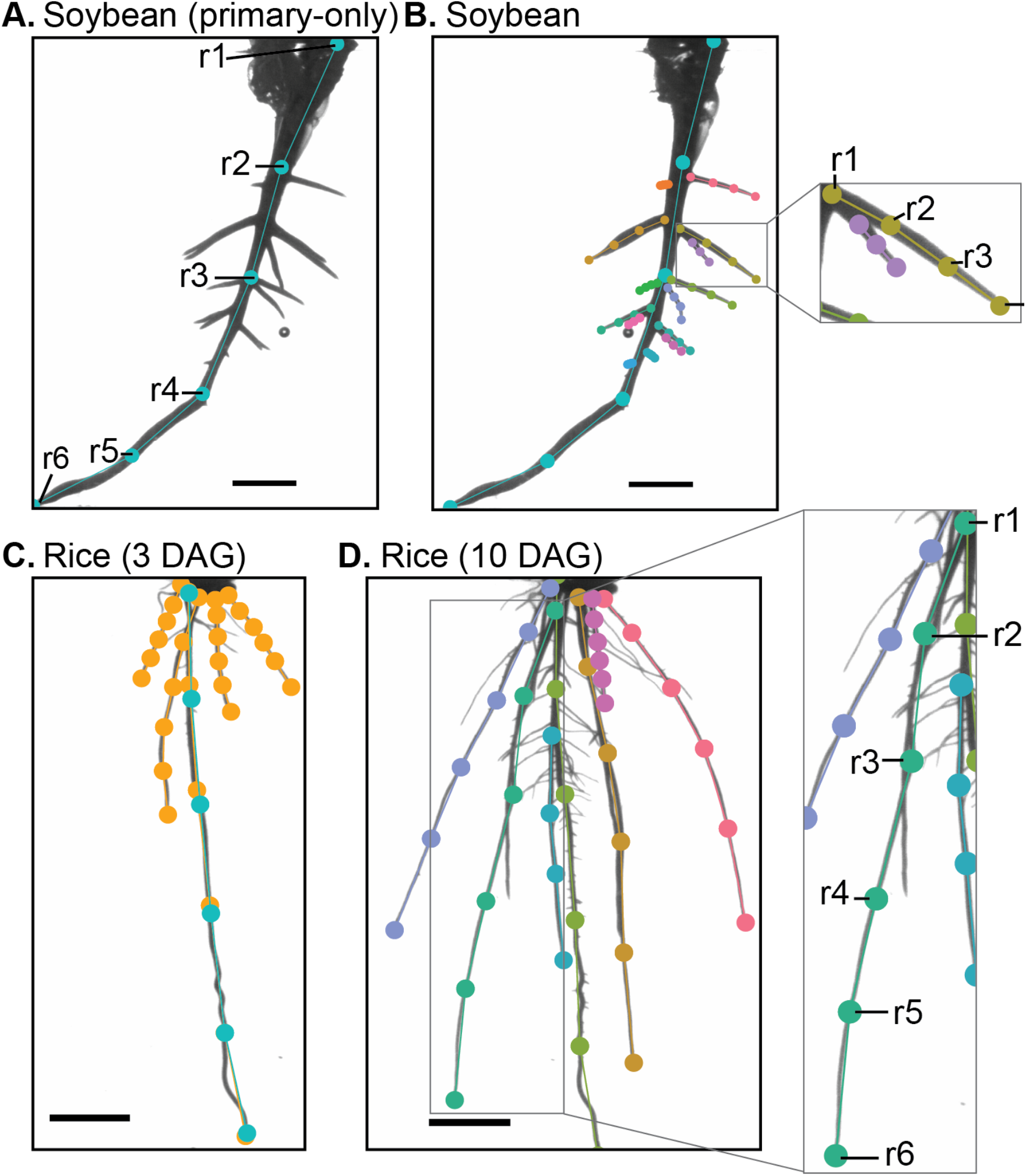
Plant and root-type labeling rules to ensure consistency. (A) Soybean, similarly to the other dicots, has a prominent primary root that can be labeled from beginning to end. (B) Lateral roots are frequently occluded and require the visibility of the first node to be toggled off. Rice roots have seminal roots which are difficult to differentiate from the primary root. For 3 DAG rice roots (C) we labeled a model for all seminal and primary roots, shown in orange, and a model only for the primary root, shown in cyan. (D) 10 DAG suffer from heavy occlusions and primary roots closely resembling seminal roots. In this case, only one class was given to all the primary and seminal roots and careful labeling rules were employed. The scale bar is 1 cm.

To enhance the precision of model predictions, rigorous labeling protocols were established for each model. In the dicots –soybean, canola, and Arabidopsis– a distinct primary root emerges, which is labeled from its beginning to its end (from r1 to rN) as depicted in **Fig. 2A**. Very slender roots (common in canola and Arabidopsis) at the bottom of the cylinder were left unlabeled as they were typically obscured. For such instances, the tip, or rN, was marked at the final discernible point–where the primary root met the cylinder’s base. Contrarily, the soybean has a thicker primary root that remains visible even at the cylinder’s bottom, facilitating its labeling. It is worth noting that lateral roots often overlap, especially when their branching points are proximate (**Fig. 2B**). In overlapping scenarios, the more pronounced lateral roots in the frame took precedence, while the visibility of obscured root bases was toggled off.

Rice, being a monocot, presents its own set of challenges. Multiple roots emerge from the radicle, leading to complications in labeling and detection due to the overlapping nature of these seminal roots at their bases. By 3 DAG, the primary root is easily distinguishable given its length, further characterized by lateral roots branching out, unlike the seminal roots. For this stage, the primary and seminal roots were collectively labeled under the model titled “Rice seminal roots (3 DAG)” in **Table 2**. Root traits associated with this class of roots are referred to as “main” root traits for brevity. Additionally, the primary root received individual labeling to allow for a more intricate extraction of its specific traits (**Fig. 2C**). By the time 10 DAG is reached, the distinction between primary and seminal roots becomes subtler since most roots have now extended to the cylinder’s base with lateral roots sprouting from them (**Fig. 2D**). As a result, one unified model was created for both primary and seminal roots, designated as “Rice seminal roots (10 DAG)” in **Table 2**. For both the 3 DAG and 10 DAG seminal root models, any base occlusion of multiple roots was addressed either by designating the root’s base as the most visible point on its upward trajectory or by deactivating the visibility of the base nodes. In the “Primary root 3 DAG” model, labelers emphasized marking the primary root up to its genuine base.

To ensure uniformity and facilitate labeling, comprehensive annotation protocols coupled with illustrative images of labeled plants were distributed among all annotators for every project. These resources are made available in the **Supplementary Materials**.

Following the conclusion of an annotation session, the model underwent training. Alternatively, it was amalgamated with other labeling initiatives and subsequently trained. This empowered the labeler to proceed with prediction-assisted annotation, a technique aiming to apprise the labeler about the model’s error types, identify where the training set needed more examples, and decrease annotation time in line with the expanding training dataset. Prediction scores in the SLEAP GUI were used to quickly identify and correct poor predictions.

### 2.5 Pose Estimation

We use the bottom-up approach of pose estimation implemented in SLEAP to perform landmark-based localization of each root. The bottom-up approach was chosen as it has been demonstrated to be successful in localizing plant landmarks with few labels [21]. In contrast to SLEAP’s top-down approach which performs better with locally distinguishable morphological features, bottom-up models explicitly learn the relationship between landmarks [22]. This is especially helpful in this application as individual landmarks along the length of the root may be better identified using their geometry as opposed to local appearance.

The root landmarks used are branch points, root tips, and equally spaced nodes in between the beginning and end of each root. Models take full-resolution raw images as input, which are processed by a deep neural network to predict multi-part confidence maps. The local peaks of the confidence maps define the locations of the landmarks (**Fig. 3, top**). The neural network simultaneously predicts part affinity fields (PAFs)–vector fields defined between connected landmarks [33]. PAFs enable scoring of connections between landmarks, which are then used to match landmarks belonging to the same root (**Fig. 3, middle**). After forming all possible connections from detected landmarks, the assembled sets of detections form instances of distinct roots (**Fig. 3, bottom**).

**Fig. 3.**
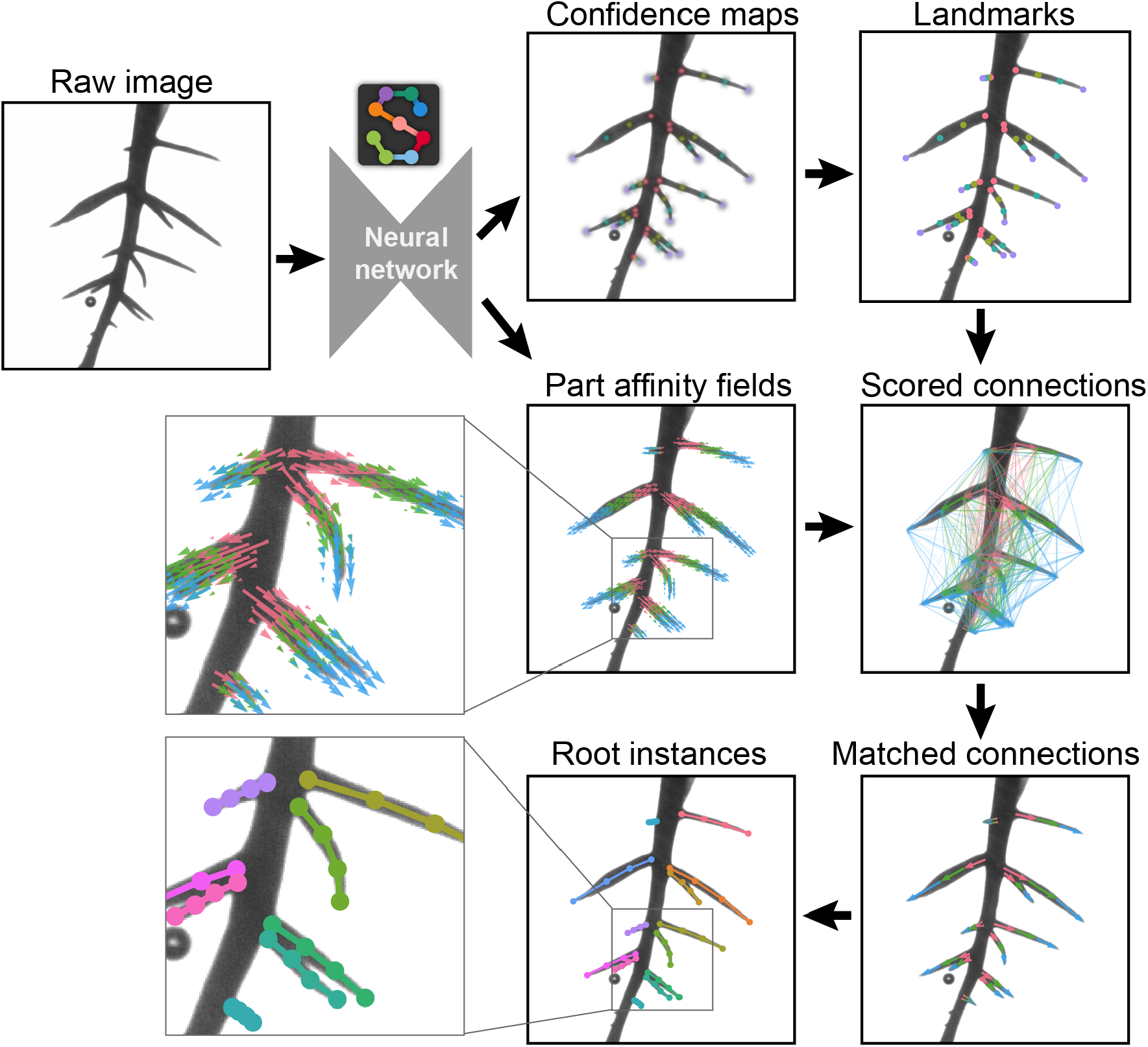
Demonstration of the bottom-up method for root pose estimation. A detailed image of a root system cultivated in a clear gel cylinder is input into a trained neural network via SLEAP. This network concurrently produces confidence maps and part affinity fields. The confidence maps, with distinct color codes for each landmark type (*r1* in pink, *r2* in yellow, *r3* in green, *r4* in purple), pinpoint probable root landmark positions. These are then sharpened into discernible peaks based on the highest probabilities within each map. Part affinity fields encode part-to-part associations in 2D vector fields across the image. Using this data, connections between root landmarks are evaluated and scored. The most highly scored connections serve to pair the root landmarks, culminating in the assembly of individual root instances, each distinguished by a unique color.

All models were trained using the same hyperparameters using SLEAP v1.3.0. Neural networks used a UNet-like architecture with 3.8 million trainable parameters, 6 downsampling blocks, 5 upsampling blocks, 24 initial convolutional filters, expanding/contracting at a rate of 1.5 per block (respectively). Downsampling blocks apply 2 convolutions with ReLU activations followed by max pooling with a stride of 2. Upsampling blocks use bilinear interpolation with 2 convolutions with ReLU activations. All convolutional layers (including pooling) use a kernel size of 3. The max stride of the model is 64, with a final output stride of 2 for confidence maps and 8 for part affinity fields. Both heads are weighed equally and trained against a mean squared error loss. Optimization is performed using the Adam optimizer with AMSgrad enabled. The only augmentations applied were random rotations of -5 to 5 degrees, however images are always downscaled by a factor of 0.5 (1024 x 544 final size) at both training and inference time. Models were trained with a batch size of 4 images per step with an initial learning rate of 1×10^-4^, which was reduced by a factor of 0.5 once the validation loss failed to decrease by at least 1×10^-6^ for at least 5 epochs, down to a minimum learning rate of 1×10^-8^. Training continued for up to 200 epochs, where an epoch is defined as the number of batches in the training set or 200 batches, whichever is larger, repeating training set images if there were fewer than 200. Shuffling was applied at the image-level to produce diverse batches of images. Training was terminated early once the validation loss failed to decrease by at least 1×10^-6^ for at least 10 epochs. The model checkpoint with the best validation loss was selected for all subsequent evaluation and inference.

### 2.6 Proofread Data

Proofreading entails a review of SLEAP predictions to ensure accurate extraction of root trait data. Given that the user will simultaneously observe the plant within the cylinder, a quality control (QC) step has been integrated into this process.

For uniform quality control both between and within individual phenotypers, a set of predetermined quality control codes has been developed. These codes pertain to specific rules governing plant exclusion. The primary criterion for exclusion is contamination. While the cylinder media is nutrient-rich, and microbial and fungal species might persist post-sterilization, slight contamination is permissible. Nevertheless, if the contamination significantly impedes clear observation of the root system architecture (RSA), the plant will be excluded (coded as “cont”). Additional exclusion criteria include the plant being fully submerged (coded as “sub”), oriented incorrectly (coded as “ori”), poor growth (coded as “pg”), and plant death (coded as “dead”).

Once a plant passes the QC stage, the phenotyper proceeds to inspect the predictions in the SLEAP GUI. Common corrections involve the detection of non-root segments in the image, such as media or cylinder deformities, or oversight of prominent RSA features. Upon making the necessary corrections, the phenotyper saves them, and the refined root trait data is then extracted from this proofread dataset.

The proofreading stage also provides phenotypers with the opportunity to discern recurring error patterns produced by the model. Identifying such consistent errors can be instrumental in model refinement. Any such instances are carefully labeled according to established protocols and subsequently incorporated into the training dataset. To foster collaboration and information exchange among phenotypers, a cloud-hosted spreadsheet was used. This spreadsheet is organized with columns for the phenotyper’s initials, QC outcomes (binary: accepted or rejected), specific QC codes, and the frames earmarked for inclusion in the training set, with each row corresponding to an individual plant.

### 2.7 Trait Extraction

The code repository has a modular design, allowing flexibility and adaptability, making it suitable for a wide range of root analysis tasks (https://github.com/talmolab/sleap-roots). *sleap-roots* can be installed with all of its dependencies using a pip package (https://pypi.org/project/sleap-roots/).

#### Series Extraction

In the *sleap_roots.series* module, the *Series* class is central to managing and analyzing data related to a single image series of root networks. This class is constructed using the *attrs* package (v23.1.0) and is designed to handle data and predictions associated with an image series. The class attributes include paths to the HDF5-formatted image series (*h5_path*), primary root predictions (*primary_labels*), lateral root predictions (*lateral_labels*), and a video representation of the image series (*video*). These attributes are primarily managed using the *sleap-io* (v0.0.11) package.

The load class method facilitates the loading of predictions for a given series. It constructs paths to the prediction files based on provided root names and uses the *sio.load_slp* function to load these predictions. Additionally, the video representation of the image series is loaded using *sio.Video.from_filename*.

The class also provides utility methods and properties, such as *series_name*, which extracts the name of the series from the HDF5 filename using the *pathlib* library. The *len* method returns the number of images in the series, while the *getitem* and *iter* methods enable indexing and iteration over the series, respectively.

The *get_frame* method returns labeled frames for both primary and lateral predictions for a specified frame index. This is achieved by finding the corresponding labeled frames using the find method from the *sio.Labels* class.

Visualization capabilities are provided by the *Series.plot* method, which overlays predictions on the image. This method utilizes the *matplotlib* (v3.8.0) and *seaborn* (v0.13.0) libraries for plotting. The *get_primary_points* and *get_lateral_points* methods retrieve the primary and lateral root points, respectively, for a given frame index. These methods return the root points as numpy arrays, with shapes indicating the number of instances and nodes.

#### Angle-Related Traits

The *sleap_roots.angle* module is designed to compute angles between vectors that represent root structures and the gravity vector **(Fig 4A, Table 4A)**. To represent a root, a vector is constructed using the base node and either the distal or proximal node. The distal node is identified as the first non-NaN node in the initial half of the root, while the proximal node is the last non-NaN node in the latter half of the root. The computation of the angle between these vectors is achieved using the *numpy.arctan2* function from the *numpy* library (v1.26.0).

**Fig. 4.**
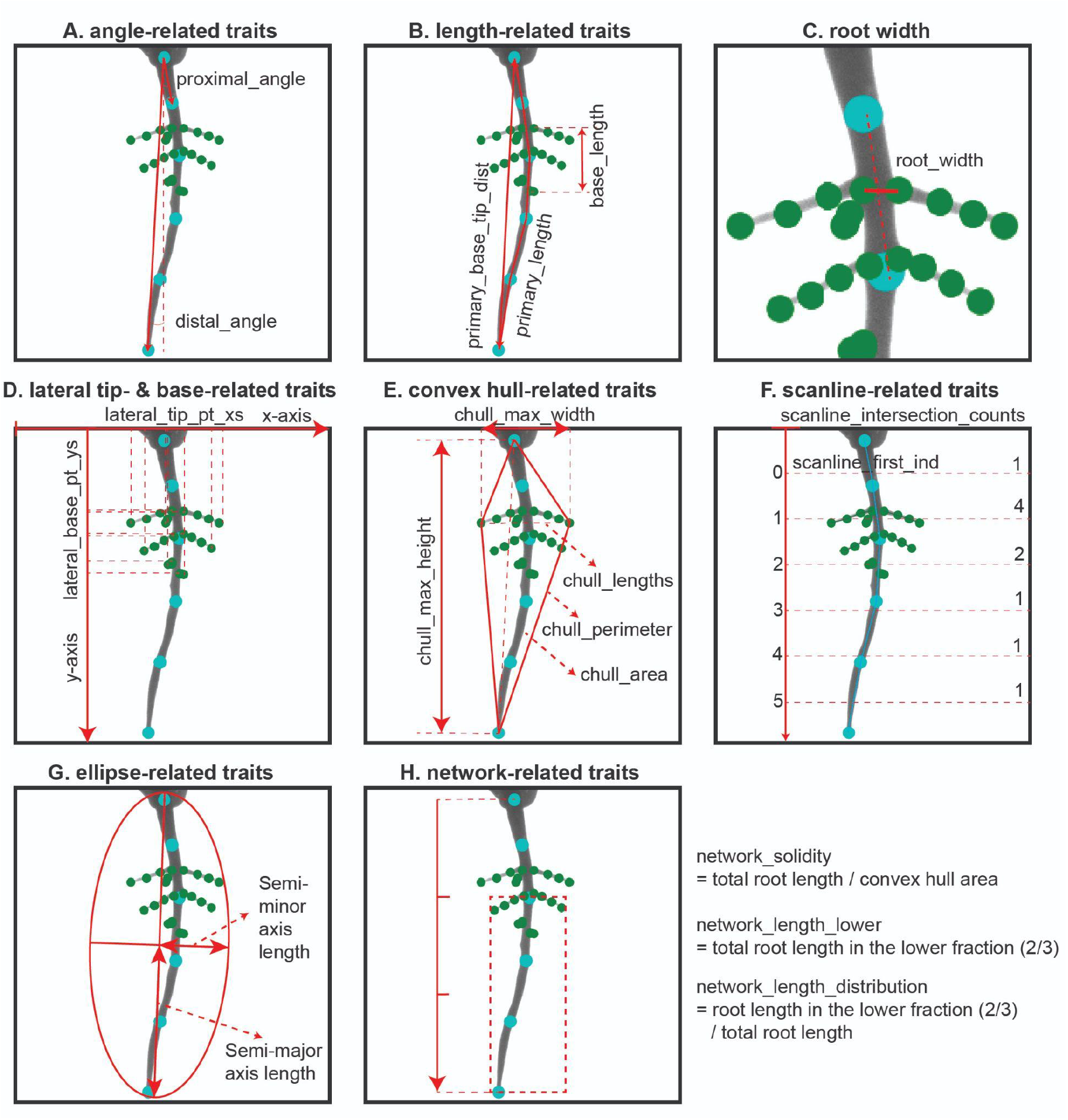
Schematics of root trait computation methods from landmark data. The landmarks in cyan are primary root predicted landmarks, while green are lateral root landmarks.

**Table 4.**
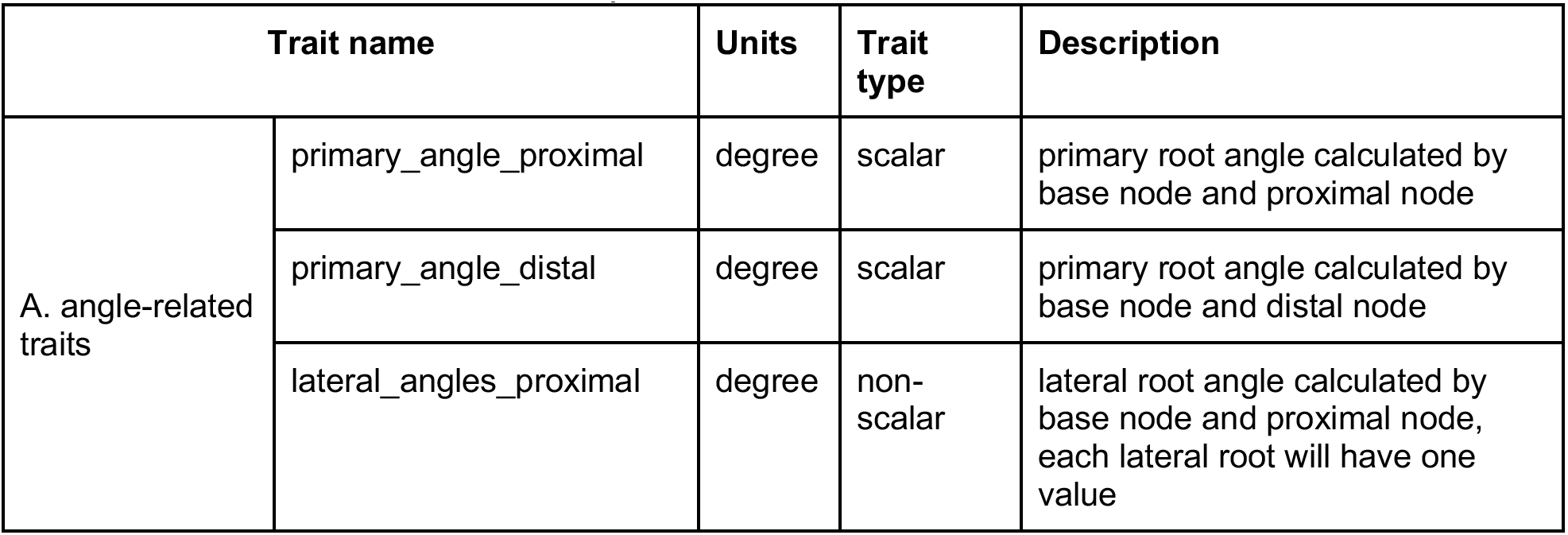
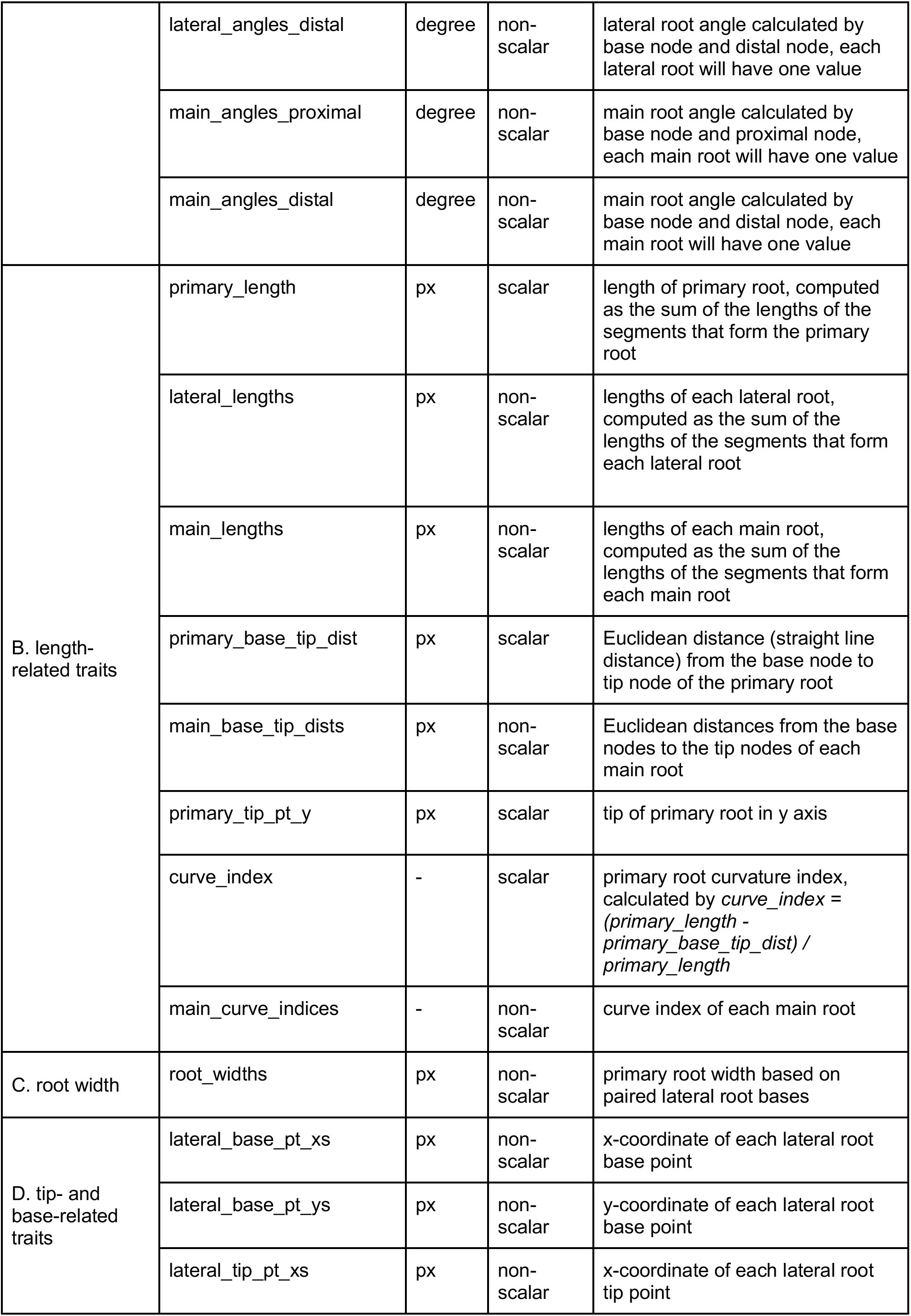
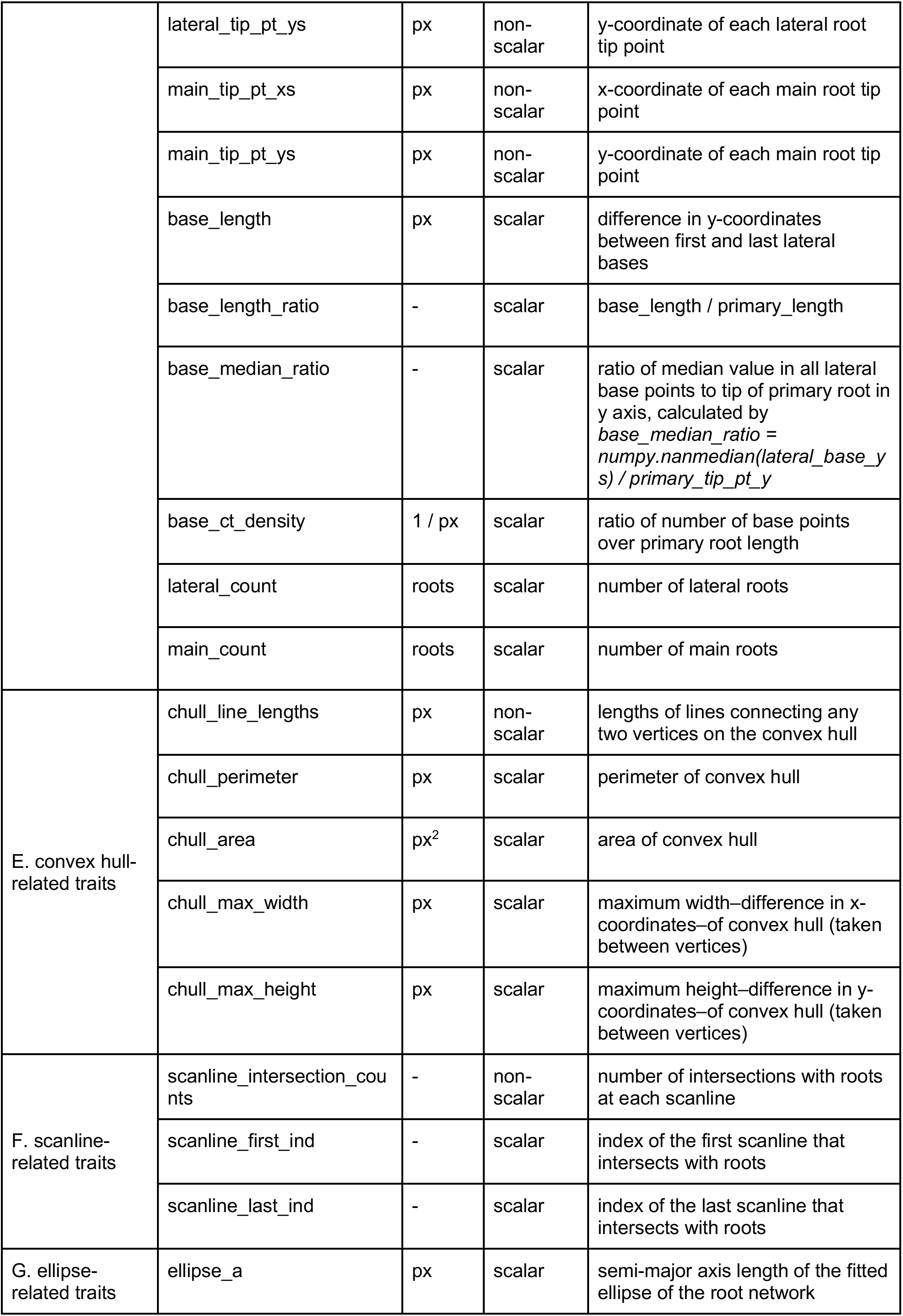
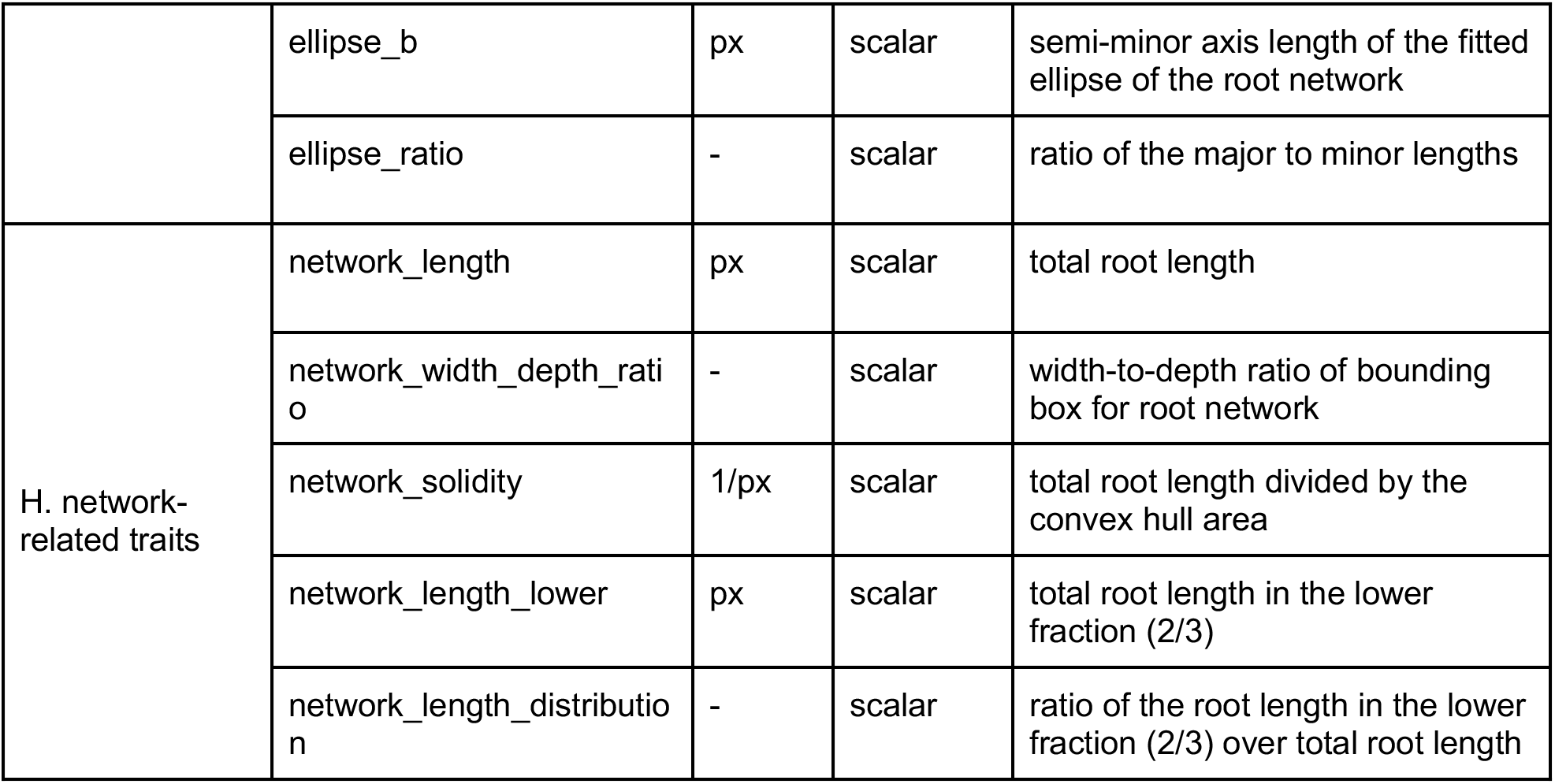
Pose-derived traits description.

#### Length-Related Traits

In the *sleap_roots.lengths* module, root lengths are computed by summing the segment lengths (Euclidean distance) between nodes for each individual root **(Fig 4B, Table 4B)**. This calculation utilizes the *numpy.diff* and *numpy.linalg.norm* functions from the *numpy* library (version 1.26.0). The total length for each root is then aggregated using *numpy.nansum*.

#### Tip-and-Base-related Traits

In the *sleap_roots.tip* module, tips of roots are identified as the terminal nodes using *numpy* array operations (v1.26.0). Conversely, bases are recognized as the initial nodes of each root. Utilizing these landmarks, various traits can be derived, including the density of base points on the primary root, denoted as *base_ct_density* **(Fig 4D, Table 4D)**.

For dicot plants, the bases of lateral roots are employed to approximate root widths **(Fig 4C, Table 4C)**. Initially, lateral root bases on both the right and left flanks of the primary root are pinpointed. Subsequently, a *LineString* object representing the primary root is constructed using the *LineString* function from *shapely.geometry* (*shapely* v2.0.1). To locate points on the primary root proximate to the bases of the lateral roots, we employ *nearest_points* from *shapely.ops* and *Point* from *shapely.geometry*. Each point’s normalized projection, relative to the length of the primary root’s *LineString*, is then determined for every base on both the left and right sides. A cost matrix is formulated based on the discrepancies in projections between the bases on the two sides. Utilizing the Hungarian matching algorithm, an optimal pairing between the left and right bases is established, aiming to minimize the total difference in projections. This is achieved using *linear_sum_assignment* from *scipy.optimize* (*scipy* v1.11.3). Post the application of a user-defined tolerance for projection differences between base pairs (default set at 0.02) and the exclusion of pairs not intersecting the primary root line, the distances between the matched base pairs are computed using *numpy.linalg.norm*.

#### Convex-hull-related Traits

Within the *sleap_roots.convhull* module, the convex hull is determined using the ConvexHull function from *scipy.spatial* (scipy v1.11.3). Traits derived from the convex hull include the convex hull perimeter, area, width, height, and distances between vertices (Fig 4E, Table 4E).

#### Scanline-related Traits

Within the *sleap_roots.scanline* module, functions employ horizontal “scanlines” to quantify the number of roots based on their intersections with these lines. This procedure is executed using *numpy* array operations (v1.26.0) on the list of points that delineate the root network (**Fig 4F**, **Table 4F**).

#### Ellipse-related Traits

Within the *sleap_roots.ellipse* module, the *fit_ellipse* function is designed to approximate an ellipse to the points characterizing the root network. This function yields the semi-major and semi-minor axes of the fitted ellipse, in addition to the ratio of the major to minor lengths **(Fig 4G, Table 4G)**. This is achieved using the *EllipseModel* from *skimage.measure* (*scikit-image* v0.22.0).

#### Network-related Traits

In the *sleap_roots.networklength* module, a suite of functions is dedicated to the analysis of the root network. The *get_network_distribution* function leverages the *Polygon* class from the *shapely* library (v2.0.1) to transform the lower fraction of the root network’s bounding box into a polygon. For each root, *LineString*s are formed, and the intersection and length properties of shapely’s *LineString* are utilized to compute the length of roots that intersect with this polygon. The total *network_length* is ascertained by aggregating the lengths of all roots in the network using *numpy.nansum*, with individual root lengths being pre-calculated via the *get_root_lengths* function. The remainder of the computations in the *sleap_roots.networklength* module predominantly employs *numpy* array operations **(Fig 4H, Table 4H)**.

#### Trait Graphs

In the *sleap_roots.trait_pipelines* module, trait maps are implemented for different types of plants. The module employs a directed graph to represent the hierarchical dependencies between root traits. The graph starts at the points from the SLEAP predictions and ends with root traits per plant. Classes are defined for each unique trait map which determine the initial points fed into the graph and the connections between nodes. The graph representation offers flexibility, making it easy to adapt the pipeline for different input predictions and relationships between traits. This adaptability ensures that the pipeline remains versatile and can cater to the specific needs of various types of plants with different trait relationships.

The graph was built using *networkx.DiGraph* from the *networkx* package (v3.1). To ensure that traits are computed in the correct sequence, respecting their dependencies, the computation order was defined using *networkx.topological_sort*. By leveraging topological ordering, the module can systematically compute each trait only after its dependent traits have been calculated, ensuring efficiency in the trait extraction process.

#### 2.7.1 Dicot Pipeline

An advanced root trait extraction pipeline was designed to process SLEAP predictions and subsequently generate root traits for dicot plants in CSV format. This pipeline is tailored for plants with primary and lateral root predictions.

From each frame, 35 distinct traits were derived, culminating in a total of 1035 traits for each plant. A visual representation of these 35 traits was constructed using a mermaid graph **(Fig. S1)**, originating from the SLEAP landmarks, where individual nodes symbolize traits and arrows between nodes show the actions of the functions in the sleap-roots code repository as defined in the trait map for the dicot pipeline.

Of the 35 root morphological traits computed, 25 are scalar traits 10 are non-scalar traits. For these 10 non-scalar traits, a suite of nine summary statistics— maximum, minimum, mean, median, standard deviation, and percentiles at 5th, 25th, 75th, and 95th—were applied. This led to the derivation of 90 summarized non-scalar traits per frame. By applying summary statistics to each frame trait, a comprehensive set of 1035 summarized traits per plant was achieved.

#### 2.7.2 Younger Monocot Pipeline

Similarly, a pipeline tailored to the younger monocot predictions, consisting of primary and seminal roots–referred to as “main” root traits for brevity–was constructed beginning at the predictions and ending at traits in CSV format. For each frame 30 distinct traits are calculated. The mermaid graph depicting the trait for younger monocots is shown in **Fig. S2**. In this pipeline, there are 21 scalar and 8 non-scalar traits per frame. The summary statistics— maximum, minimum, mean, median, standard deviation, and percentiles at 5th, 25th, 75th, and 95th—were applied to the non-scalar traits per frame and again over the 72 images in the series, resulting in 918 traits per plant.

### 2.8 Manually measured traits using FIJI

Given the labor-intensive nature and time demands of manual trait measurements, we opted to focus our error evaluation using manual trait measurements on a select number of traits for rice, a monocot, and soybean, a dicot. These traits were chosen because they are representative descriptors of the root system architecture and are more straightforward to measure manually, minimizing potential measurement errors. We used FIJI’s multi-point tool for point measurement and the segmented line tool for root lengths [34]. For rice, we evaluated the y-coordinate of the deepest root tip, where roots could be either primary or seminal roots, and the primary root length, which correspond to the traits in **Table 4** of *main_tip_pt_ys_max_median* and *primary_length_median* from the younger monocot pipeline. For soybean, we assessed the y-coordinates of the first lateral base and the deepest lateral tip, corresponding to the traits of *lateral_base_ys_min_median* and *lateral_tip_pt_ys_max_median* from the dicot pipeline.

### 2.9 Comparison with RootPainter

The study aimed to assess our computer vision approach for plant phenotyping in comparison with the established RootPainter and Rhizovision method. Both RootPainter and SLEAP were used to train minimal models, and root traits derived from each method were compared with the ground truth. Time metrics for every phase were documented. The goal was to label sufficiently in both software to yield acceptable predictions – sufficiently discerning between root and non-root regions–and compare the time it took to create a “minimally accurate model” in each. We used 3 DAG rice images, labeling both seminal and primary roots, to train a model on each platform.

Our workstation ran on Windows 10 Pro, powered by an Intel(R) Core(TM) i7-9700K CPU (3.60GHz, 8 cores, 8 logical processors), paired with an NVIDIA GeForce RTX 2070 Super boasting 8 GB of VRAM, operating with driver version 531.79 and CUDA version 12.1.

For the SLEAP benchmark, we made a project for the primary and seminal roots of 3 DAG rice samples. From a randomly chosen subset of 10 plants, 2 frames per plant were labeled (20 frames in total), noting the labeling duration for each. Adhering to the labeling rules defined for the “Rice seminal roots (3 DAG)” model, the first and 30th frames from each plant were labeled. Following this, the model underwent training based on the configuration detailed elsewhere in this article. Testing was conducted on a distinct set of 75 user-labeled frames. The resultant predictions met acceptable standards. The derived root traits from SLEAP’s model and the labeled frames were compared, and errors were assessed as the absolute difference between the two sets. For the RootPainter benchmark, the aforementioned 10 rice plants chosen for SLEAP were reintroduced to RootPainter (version 0.2.25; [16]). The images were resized to half, matching input scaling used in SLEAP, and cropped to isolate the cylinders, resulting in a final 512 px X 512 px dimension. One tile per image was used. The initial labeling spanned 38 min 46 s. The instructions from the RootPainter github page [35] guided the labeling process, maintaining a consistent pace. Training was initiated after completing 2 labeled images. Upon evaluating the predictions after six labels, they were still essentially random. Extended training was permitted until corrective predictions seemed viable.

Notably, enhanced accuracy required more extensive labeling, and corrective annotations markedly refined the model. In total, 14 frames were labeled, achieving desirable predictions. The best model was used to segment the same 75 frames tested with SLEAP (these frames are also distinct from the training set used in RootPainter). The derived segmentations were directed to Rhizovision for root trait extraction, employing specific settings: whole-root mode, retention of the largest component, image threshold set at 200, and root pruning active at a threshold of 5. Root traits derived from Rhizovision were benchmarked against the ground truth for error computation.

### 2.10 Statistical analysis

We use the localization error to evaluate the accuracy of the models (**Fig. 5**, **Fig. 8**). The localization error is the distance of the predicted landmark from the ground truth landmark–the landmark labeled by a human. For each model, we created three randomly selected test sets of plants we had ground-truth labels for. The test set is approximately 10% of the total labeled data for each model (**Table S1, S6**). We trained a model with each training set and evaluated the model on the plant-wise test set hidden from the model during the training.

**Fig. 5.**
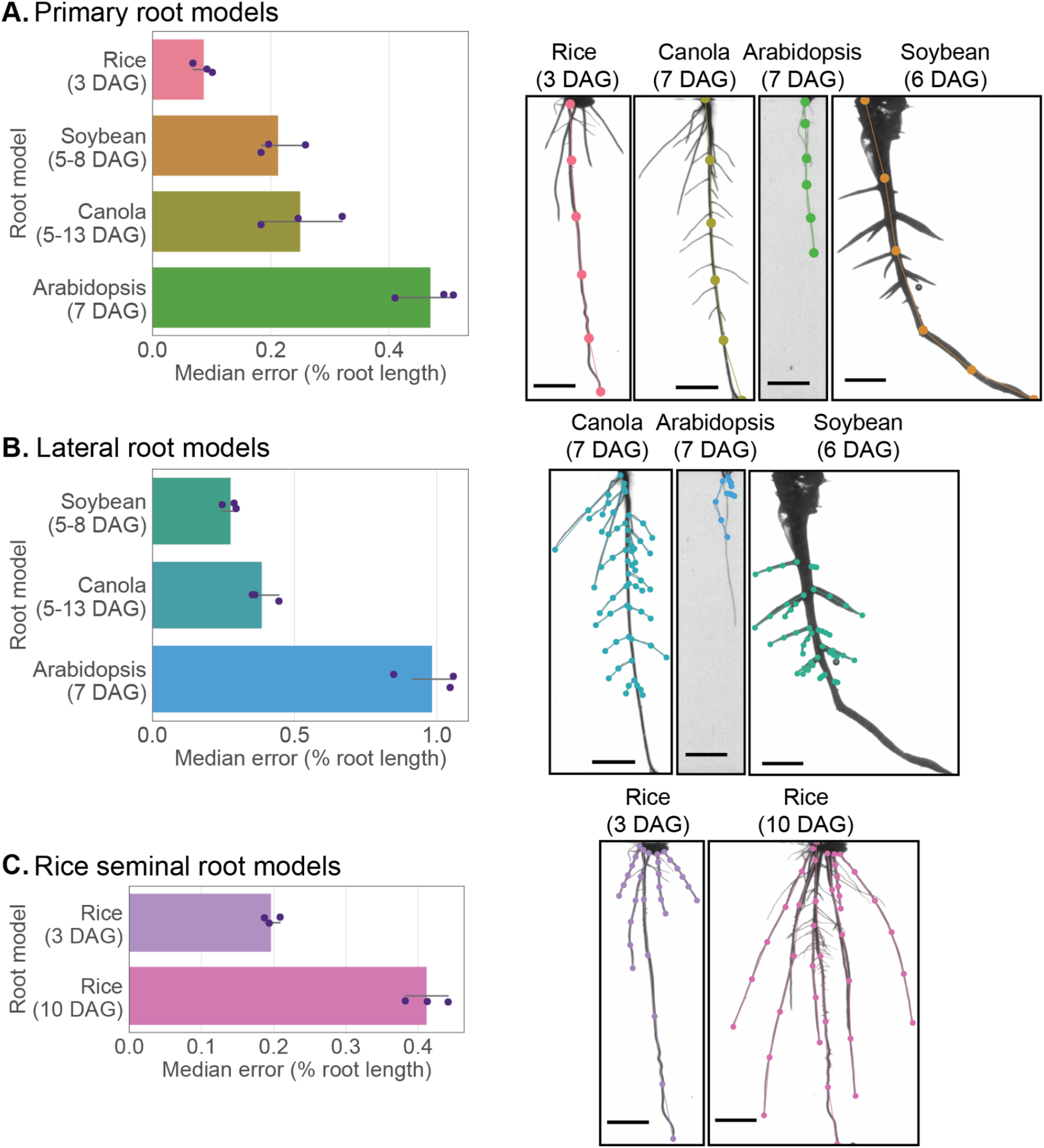
SLEAP models accurately located root landmarks in 4 species and 3 classes of roots. (A) Primary root, (B) lateral root, and (C) seminal root accuracies and predictions are displayed. The accuracy is calculated as the median localization error for a randomly selected held-out test set normalized by the average root length for that dataset. The bar graph presents the models ordered by mean accuracy, with error bars showing the 95% confidence interval (n=3). The scale bar is 1 cm.

The SLEAP-based root traits error was evaluated by fitting a regression line using predicted traits value and the corresponding manually measured values using Fiji (**Fig. 6**). The coefficients of determination (R^2^) and the coefficient of regression lines were calculated. Two rice traits (deepest root tip depth and maximum primary root length) and two soybean traits (deepest lateral root tip depth and top lateral root base depth) were used to evaluate the SLEAP-based traits error.

**Fig. 6.**
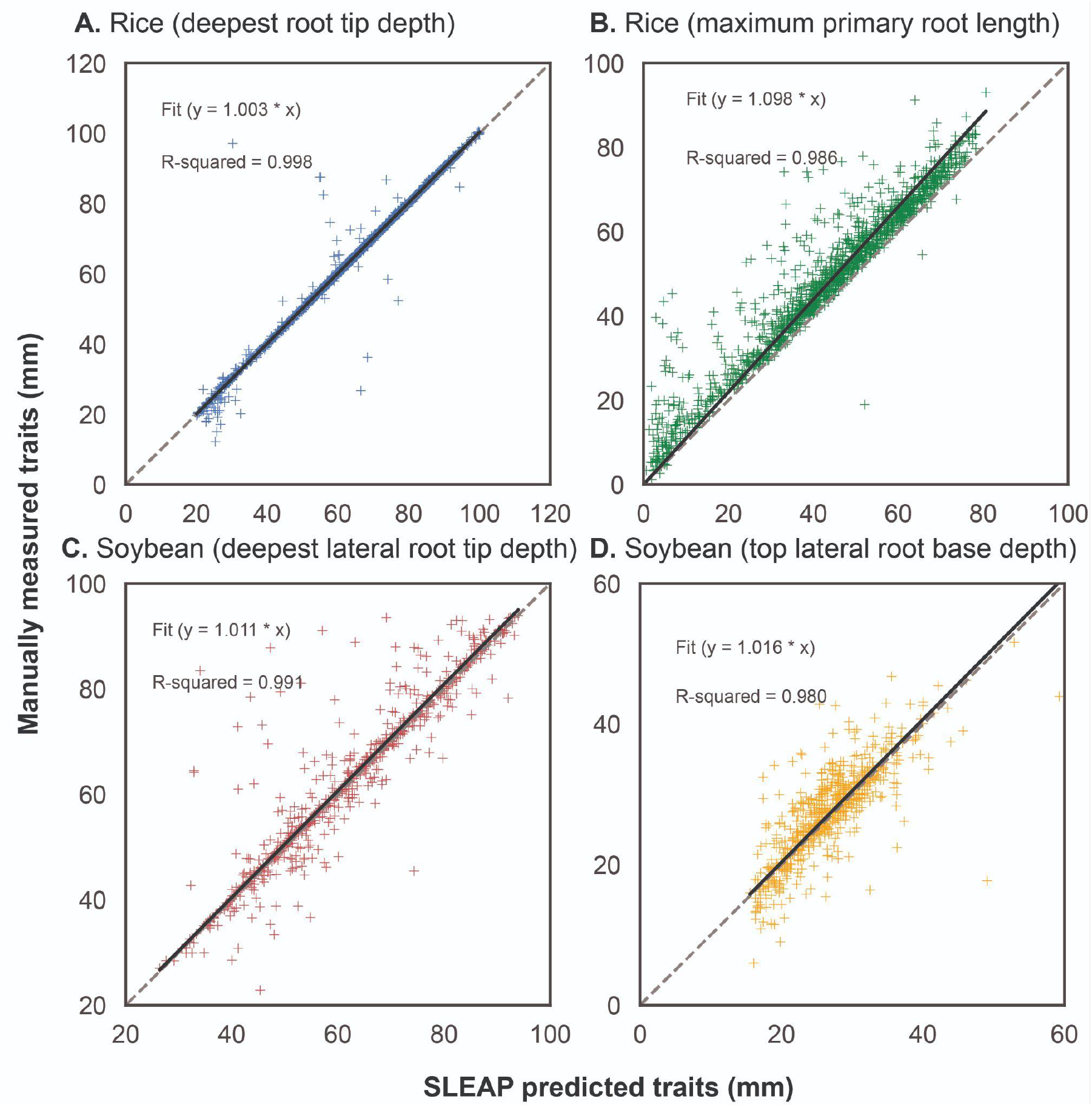
Comparison of root traits from SLEAP with manually measured traits. Scatter plots of trait errors for (A, B) rice (3 DAG) and (C, D) soybean datasets. For the rice dataset with 1301 plants, manually measured and SLEAP predicted traits (deepest tip of all roots in y coordination and maximum primary root length) were compared. For the soybean dataset with 600 plants, manually measured and SLEAP predicted traits (y-coordinate of deepest lateral root tip and lateral origin) were compared. The gray dashed lines represent the 1:1 lines (y=x), while the black solid lines depict the regression lines, with the associated regression functions and R² values displayed in each panel..

Proofreading was conducted with the SLEAP predicted landmarks. The absolute difference of z-scores were used to compare each trait of each plant before and after proofreading (**Fig. 7**). The z-score for each trait and plant was calculated based on the formula *z* = (*x* – *μ*)/*σ*, where x is the trait value, *μ* is the average trait value for proofread plants, *σ* is the standard deviation of trait value for proofread plants. In addition, Pearson correlation was calculated for each trait using proofread and non-proofread data (**Tables S3-5**).

**Fig. 7.**
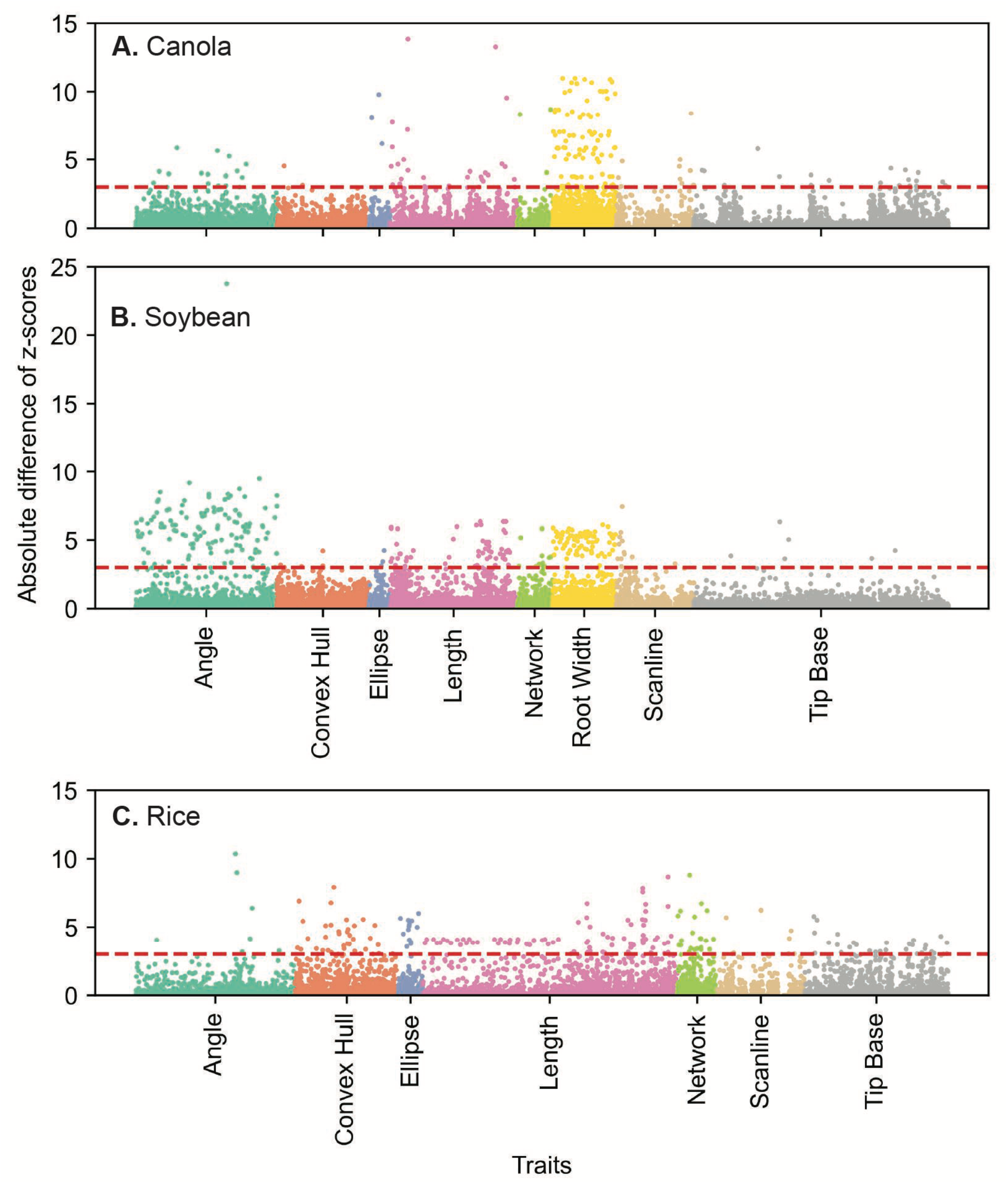
Comparison of proofread and non-proofread traits for each crop. Manhattan plots of the normalized difference between traits derived from proofread and non-proofread data for (A) canola (7 DAG), (B) soybean (6 DAG) and (C) rice (3 DAG). The x-axis of each figure indicates unique traits grouped by category. The y-axis of each figure represents the absolute difference between z-scored proofread and non-proofread trait values. Points correspond to individual trait values for different plants. The red dashed line represents three standard deviations.

**Fig. 8.**
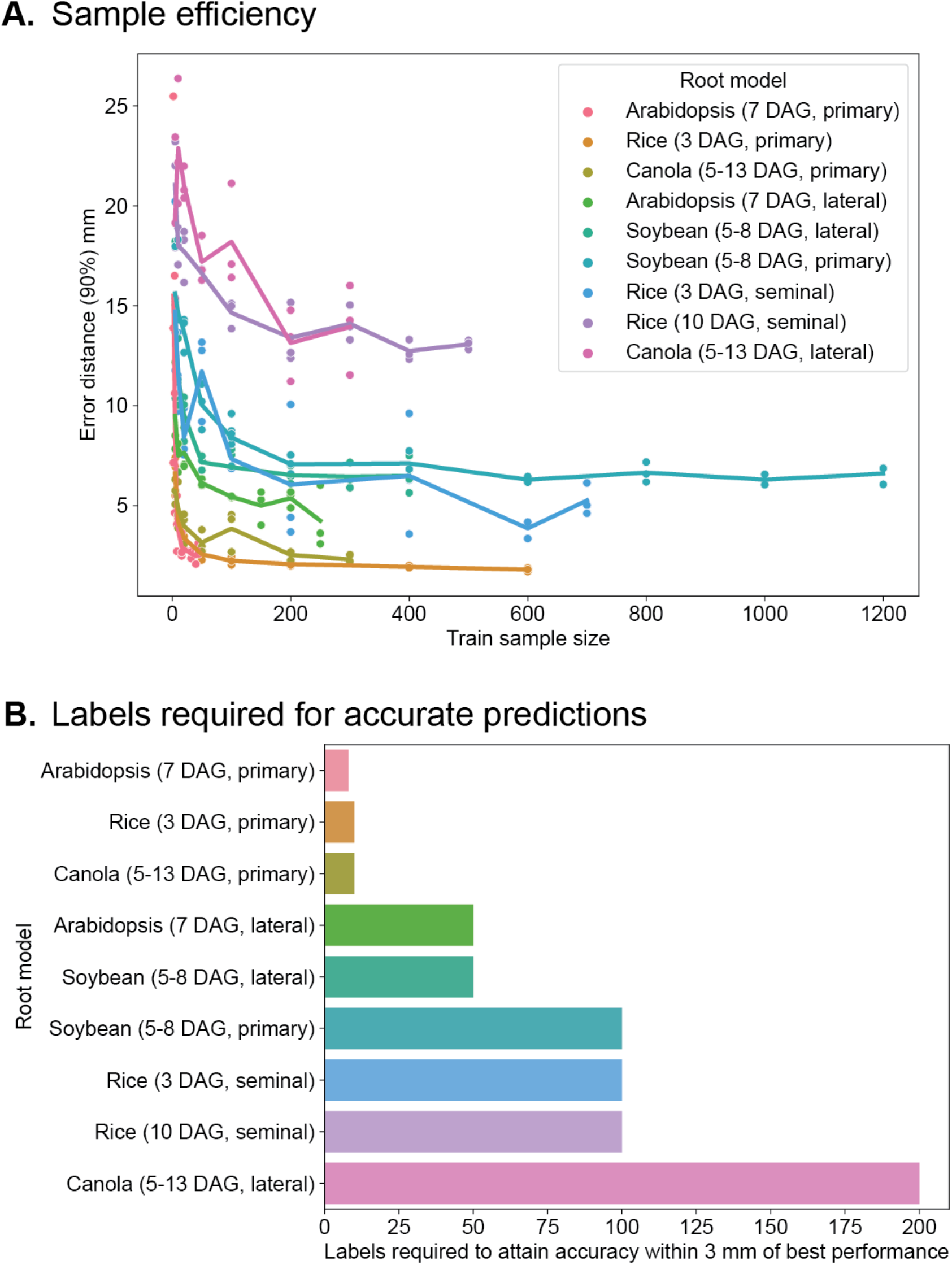
Efficient models require fewer labels for accurate predictions and facilitate diverse training sets. (A) Models with the number of labeled frames given by the train sample size were trained for each root model. The y-axis shows the localization error at the 90 percentile so that error-prone examples are included in this error estimation (we use the high-end of the error distribution). The line plot connects the average error distance at each sample size (n=3). The sample efficiency curves plateau as the increase in training sample size does not further reduce the localization error. (B) We can estimate the number of labeled frames needed for accurate predictions by asserting a threshold of 3 mm as the change in localization error with a change in the number of labeled frames in the training set. For most models, we are within 3 mm of the best model within 100 labeled frames.

**Fig. 9.**
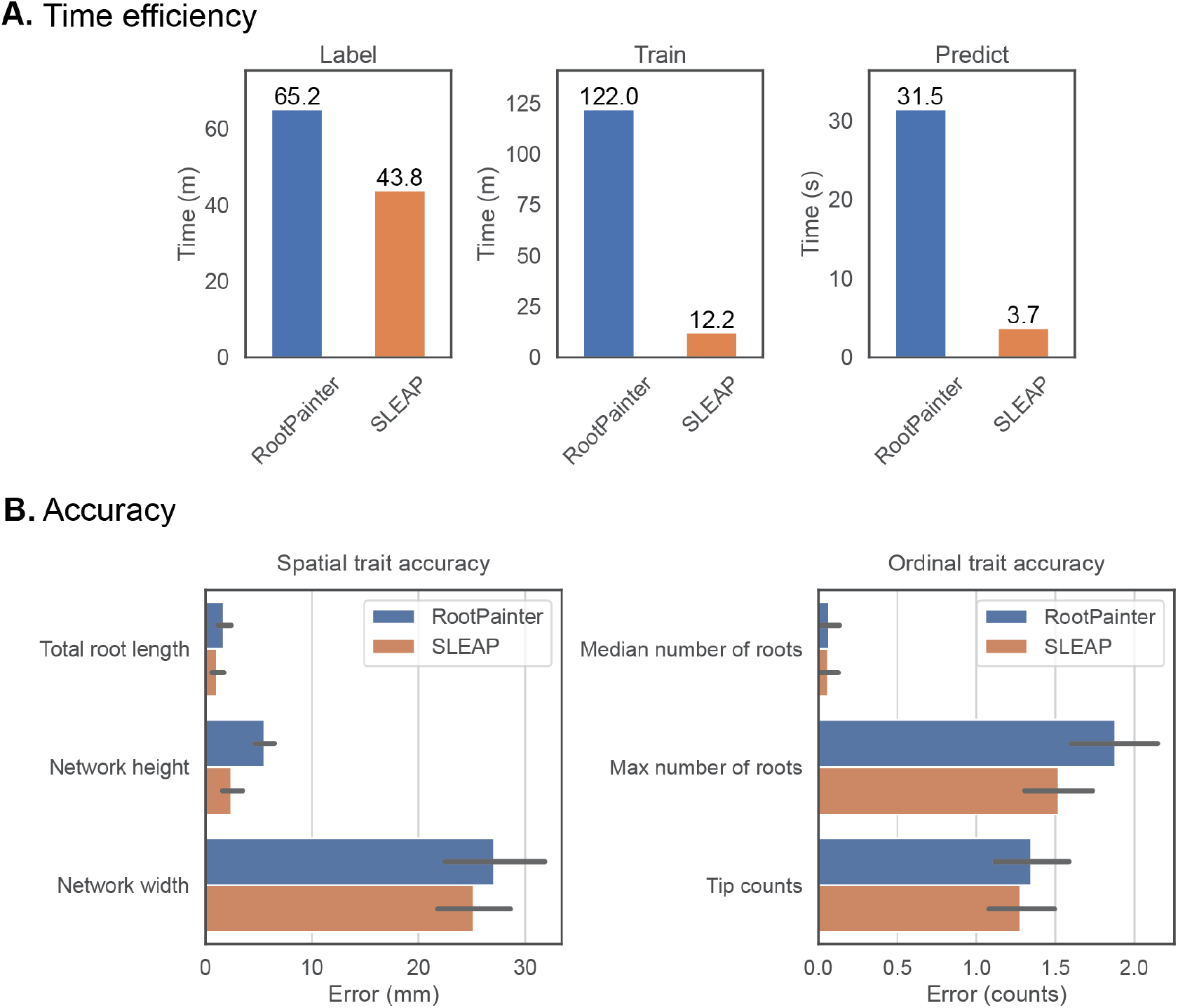
Root traits are extracted in a fraction of the time with the same or better accuracy as segmentation-based methods. (A) Minimal models for accurate root predictions are labeled and trained in both RootPainter and SLEAP and the time is recorded. 75 root images are predicted using these trained models. (B) The accuracy of the predictions from this minimally trained model is evaluated for the common set of traits between Rhizovision and SLEAP. The height of the bar graphs is the mean error, with error bars showing the 95% confidence interval (n=75).

SLEAP-based plant traits were used to do plant accession classification using random forest classifiers (**Fig. 10**). We selected 70 plants with 10 accessions and more than six replications per accession. The plant traits were derived based on proofread SLEAP predictions. Five-fold cross-validation overall accuracy was used to evaluate random forest performance. Four random forest classifier parameters were fine-tuned with a randomized search algorithm. The parameters were number of trees, maximum depth of the tree, minimum samples per split, and minimum samples per leaf. The best model was selected based on the highest accuracy. The parameter settings of the best classifier are *max_depth* of 10, *min_samples_leaf* of 1, *min_samples_split* of 4, and *n_estimators* of 712.

**Fig. 10.**
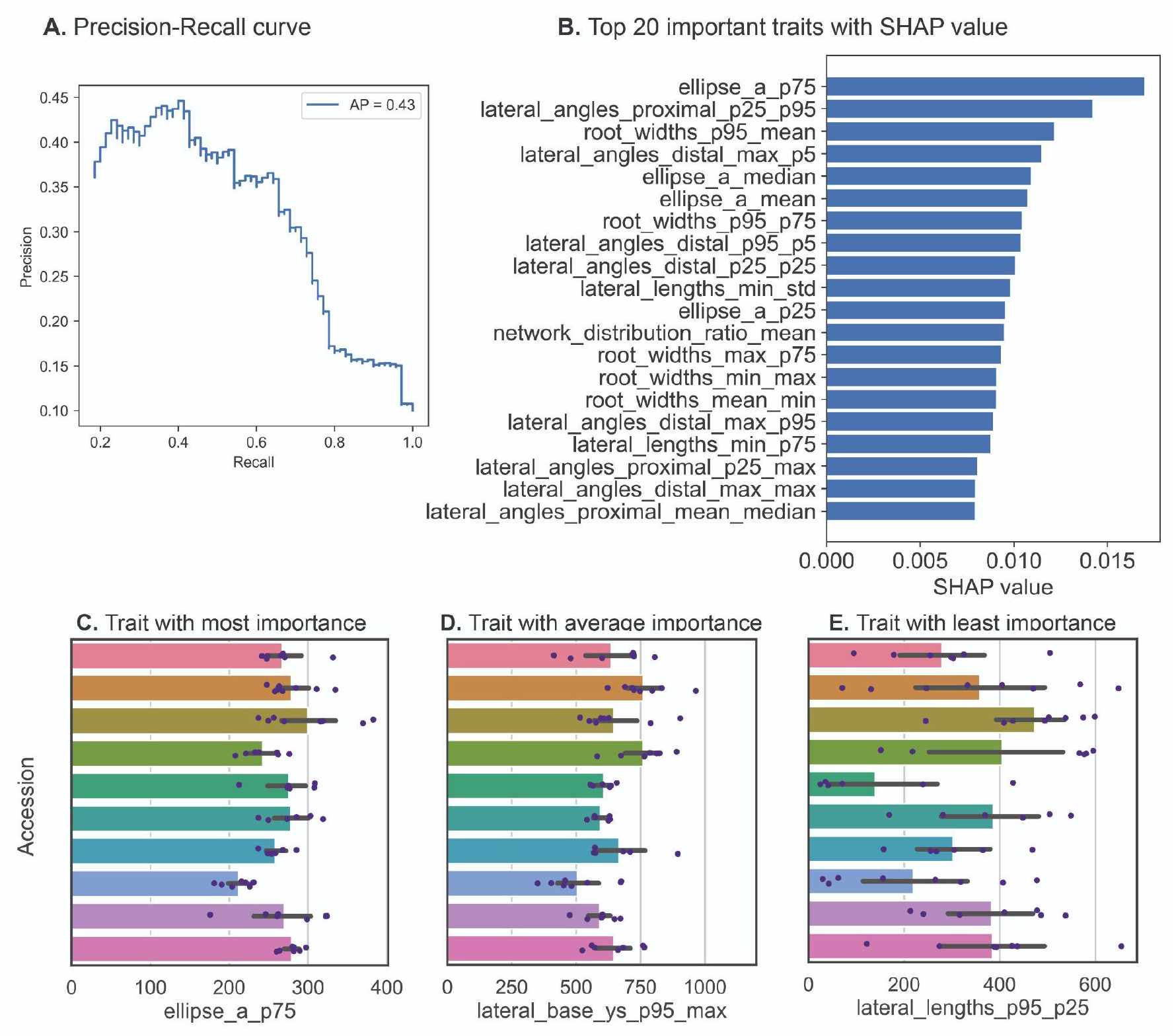
Genotype prediction using phenotypic traits. (A) Classification performance using precision-recall curve of accession classification of 70 diverse soybean plants. AP stands for average precision value. (B) The top 20 most important traits for accession classification using fine tuned random forest best parameters. (C-E) Three traits showing varying amounts of importance in genotype differentiation of samples. The x-axis is the trait value, while the y-axis represents the accession.

The SHapley Additive exPlanations (SHAP) algorithm was used to evaluate the accession classification trait importance [27]. The input traits were the same as the accession classification dataset, while the target is the plant accession as well. SHAP determined the importance value for each trait. A higher SHAP value of one trait means this trait plays a more important role in the classification of accessions. A precision-recall curve was conducted to show the tradeoff between precision and recall. The average precision value was used based on the precision values of ten accessions with macro metrics. A higher area under the curve, a higher precision and recall value.

To confirm the reliability of root traits derived from sleap-roots, we conducted exploratory data analysis and visualized phenotypic trait spaces (**Fig. 11**). Our *DicotPipeline* aggregated 1035 root traits across 2082 soybean samples. Only samples without any missing (NaN) values across all traits were retained, resulting in a dataset of 2009 samples. We standardized the data using z-score normalization followed by principal component analysis (PCA) employing the *StandardScaler* and *PCA* modules from *scikit-learn* (v1.0), respectively. To robustly estimate the covariance of the dataset, a Minimum Covariance Determinant (MCD) [36] estimator was fitted using scikit-learn’s *MinCovDet* on 75% of the data variability represented by the first eight principal components. For dataset quality assurance, we adopted the Mahalanobis distance as a metric to flag potential outliers, leading to manual inspection of all samples with a distance value of 10 or above. This quality control step resulted in the retention of 1877 samples. Subsequent PCA was performed post-outlier exclusion to identify the traits with the highest variance explanation, which were then employed to annotate UMAP visualizations of the phenotypic trait space. The UMAP embeddings were generated using default settings on the z-score normalized dataset.

**Fig. 11.**
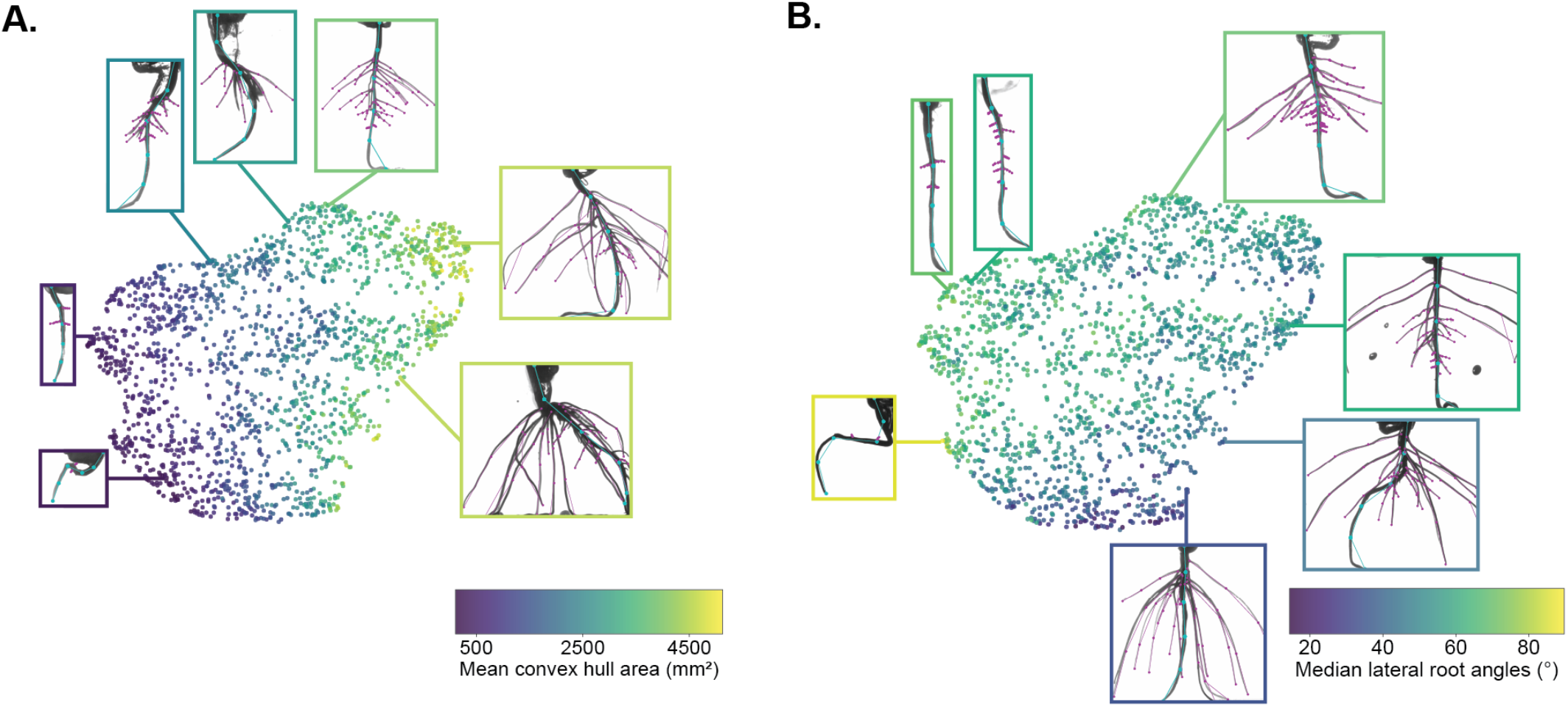
Visualization of phenotypic space through large-scale root trait mapping. (A) The UMAP, color-coded according to the mean convex hull perimeter, illustrates the significant role this trait plays in shaping the trait space. (B) A similar color-coding strategy is used to highlight the median lateral root angle’s contribution to the global trait structure. These examples, each uniquely colored based on their corresponding trait, offer a comprehensive view of the phenotypic diversity observed in the screen.

## 3. Results

### 3.1 Pose estimation robustly localizes root system landmarks across species

We find that all of our models have median localization error of less than 1% of the root length (**Fig. 5**, S2). The most accurate model relative to its root length was the rice primary root 3 DAG model at 0.087% root length (0.540 mm) followed by the rice seminal root 3 DAG model at 0.197% root length (0.662 mm). The model with the highest error relative to its root length was the Arabidopsis lateral 7 DAG model at 0.984% root length (0.843 mm) followed by the Arabidopsis primary 7 DAG model at 0.470% root length (1.537 mm) and the rice seminal 10 DAG model at 0.412% root length (2.281 mm).

### 3.2 Phenotypic traits are accurately extracted from root system landmarks

Two rice traits from the younger monocot pipeline were compared with manually-measured traits (**Fig. 6A, B**). The R^2^ values were 0.998 and 0.986, while the coefficient of regression lines were 1.003 and 1.098 for the deepest root tip depth (*main_tip_ys_max_median*) and maximum primary root length (*primary_length_median*), respectively. Two soybean traits from the dicot pipeline were selected to evaluate the SLEAP traits by comparing with manually-measured traits as well (**Fig. 6C, D**). The R^2^ values were 0.991 and 0.980, while the coefficient of regression lines were 1.011 and 1.016 for the deepest lateral root tip depth (*lateral_tip_pt_ys_max_median*) and top lateral root base depth (*lateral_base_pt_ys_min_median*), respectively. In both rice and soybean, the traits derived from the younger monocot and dicot pipelines respectively, demonstrated a high degree of correlation with manually-measured traits, as indicated by the high R² values and coefficients of regression lines close to 1, underscoring the reliability and accuracy of these pipelines in phenotypic trait assessment.

As it was not feasible to extract all traits for all plants via manual annotation, we next compare the accuracy of traits derived from fully automated predictions to those that were manually proofread (**Fig. 7**). Manual proofreading is drastically faster than manual annotation, but trait results may differ since proofreading corrects predictions while manual measurements are not informed by predictions. We estimate trait computation accuracy as the absolute difference of z-scores between proofread and non-proofread predictions for each trait and plant. The z-scores were calculated using the estimators of the distributions of the proofread traits, so that the difference in z-scores between proofread and non-proofread traits is how many standard deviations of the proofread trait the non-proofread data differed from the proofread trait.

For canola (**Fig. 7A**), we find that the most inaccurate traits were the *lateral_lengths_max_std* (Δz = 0.576), *lateral_lengths_std_max* (Δz = 0.570), and *lateral_lengths_max_max* (Δz = 0.516); for soybean (**Fig. 7B**): *base_ct_density_p95* (Δz = 0.542), *primary_angle_distal_min* (Δz = 0.472), and *base_length_ratio_p95* (Δz = 0.419); and for rice (**Fig. 7C**): *primary_length_std* (Δz = 0.300), *primary_base_tip_dist_std* (Δz = 0.229), and *primary_tip_pt_y_std* (Δz = 0.226). For the canola dataset, we have in total 110,745 data points (107 plants * 1035 traits), of which 96.2% data points are within one standard deviation; for soybean (n = 70 plants * 1035 traits = 72,750 data points), 96.3% are within one standard deviation. For the rice dataset (n = 270 plants * 918 traits = 247,860 data points), 99.5% data points are within one standard deviation (fewer traits are computed for monocots). The average correlation values of proofread and non-proofread traits were 0.958, 0.927, and 0.982 for canola, soybean, and rice, respectively (**Tables S3-5**). In our analysis across canola, soybean, and rice datasets, specific traits exhibited higher inaccuracy, as evidenced by the Δz values. However, despite these discrepancies, the vast majority of data points fall within one standard deviation, underlining the overall precision of the datasets. Furthermore, average correlation values between proofread and non-proofread traits remained robust across all three crops, suggesting consistent reliability in our trait measurements.

### 3.3. Landmark-based methods are faster and more efficient than segmentation-based approaches

A key advantage of pose estimation is that it requires less laborious annotation than segmentation since only the landmark points need to be annotated by the user. In SLEAP, this is further sped up by human-in-the-loop annotation, in which models are iteratively trained, used to generate predictions, and those predictions imported to the annotation GUI for proofreading.

While annotation speed is a key factor for generating reliable models, it is not clear how many annotations are necessary to achieve high accuracy. To determine this, we conducted a series of sample efficiency experiments, in which subsamples of each dataset were used to train SLEAP models (**Fig. 8**). In agreement with previous work using SLEAP in plants [21], we observe diminishing improvements in accuracy beyond tens to hundreds of labels (**Fig. 8A**). Relative to the peak accuracy, most datasets require <100 labeled images, with the exception being canola lateral root models that achieve peak accuracy at 200 labels (**Fig. 8B**). Canola is the exception, due to the large amount of diversity of the training set; these plants have the widest range in age, from 5-13 DAG, and were randomly selected from a very large diversity screen spanning over many months.

Next, we compare the speed and accuracy of our approach to a segmentation-based approach. RootPainter [16] is a commonly used, user-friendly segmentation tool that employs deep learning and human-in-the-loop annotation workflow which continuously trains and generates predictions during the annotation process. We find that we can train a model in SLEAP using relatively few labeled frames and the time to label, train and predict using landmark detection is significantly lower than RootPainter. We see a 32.8% decrease in labeling time when using SLEAP to annotate landmarks (**Fig. 9A**). Training and inference time was significantly decreased in SLEAP: there was a 90.0% decrease in training time and 88.1% decrease in inference time (**Fig. 9A**). When we compare the root traits extracted from the two pipelines we find that SLEAP has the same or better accuracy in our dataset (**Fig. 9B**).

Root trait extraction times using the younger monocot pipeline with all 918 traits took an average of 0.44 seconds per plant (n=1153). The dicot pipeline with all 1035 traits took an average of 0.70 seconds per plant (n=2082).

### 3.4. Pose estimation-based traits can be used for downstream analyses

As a proof-of-principle that our pipeline can be used in common downstream analyses of plant RSA phenotypes, we used the traits we extracted in both supervised (genotype classification) and unsupervised (UMAP visualization) tasks.

For classification, we used traits from our pipeline to predict the genotype of soybean plants. 70 plants were sampled from ten accessions to perform supervised classification. Each accession had at least six replicates. The best random forest classifier achieved an overall accuracy of 0.400 using 5-fold cross validation after the parameter finetuning. A precision-recall curve was generated based on the accession classification results (**Fig. 10A**) with an average precision of 0.43. We then used SHAP [27] to evaluate the importance of each trait with the best random forest classifier. The top 20 most important traits are shown in **Fig. 10B**, including the lateral angle-, root width- and ellipse-based traits. The least important traits were scanline count-, lateral base-, and lateral tip-based traits. Three typical traits with the most, average, and least importance for accession classification are visualized in **Fig. 10C-E**. These analyses demonstrate that traits derived from our pipeline can be used to train genotype classifiers with sufficient accuracy and explainable predictions.

Next, we used the predicted traits for soybean across 1,877 plants to map out the phenotypic space of these plants, a common task in high throughput screening (**Fig. 11**). Upon conducting a principal component analysis (PCA), we observed that the trait representing the mean convex hull area (*chull_area_mean*) emerged with one of the highest coefficient magnitudes in the linear transformation pertaining to the first principal component. This indicates a predominant influence of this trait on the variance captured by the first principal component. Similarly, the median distal lateral root angle (*lateral_angles_distal_median_median*) demonstrated a significant coefficient magnitude in relation to the second principal component, suggesting its considerable impact on the variance described by this component. These findings illustrate that these two traits play a major role in their respective principal component axes, providing an orthogonal representation of the trait space within the first two dimensions, which accounted for 50.62% of the total explained variance. When these traits were used to inform the coloring of the UMAP, the resulting visualization offered an interpretable map of the phenotypic trait space. The traits serve as effective indicators within the UMAP, enabling clear differentiation and understanding of the complex, nonlinear relationships among the samples. Using UMAP, we find that these plants span a continuous space characterized by different high variance traits, including the mean convex hull area (**Fig. 11A**) and the median lateral root angles (**Fig. 11B**).

## 4. Conclusion

We have developed pipelines for efficient and accurate root trait phenotyping using a pose estimation-based approach. While previous approaches have leveraged landmark detection [13], here we show the feasibility of a segmentation-free, landmark localization and grouping pipeline for RSA phenotyping.

In summary, the key contributions of this work include:

i. We use SLEAP [22] to train pose estimation models to localize and group landmarks on primary and lateral roots across a range of plant species, including the crop plants soybean (*Glycine max)*, rice (*Oryza sativa)*, canola (*Brassica napus*), and the model plant Arabidopsis (*Arabidopsis thaliana*). We show that our pipelines can detect root landmarks with 0.087% root length or 0.540 mm (rice primary root 3 DAG) to 0.984% root length or 0.843 mm (Arabidopsis lateral 7 DAG) median localization error.
ii. We developed a plant RSA trait extraction tool (*sleap-roots*) that can extract up to 1035 traits per plant using predicted landmarks. We show that the computed traits are highly accurate, with 96.2% (canola) to 99.5% (Arabidopsis) of individual trait values derived from fully automated inference falling within 1 standard deviation from those derived from manually proofread data.
iii. We show that accurate pose estimation-based root landmark detection can be achieved with <100 labeled images for most datasets, and is 1.5x faster to annotate and 10x faster to train and predict than segmentation-based approaches.
iv. We demonstrate the applicability of pose estimation-derived root traits for common downstream analyses, such as genotype classification and large-scale phenotypic trait mapping.
v. We provide an exhaustive list of traits and their estimated accuracies using our automated pipeline for practitioners (**Supplementary Materials**).
vi. We make all data, code, and models openly available: https://github.com/talmolab/sleap-roots, https://osf.io/k7j9g/.

### 4.1 Limitations and Future Directions

While this work demonstrates clear advantages to pose estimation-based plant phenotyping, we note that this approach has inherent limitations:

i. We approximate root centerlines using a fixed number of landmarks per model, which means that complex root curvatures will not be as well captured in some cases. Future work may be able to generalize our pose estimation algorithm to variable numbers of landmarks to resolve this.
ii. Root widths are not as reliably and densely estimated using our lateral root-matching method as would be possible via segmentation since we explicitly do not predict root boundaries. Future work combining segmentation with pose estimation may resolve this limitation.
iii. While our explicit representation of connectivity is more robust to self-intersections than segmentation-based postprocessing (e.g., skeletonization, connected components), dense and highly overlapping root systems are challenging in either case. Here we mitigate this by computing summary measures of traits across 72 different viewpoints using a cylindrical imaging system, under the assumption that ambiguities are resolvable in sufficient views. This will not be the case in single-view imaging systems, and self-intersections may present more significant challenges in 2D growth systems (e.g., plates).
iv. For manually measured root traits, the annotator selected one frame among the 72 frames that had the fewest occlusions. To evaluate the SLEAP-based root trait estimation accuracy, we used the median value summarized across all 72 frames. The majority of traits comparison were good (**Fig. 6A, C, D**), while the SLEAP-based maximum primary root lengths were smaller than manually measured ones (**Fig. 6B**). Future work comparing to manual annotation may provide more nuanced views on the automated system performance by including repeated manual trait measurements on the same images, and measurements across all images to circumvent the frame selection bias.
v. Despite our use of multi-view data, we do not estimate root landmarks in 3D. This is achievable using standard triangulation-based methods, but will require new approaches for matching detections across views. Datasets with heavy occlusion from overlapping roots, such as the rice seminal 10 DAG model (**Fig. 5C**), would benefit the most from root traits using 3D reconstructions.
vi. As mentioned in the methods section, all models were trained using the same hyperparameters. In the future, optimal hyperparameters could be found for each model. This would make the most difference for Arabidopsis models (**Fig. 5C**, **S2**) which have smaller and thinner roots than the crop plants.

## Acknowledgments

We thank Pamela Ronald and Artur Teixeira de Araujo Junior (University of California, Davis) for providing the rice seed from the fast-neutron-induced mutant collection, Henry T. Nguyen and Heng Ye (University of Missouri, Columbia) for providing seeds for soybean lines, and Michael J. Stamm (Kansas State University, Kansas) for providing the seeds for canola lines. We thank Shane Hunt, Nolan Mitschke, and Chad Morris for maintaining the greenhouse in Encinitas. We thank Mira Gowda for her contributions to this project as a summer intern. We thank Dan Butler for valuable discussions and insights.

## Author contributions

Conceptualization: EB, WB, TP Data curation: EB, LW

Investigation (data collection and annotation): EB, HC, KE, MK, JT, AA, EM, ES, CC, LR, CG, DR, AT, JG, MR

Supervision: SP, SL, LZ, AR, ED, MR, WB, TP

Software: EB, LW, LM, TP Writing: EB, LW, SP, WB, TP

## Funding

This work was supported by gifts from the Bezos Earth Fund, the Hess Corporation, through the TED Audacious Project, and the National Institutes of Health (1RF1MH132653).

## Competing interests

W.B. is a co-founder of Cquesta, a company that works on crop root growth and carbon sequestration.

## Data Availability

The datasets generated and/or analyzed during the current study are available in the Open Science Framework (OSF) repository. This includes the labeled data, predictive models, and analysis files which can be accessed via the following link: https://osf.io/k7j9g/.

The codebase used for the root tracking and analysis is available on GitHub under the *sleap-roots* repository and can be accessed here: https://github.com/talmolab/sleap-roots. Additionally, the specific code utilized for replicating the figures presented in this study can be found in a separate GitHub repository here: https://github.com/talmolab/Berrigan_et_al_sleap-roots.

Researchers interested in replicating our findings or conducting further analysis can find all necessary resources at the aforementioned links. For any further inquiries or assistance regarding the datasets or code, please contact the corresponding authors.

---

**Fig. S1.**
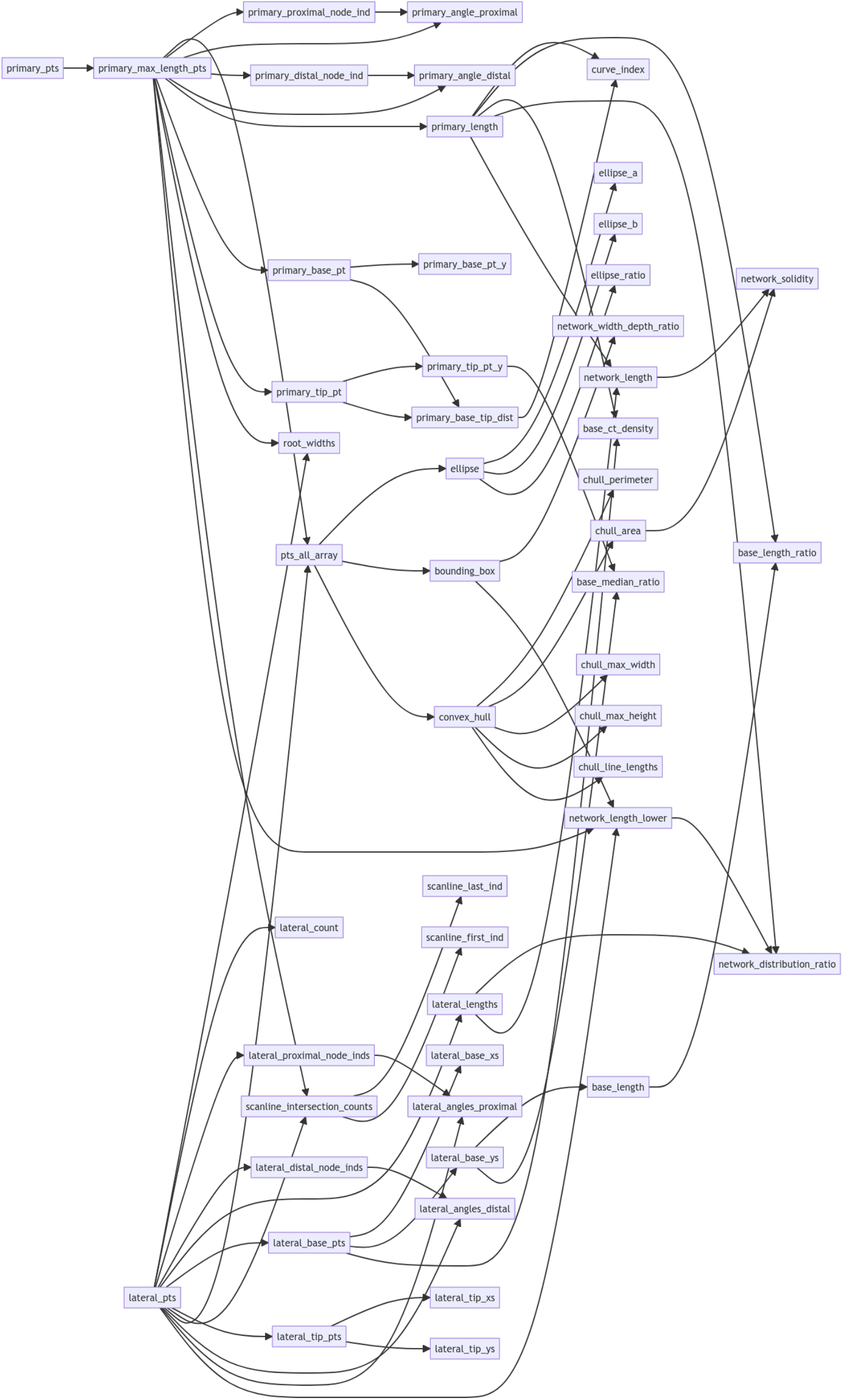
Dicot trait computation pipeline. Schematic of the trait computation pipeline for dicot species. Nodes correspond to traits or intermediate values, while arrows represent the dependencies between them in the computation graph.

**Fig. S2.**
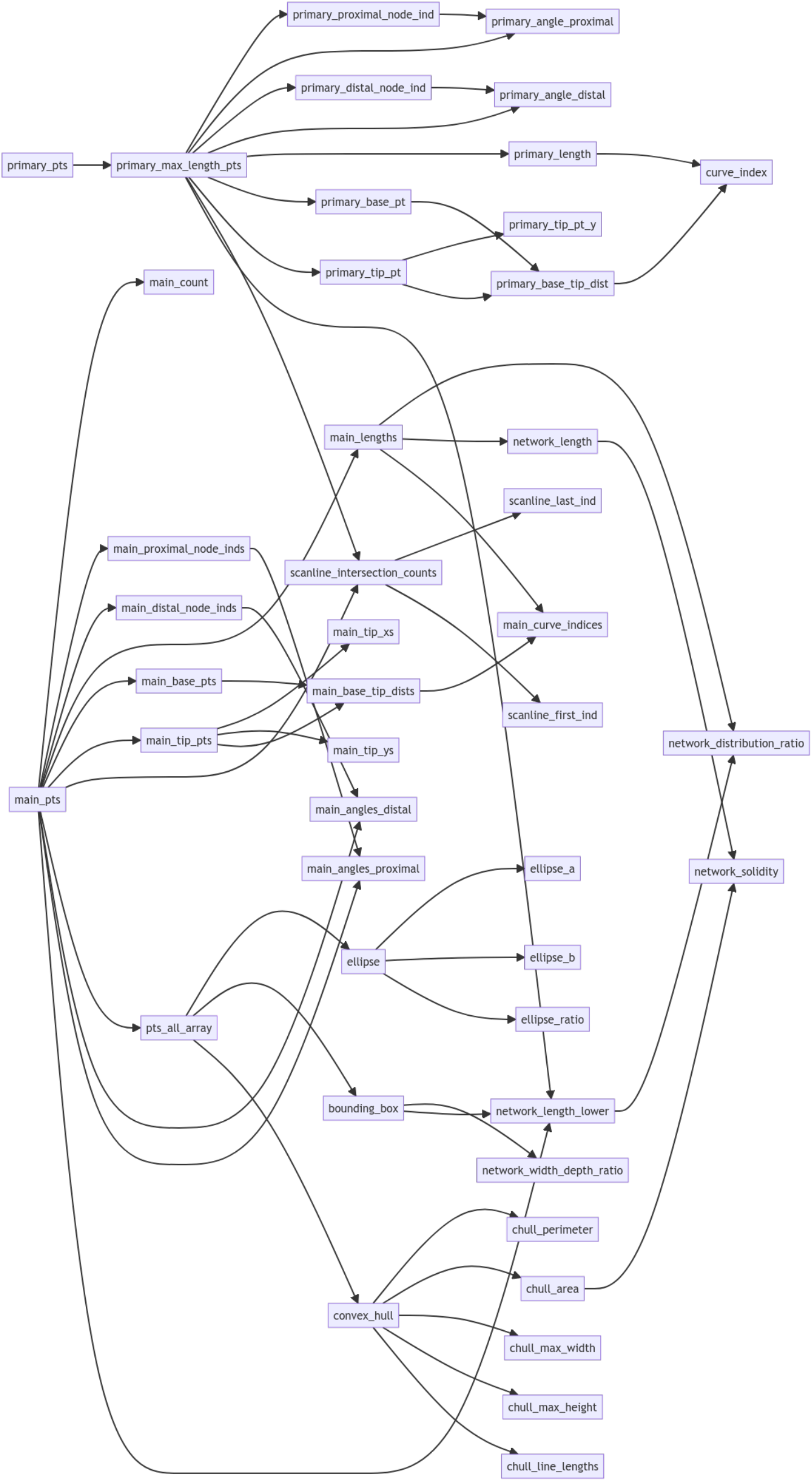
Younger monocot trait computation pipeline. Schematic of the trait computation pipeline for monocot species at early ages. Nodes correspond to traits or intermediate values, while arrows represent the dependencies between them in the computation graph.

**Table S1.** Number of labeled frames in each test set for Figure 5.

| **Model name** | **Test set replicate** | **Number of labeled frames in the test set** |
| --- | --- | --- |
| Soybean primary roots (5-8 DAG) | 1 | 131 |
| Soybean primary roots (5-8 DAG) | 2 | 139 |
| Soybean primary roots (5-8 DAG) | 3 | 112 |
| Soybean lateral roots (5-8 DAG) | 1 | 42 |
| Soybean lateral roots (5-8 DAG) | 2 | 50 |
| Soybean lateral roots (5-8 DAG) | 3 | 41 |
| Canola primary roots (5-13 DAG) | 1 | 30 |
| Canola primary roots (5-13 DAG) | 2 | 33 |
| Canola primary roots (5-13 DAG) | 3 | 32 |
| Canola lateral roots (5-13 DAG) | 1 | 41 |
| Canola lateral roots (5-13 DAG) | 2 | 39 |
| Canola lateral roots (5-13 DAG) | 3 | 45 |
| Arabidopsis primary roots (7 DAG) | 1 | 6 |
| Arabidopsis primary roots (7 DAG) | 2 | 6 |
| Arabidopsis primary roots (7 DAG) | 3 | 6 |
| Arabidopsis lateral roots (7 DAG) | 1 | 30 |
| Arabidopsis lateral roots (7 DAG) | 2 | 26 |
| Arabidopsis lateral roots (7 DAG) | 3 | 32 |
| Rice primary root (3 DAG) | 1 | 82 |
| Rice primary root (3 DAG) | 2 | 70 |
| Rice primary root (3 DAG) | 3 | 66 |
| Rice seminal roots (3 DAG) | 1 | 85 |
| Rice seminal roots (3 DAG) | 2 | 81 |
| Rice seminal roots (3 DAG) | 3 | 72 |
| Rice seminal roots (10 DAG) | 1 | 63 |
| Rice seminal roots (10 DAG) | 2 | 62 |
| Rice seminal roots (10 DAG) | 3 | 56 |

**Table S2.** Summary of results presented in Figure 5. The median localization error for each test set was normalized using the average root length (over all test sets for that model). The average values per model of the median localization error are listed in the table below and plotted in Figure 5.

| Model | Median error (mm) | Average Root Length (mm) | Median error (% root length) |
| --- | --- | --- | --- |
| Arabidopsis primary (7 DAG) | 1.537197447 | 326.8307267 | 0.470334434 |
| Canola primary (5-13 DAG) | 1.282755947 | 512.8514316 | 0.250122329 |
| Rice primary (3 DAG) | 0.53964299 | 618.6669212 | 0.087226741 |
| Soybean primary (5-8 DAG) | 1.668540805 | 784.0171943 | 0.212819415 |
| Arabidopsis lateral (7 DAG) | 0.84270209 | 85.57050619 | 0.984804377 |
| Canola lateral (5-13 DAG) | 0.365802449 | 94.90145824 | 0.385455035 |
| Soybean lateral (5-8 DAG) | 0.327267925 | 118.7363843 | 0.275625645 |
| Rice seminal (10 DAG) | 2.2808718 | 553.2915721 | 0.412236859 |
| Rice seminal (3 DAG) | 0.661920696 | 336.6339403 | 0.196629221 |

**Table S3.** Pearson correlation of proofread and non-proofread canola traits for Figure 7A. Only the traits with top 50 or bottom 50 correlation values are displayed.

| Trait category | Trait name | Correlation |
| --- | --- | --- |
| Scanline | scanline_intersection_counts_median_median | 1.0000 |
| Scanline | scanline_intersection_counts_median_p25 | 1.0000 |
| Scanline | scanline_intersection_counts_median_p75 | 1.0000 |
| Scanline | scanline_intersection_counts_p75_min | 1.0000 |
| Scanline | scanline_intersection_counts_p75_p25 | 1.0000 |
| Scanline | scanline_last_ind_p95 | 1.0000 |
| Scanline | scanline_last_ind_median | 1.0000 |
| Scanline | scanline_last_ind_p75 | 1.0000 |
| Length | primary_tip_pt_y_p95 | 1.0000 |
| Length | primary_tip_pt_y_p75 | 1.0000 |
| Length | primary_tip_pt_y_median | 1.0000 |
| Scanline | scanline_last_ind_p25 | 1.0000 |
| Length | primary_tip_pt_y_p25 | 1.0000 |
| Length | primary_tip_pt_y_max | 0.9999 |
| Tip Base | lateral_base_xs_median_mean | 0.9999 |
| Tip Base | lateral_base_xs_mean_mean | 0.9999 |
| Tip Base | lateral_base_xs_p25_mean | 0.9999 |
| Tip Base | lateral_base_xs_p75_mean | 0.9998 |
| Tip Base | lateral_base_xs_p5_mean | 0.9998 |
| Tip Base | lateral_base_xs_p75_p75 | 0.9998 |
| Tip Base | lateral_base_xs_median_p95 | 0.9997 |
| Tip Base | lateral_base_xs_p75_p95 | 0.9997 |
| Scanline | scanline_last_ind_mean | 0.9997 |
| Scanline | scanline_last_ind_max | 0.9997 |
| Tip Base | lateral_tip_xs_p5_p25 | 0.9997 |
| Tip Base | lateral_base_xs_max_p75 | 0.9997 |
| Tip Base | lateral_base_xs_p95_mean | 0.9997 |
| Tip Base | lateral_base_xs_max_min | 0.9997 |
| Tip Base | lateral_base_xs_mean_p95 | 0.9997 |
| Tip Base | lateral_tip_xs_min_p25 | 0.9996 |
| Tip Base | lateral_base_xs_min_mean | 0.9996 |
| Tip Base | lateral_base_xs_p5_max | 0.9996 |
| Tip Base | lateral_base_xs_min_p75 | 0.9996 |
| Tip Base | lateral_tip_xs_p5_mean | 0.9996 |
| Tip Base | lateral_tip_xs_min_mean | 0.9996 |
| Tip Base | lateral_base_xs_p95_p75 | 0.9996 |
| Tip Base | lateral_base_xs_max_mean | 0.9996 |
| Tip Base | lateral_base_xs_p25_median | 0.9996 |
| Tip Base | lateral_base_xs_p25_p95 | 0.9996 |
| Tip Base | lateral_base_xs_p95_min | 0.9996 |
| Tip Base | lateral_base_xs_median_p75 | 0.9996 |
| Tip Base | lateral_tip_xs_mean_mean | 0.9996 |
| Tip Base | lateral_base_xs_median_std | 0.9995 |
| Tip Base | lateral_base_xs_mean_p25 | 0.9995 |
| Tip Base | lateral_base_xs_p5_p75 | 0.9995 |
| Tip Base | lateral_base_xs_median_median | 0.9995 |
| Tip Base | lateral_base_xs_p5_p25 | 0.9995 |
| Tip Base | lateral_base_xs_p25_p25 | 0.9995 |
| Tip Base | lateral_tip_xs_p95_mean | 0.9995 |
| Tip Base | lateral_base_xs_min_p25 | 0.9995 |
| ... | ... | ... |
| Tip Base | lateral_base_ys_max_std | 0.7819 |
| Length | lateral_lengths_max_max | 0.7805 |
| Root Width | root_widths_p5_p25 | 0.7764 |
| Root Width | root_widths_min_p25 | 0.7763 |
| Scanline | scanline_intersection_counts_median_max | 0.7756 |
| Root Width | root_widths_p25_p25 | 0.7755 |
| Length | base_length_std | 0.7732 |
| Tip Base | lateral_tip_ys_std_max | 0.7642 |
| Length | lateral_lengths_max_std | 0.7642 |
| Tip Base | lateral_base_ys_std_std | 0.7434 |
| Length | lateral_lengths_std_max | 0.7411 |
| Length | grav_index_max | 0.7401 |
| Convex Hull | chull_line_lengths_min_min | 0.7390 |
| Ellipse | ellipse_b_max | 0.7383 |
| Tip Base | lateral_tip_ys_max_std | 0.7292 |
| Tip Base | lateral_tip_ys_std_std | 0.7273 |
| Root Width | root_widths_max_p5 | 0.7224 |
| Root Width | root_widths_p95_p5 | 0.7205 |
| Root Width | root_widths_p75_min | 0.7189 |
| Length | primary_tip_pt_y_min | 0.7188 |
| Root Width | root_widths_max_min | 0.7180 |
| Root Width | root_widths_p95_min | 0.7179 |
| Root Width | root_widths_p75_p5 | 0.7136 |
| Root Width | root_widths_median_min | 0.7112 |
| Root Width | root_widths_mean_min | 0.7112 |
| Root Width | root_widths_median_p5 | 0.6997 |
| Root Width | root_widths_mean_p5 | 0.6983 |
| Angle | lateral_angles_distal_p95_max | 0.6799 |
| Ellipse | ellipse_a_max | 0.6761 |
| Root Width | root_widths_p25_p5 | 0.6712 |
| Root Width | root_widths_p25_min | 0.6709 |
| Angle | lateral_angles_proximal_min_min | 0.6684 |
| Root Width | root_widths_p5_p5 | 0.6486 |
| Root Width | root_widths_min_p5 | 0.6451 |
| Root Width | root_widths_p5_min | 0.6443 |
| Length | base_median_ratio_std | 0.6443 |
| Root Width | root_widths_min_min | 0.6397 |
| Root Width | root_widths_max_std | 0.6311 |
| Root Width | root_widths_p95_std | 0.6283 |
| Root Width | root_widths_min_std | 0.6272 |
| Root Width | root_widths_p5_std | 0.6250 |
| Root Width | root_widths_p75_std | 0.6202 |
| Root Width | root_widths_p25_std | 0.6187 |
| Root Width | root_widths_mean_std | 0.6164 |
| Root Width | root_widths_median_std | 0.6163 |
| Angle | lateral_angles_distal_max_max | 0.6085 |
| Length | base_median_ratio_max | 0.5622 |
| Network | network_solidity_std | 0.5480 |
| Length | primary_tip_pt_y_std | 0.5181 |
| Network | network_solidity_max | 0.4764 |

**Table S4.** Pearson correlation of proofread and non-proofread soybean traits for Figure 7B. Only the traits with top 50 or bottom 50 correlation values are displayed.

| Trait category | Trait name | Correlation |
| --- | --- | --- |
| Tip Base | lateral_tip_ys_p75_p25 | 0.9997 |
| Tip Base | lateral_base_xs_mean_mean | 0.9996 |
| Scanline | scanline_last_ind_max | 0.9996 |
| Tip Base | lateral_tip_ys_median_median | 0.9996 |
| Tip Base | lateral_tip_ys_p25_mean | 0.9995 |
| Tip Base | lateral_tip_xs_min_p75 | 0.9995 |
| Tip Base | lateral_tip_ys_median_p5 | 0.9995 |
| Tip Base | lateral_base_xs_p25_mean | 0.9995 |
| Tip Base | lateral_tip_ys_mean_p25 | 0.9995 |
| Tip Base | lateral_tip_ys_p75_median | 0.9995 |
| Tip Base | lateral_tip_xs_min_mean | 0.9994 |
| Tip Base | lateral_base_xs_median_mean | 0.9994 |
| Tip Base | lateral_base_xs_p5_p5 | 0.9994 |
| Tip Base | lateral_base_ys_p75_p75 | 0.9994 |
| Tip Base | lateral_tip_ys_median_mean | 0.9994 |
| Scanline | scanline_first_ind_p75 | 0.9994 |
| Tip Base | lateral_base_ys_p5_mean | 0.9994 |
| Tip Base | lateral_base_xs_mean_std | 0.9993 |
| Tip Base | lateral_tip_ys_p95_p25 | 0.9993 |
| Tip Base | lateral_tip_ys_p25_p75 | 0.9993 |
| Tip Base | lateral_base_ys_p5_p75 | 0.9993 |
| Tip Base | lateral_base_xs_median_std | 0.9993 |
| Scanline | scanline_last_ind_p95 | 0.9993 |
| Tip Base | lateral_tip_ys_min_max | 0.9993 |
| Tip Base | lateral_base_ys_p75_median | 0.9992 |
| Tip Base | lateral_base_ys_median_median | 0.9992 |
| Tip Base | lateral_tip_ys_p75_mean | 0.9992 |
| Tip Base | lateral_base_ys_min_p95 | 0.9992 |
| Convex Hull | chull_max_height_max | 0.9992 |
| Tip Base | lateral_base_xs_p5_std | 0.9992 |
| Tip Base | lateral_tip_ys_mean_median | 0.9992 |
| Convex Hull | chull_max_height_p95 | 0.9992 |
| Tip Base | lateral_base_ys_median_mean | 0.9992 |
| Tip Base | lateral_base_ys_min_mean | 0.9991 |
| Tip Base | lateral_base_ys_p75_p25 | 0.9991 |
| Tip Base | lateral_base_xs_p75_std | 0.9991 |
| Tip Base | lateral_base_ys_p75_mean | 0.9991 |
| Tip Base | lateral_tip_ys_p95_p5 | 0.9990 |
| Tip Base | lateral_tip_ys_mean_mean | 0.9990 |
| Tip Base | lateral_tip_ys_median_p25 | 0.9990 |
| Tip Base | lateral_tip_ys_p5_max | 0.9990 |
| Tip Base | lateral_base_xs_p25_std | 0.9989 |
| Tip Base | lateral_tip_ys_p5_mean | 0.9989 |
| Tip Base | lateral_base_xs_min_std | 0.9989 |
| Tip Base | lateral_tip_xs_std_mean | 0.9989 |
| Tip Base | lateral_tip_ys_mean_p5 | 0.9988 |
| Tip Base | lateral_base_ys_p5_p25 | 0.9988 |
| Tip Base | lateral_tip_ys_p5_p95 | 0.9988 |
| Tip Base | lateral_base_xs_min_p5 | 0.9988 |
| Tip Base | lateral_base_xs_mean_p5 | 0.9987 |
| ... | ... | ... |
| Angle | lateral_angles_distal_p95_p95 | 0.7398 |
| Root Width | root_widths_p75_min | 0.7394 |
| Root Width | root_widths_p95_min | 0.7380 |
| Length | base_length_ratio_p75 | 0.7379 |
| Root Width | root_widths_max_min | 0.7377 |
| Angle | lateral_angles_distal_max_p25 | 0.7362 |
| Root Width | root_widths_p25_min | 0.7354 |
| Length | primary_length_std | 0.7345 |
| Angle | lateral_angles_distal_p95_p25 | 0.7331 |
| Angle | lateral_angles_distal_max_p95 | 0.7306 |
| Angle | lateral_angles_distal_p95_mean | 0.7274 |
| Length | primary_length_min | 0.7249 |
| Angle | lateral_angles_proximal_std_min | 0.7238 |
| Angle | lateral_angles_distal_max_mean | 0.7235 |
| Angle | lateral_angles_distal_max_p75 | 0.7228 |
| Angle | lateral_angles_distal_p95_p75 | 0.7216 |
| Tip Base | lateral_base_xs_std_min | 0.7189 |
| Angle | lateral_angles_distal_p95_median | 0.7171 |
| Angle | lateral_angles_proximal_std_p5 | 0.7163 |
| Angle | lateral_angles_distal_std_p25 | 0.7099 |
| Length | grav_index_p75 | 0.7097 |
| Angle | lateral_angles_distal_max_median | 0.7020 |
| Length | lateral_lengths_min_min | 0.6924 |
| Angle | lateral_angles_proximal_min_p5 | 0.6867 |
| Length | grav_index_mean | 0.6854 |
| Length | base_ct_density_p95 | 0.6831 |
| Angle | lateral_angles_distal_min_min | 0.6751 |
| Length | grav_index_median | 0.6552 |
| Scanline | scanline_first_ind_p95 | 0.6423 |
| Length | base_ct_density_std | 0.6364 |
| Length | base_median_ratio_std | 0.6255 |
| Length | grav_index_p25 | 0.6225 |
| Scanline | scanline_first_ind_max | 0.6143 |
| Length | grav_index_p5 | 0.6089 |
| Angle | lateral_angles_distal_p5_min | 0.6069 |
| Angle | primary_angle_distal_min | 0.5986 |
| Angle | lateral_angles_proximal_p5_min | 0.5793 |
| Network | network_distribution_ratio_std | 0.5617 |
| Angle | lateral_angles_distal_std_min | 0.5509 |
| Convex Hull | chull_max_height_std | 0.5453 |
| Scanline | scanline_last_ind_std | 0.5290 |
| Scanline | scanline_first_ind_std | 0.5260 |
| Length | base_ct_density_max | 0.5058 |
| Length | base_length_ratio_std | 0.4983 |
| Angle | lateral_angles_distal_std_p5 | 0.4849 |
| Length | base_length_ratio_p95 | 0.4705 |
| Length | base_length_ratio_max | 0.4380 |
| Length | primary_tip_pt_y_std | 0.3997 |
| Length | grav_index_min | 0.3441 |
| Angle | lateral_angles_proximal_min_min | 0.3379 |

**Table S5.** Pearson correlation of proofread and non-proofread rice traits for Figure 7C. Only the traits with top 50 or bottom 50 correlation values were displayed.

| Trait category | Trait name | Correlation |
| --- | --- | --- |
| Scanline | scanline_intersection_counts_max_min | 1.0000 |
| Angle | main_angles_distal_std_min | 1.0000 |
| Scanline | scanline_intersection_counts_median_median | 1.0000 |
| Length | main_grav_indices_std_min | 1.0000 |
| Angle | main_angles_distal_std_p5 | 1.0000 |
| Scanline | scanline_intersection_counts_p75_median | 1.0000 |
| Angle | main_angles_proximal_min_min | 1.0000 |
| Length | main_base_tip_dists_std_min | 1.0000 |
| Scanline | scanline_intersection_counts_max_p25 | 1.0000 |
| Scanline | scanline_intersection_counts_median_p25 | 1.0000 |
| Angle | main_angles_proximal_std_min | 1.0000 |
| Length | main_count_p5 | 1.0000 |
| Tip Base | main_tip_ys_min_median | 1.0000 |
| Tip Base | main_tip_ys_p5_median | 1.0000 |
| Tip Base | main_tip_ys_max_median | 1.0000 |
| Length | main_grav_indices_min_p25 | 1.0000 |
| Length | main_base_tip_dists_min_p25 | 1.0000 |
| Length | main_base_tip_dists_p5_p25 | 1.0000 |
| Length | main_grav_indices_p5_p25 | 1.0000 |
| Tip Base | main_tip_ys_p25_median | 1.0000 |
| Scanline | scanline_last_ind_median | 1.0000 |
| Angle | main_angles_proximal_max_min | 1.0000 |
| Angle | main_angles_proximal_p95_min | 1.0000 |
| Length | main_lengths_p5_p75 | 1.0000 |
| Length | main_lengths_min_p75 | 1.0000 |
| Convex Hull | chull_max_height_median | 1.0000 |
| Convex Hull | chull_line_lengths_max_median | 1.0000 |
| Length | main_grav_indices_median_p25 | 1.0000 |
| Length | main_base_tip_dists_median_p25 | 1.0000 |
| Length | main_grav_indices_p5_p75 | 1.0000 |
| Length | main_base_tip_dists_p5_p75 | 1.0000 |
| Length | main_grav_indices_p25_p25 | 1.0000 |
| Length | main_base_tip_dists_p25_p25 | 1.0000 |
| Length | main_grav_indices_min_p95 | 1.0000 |
| Length | main_base_tip_dists_min_p95 | 1.0000 |
| Length | main_lengths_p25_median | 1.0000 |
| Length | main_grav_indices_min_p75 | 1.0000 |
| Length | main_base_tip_dists_min_p75 | 1.0000 |
| Length | main_lengths_p5_median | 1.0000 |
| Network | network_length_lower_median | 1.0000 |
| Length | main_lengths_min_median | 1.0000 |
| Tip Base | main_tip_ys_p25_p75 | 1.0000 |
| Length | main_base_tip_dists_p5_p95 | 0.9999 |
| Length | main_grav_indices_p5_p95 | 0.9999 |
| Angle | main_angles_proximal_p75_min | 0.9999 |
| Angle | main_angles_distal_p25_min | 0.9999 |
| Tip Base | main_tip_ys_min_max | 0.9999 |
| Convex Hull | chull_line_lengths_p95_median | 0.9999 |
| Tip Base | main_tip_xs_mean_mean | 0.9999 |
| Length | main_lengths_p25_p75 | 0.9999 |
| ... | ... | ... |
| Length | main_lengths_min_p5 | 0.9380 |
| Length | main_lengths_p5_p5 | 0.9377 |
| Tip Base | main_tip_xs_std_p95 | 0.9377 |
| Tip Base | main_tip_xs_p95_max | 0.9334 |
| Network | network_distribution_ratio_min | 0.9334 |
| Convex Hull | chull_area_p95 | 0.9330 |
| Ellipse | ellipse_b_p95 | 0.9327 |
| Length | main_lengths_mean_min | 0.9305 |
| Network | network_distribution_ratio_std | 0.9293 |
| Convex Hull | chull_line_lengths_p5_min | 0.9292 |
| Length | primary_length_p5 | 0.9284 |
| Network | network_distribution_ratio_median | 0.9272 |
| Convex Hull | chull_line_lengths_min_min | 0.9261 |
| Network | network_distribution_ratio_p25 | 0.9259 |
| Network | network_width_depth_ratio_std | 0.9249 |
| Length | primary_length_min | 0.9237 |
| Length | main_lengths_median_min | 0.9230 |
| Scanline | scanline_intersection_counts_median_std | 0.9205 |
| Length | main_lengths_p25_min | 0.9198 |
| Ellipse | ellipse_ratio_min | 0.9165 |
| Tip Base | main_tip_xs_max_max | 0.9161 |
| Scanline | scanline_last_ind_std | 0.9118 |
| Convex Hull | chull_line_lengths_p95_std | 0.9099 |
| Length | main_lengths_p5_min | 0.9086 |
| Convex Hull | chull_max_height_std | 0.9080 |
| Length | main_lengths_min_min | 0.9064 |
| Tip Base | main_tip_ys_p95_std | 0.9061 |
| Tip Base | main_tip_ys_max_std | 0.9049 |
| Convex Hull | chull_max_width_std | 0.9045 |
| Convex Hull | chull_area_max | 0.9032 |
| Angle | primary_angle_distal_p5 | 0.8999 |
| Tip Base | main_tip_xs_std_std | 0.8982 |
| Ellipse | ellipse_b_std | 0.8926 |
| Network | network_solidity_min | 0.8894 |
| Network | network_solidity_p5 | 0.8888 |
| Convex Hull | chull_max_width_max | 0.8875 |
| Convex Hull | chull_line_lengths_max_std | 0.8812 |
| Scanline | scanline_intersection_counts_p75_min | 0.8803 |
| Network | network_width_depth_ratio_max | 0.8784 |
| Convex Hull | chull_area_std | 0.8774 |
| Ellipse | ellipse_b_max | 0.8770 |
| Tip Base | main_tip_xs_std_max | 0.8699 |
| Network | network_distribution_ratio_max | 0.8676 |
| Network | network_length_lower_std | 0.8475 |
| Convex Hull | chull_perimeter_std | 0.8306 |
| Angle | primary_angle_proximal_min | 0.8269 |
| Length | primary_base_tip_dist_std | 0.8252 |
| Angle | primary_angle_distal_min | 0.8205 |
| Length | primary_length_std | 0.8130 |
| Length | primary_tip_pt_y_std | 0.8056 |

**Table S6.** Number of labeled frames in each test set for Figure 8.

| Model name | Number of labeled frames in the test set |
| --- | --- |
| Soybean primary roots (5-8 DAG) | 112 |
| Soybean lateral roots (5-8 DAG) | 41 |
| Canola primary roots (5-13 DAG) | 32 |
| Canola lateral roots (5-13 DAG) | 39 |
| Arabidopsis primary roots (7 DAG) | 6 |
| Arabidopsis lateral roots (7 DAG) | 26 |
| Rice primary root (3 DAG) | 66 |
| Rice seminal roots (3 DAG) | 72 |
| Rice seminal roots (10 DAG) | 56 |

## Notes

https://github.com/talmolab/sleap-roots

https://osf.io/k7j9g/

