## Supplemental Methods for "Fast and efficient root phenotyping via pose estimation"

### Arabidopsis 7DAP Lateral Labeling Rules

### Every root except the primary root is lateral root in this context

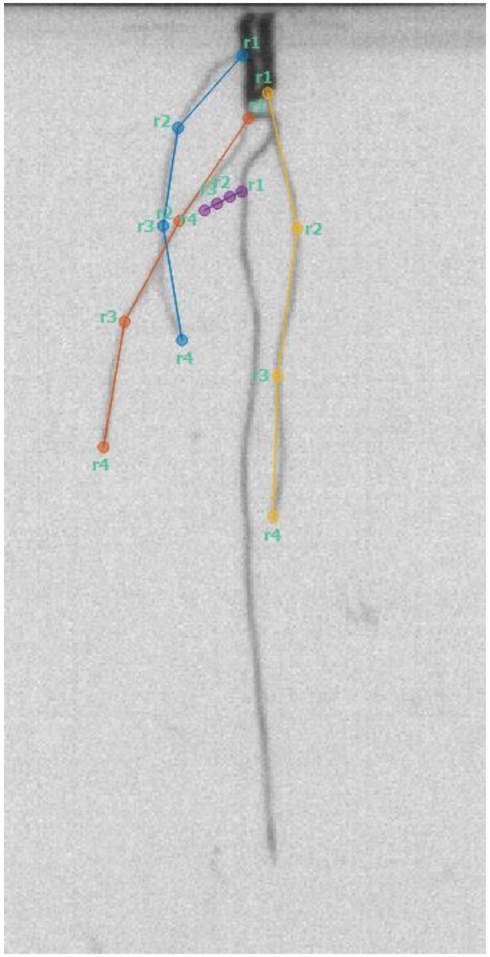

### Skeleton

- 4 nodes
  - {r1, r2, r3, r4}
- Connected
  - r1 -> r2, r2 -> r3, r3 -> r4

# r1

- Base of lateral root
- Furthest point up the root that is visible
- Sometime may be on the planting dark "hole"

### Evenly space the nodes as much as possible

### Center the nodes along the midline of the root

### EX. Of smallest root acceptable

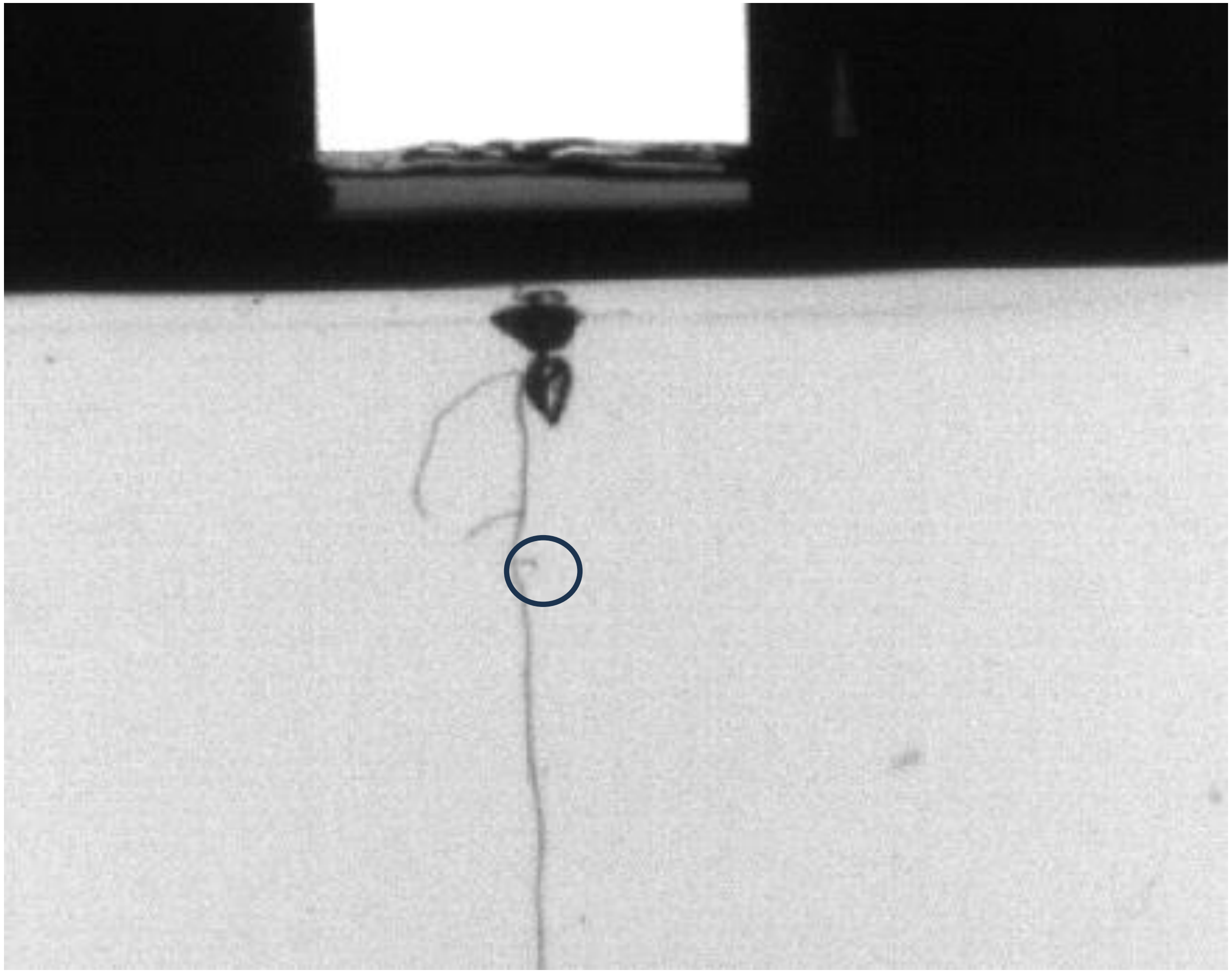

### EX. Too small to label

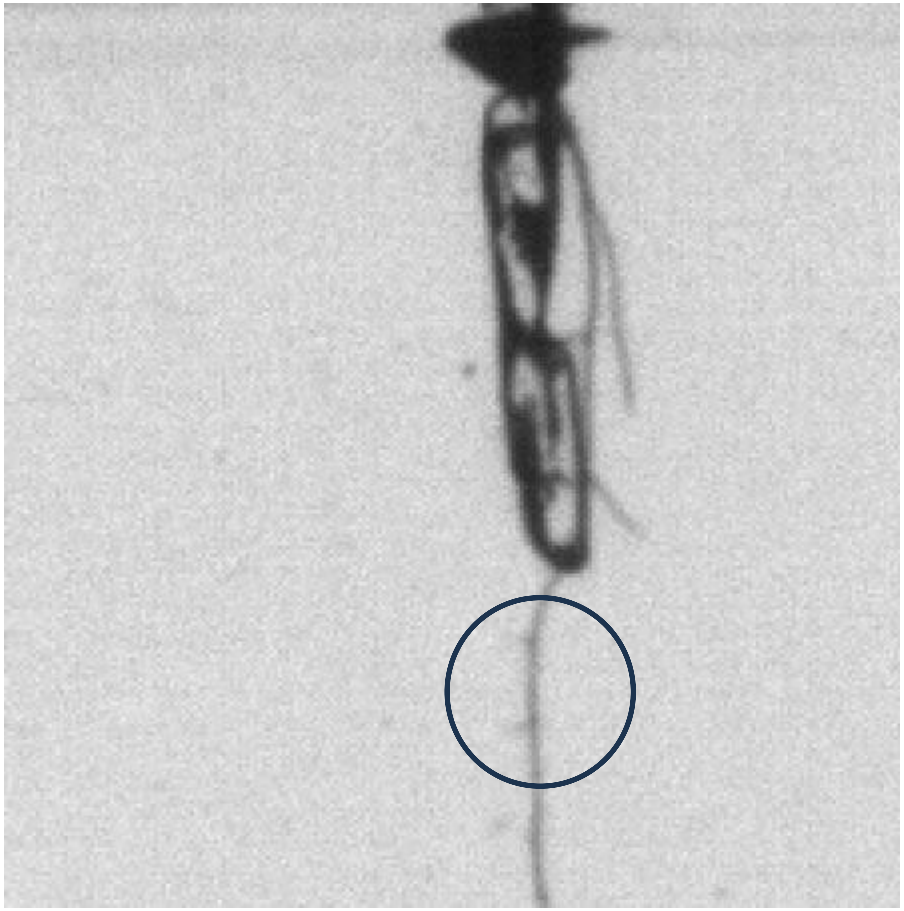

### PR will always be the longest

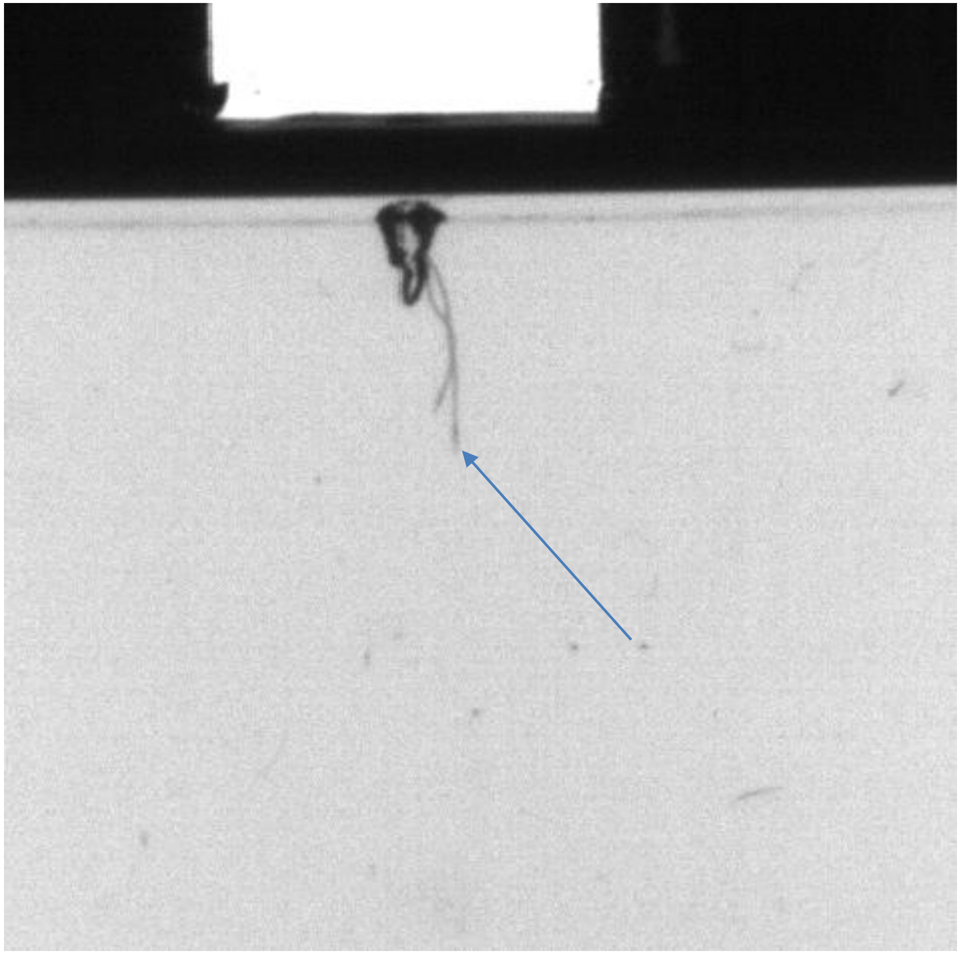

### Start 1st node at closest to PR as possible, even if on hole

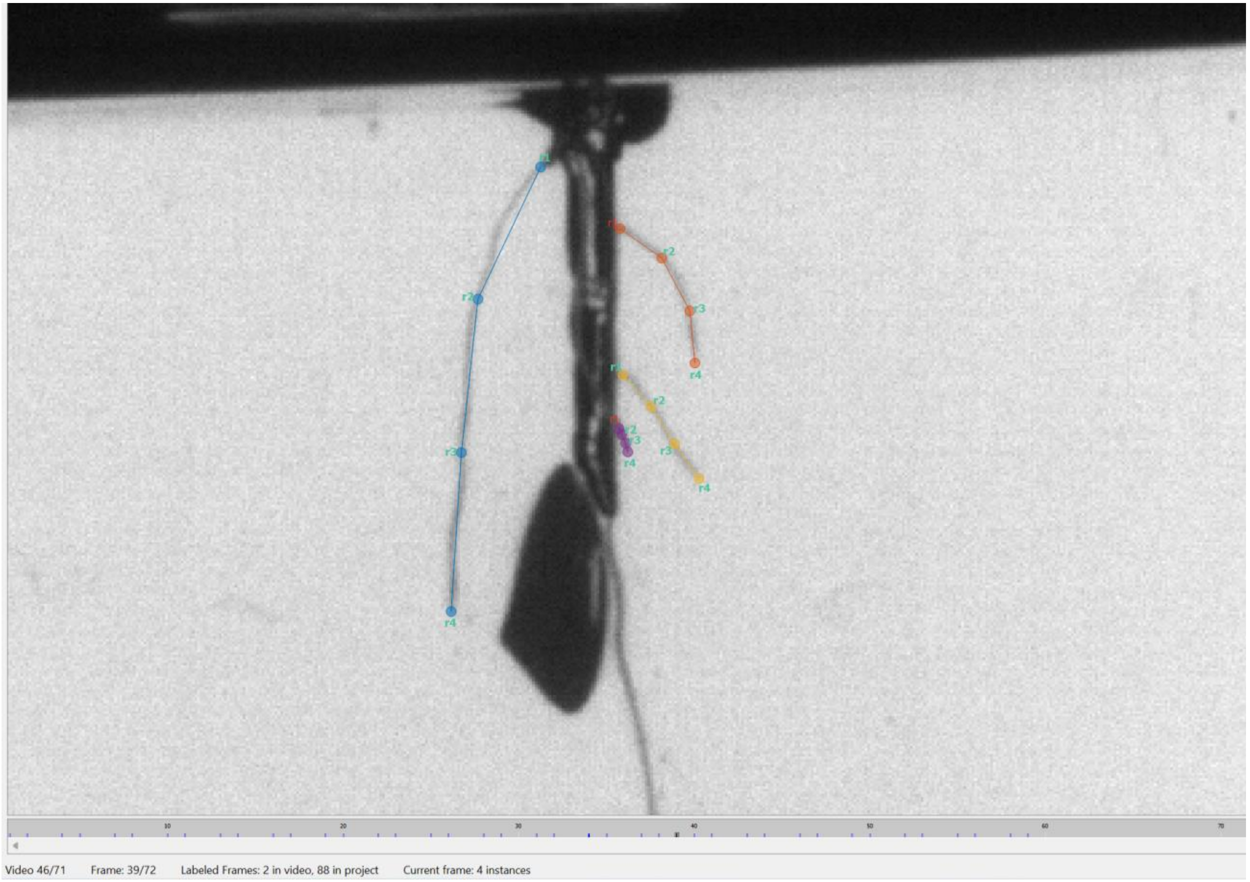

### Proof Reading LR

**Take out a timer. Time yourself per plant as you quickly go through the predictions using the right arrow. Look out for any anomalies such as:**
a. Portions of the background and cylinder detected as root
b. Important roots for the RSA not detected
c. (Note): criss-crossing is OK
d. (Note): ambiguities between primary and lateral root is OK if you cannot tell which one is which
e. (Note): Un-detected roots under 15 px is OK
f. (Note): primary root labeled as lateral root in absence of any lateral roots is OK

**If you feel that the predictions are hopeless, you need to start from scratch, and you would like these labels to be included in the training set** so that the predictions for that sort of plant are more accurate in the future, label that frame and record the frame number you labeled in the appropriate column (you can list more than one). I will add it to the labeled training set later.

**Note that labels must follow the labeling rules to be included in a training set.**
a. In this case, meaning predictions are bad enough that you would need to correct multiple predictions per frame, instead of proofreading all frames for this plant, just label 5 frames really well to include in the training set, delete all predictions from the plant (tool bar: label --> delete all predictions...) and make a note that you did this on the excel (along with adding which frames you labeled)

### Arabidopsis 7DAP Lateral Labeling Rules

### Every root except the primary root is lateral root in this context

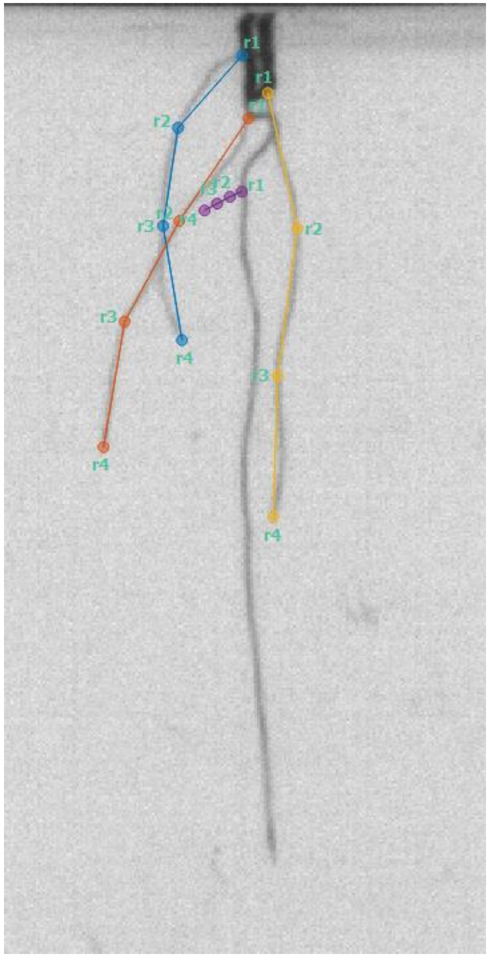

### Skeleton

- 4 nodes
  – {r1, r2, r3, r4}
- Connected
  – r1 -> r2, r2 -> r3, r3 -> r4

# r1

- Base of lateral root
- Furthest point up the root that is visible
- Sometime may be on the planting dark "hole"

### Evenly space the nodes as much as possible

### Center the nodes along the midline of the root

### EX. Of smallest root acceptable

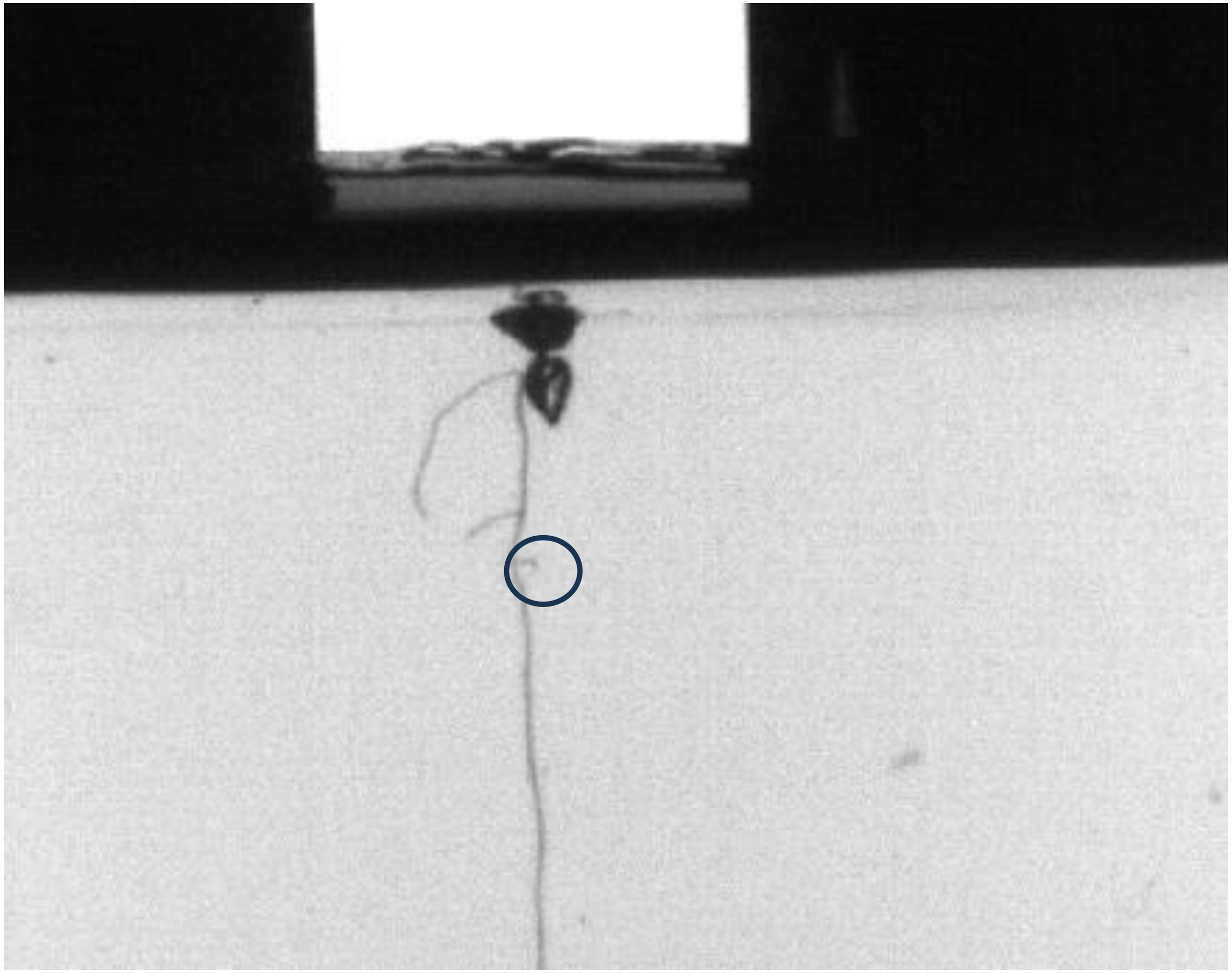

### EX. Too small to label

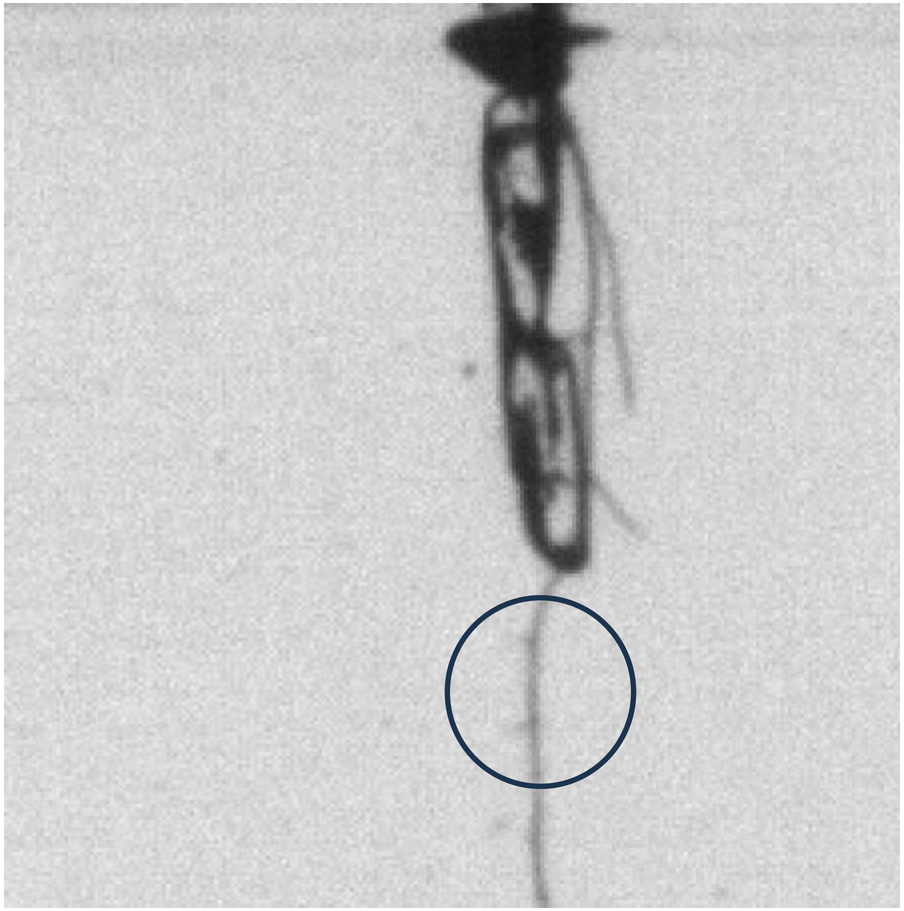

### PR will always be the longest

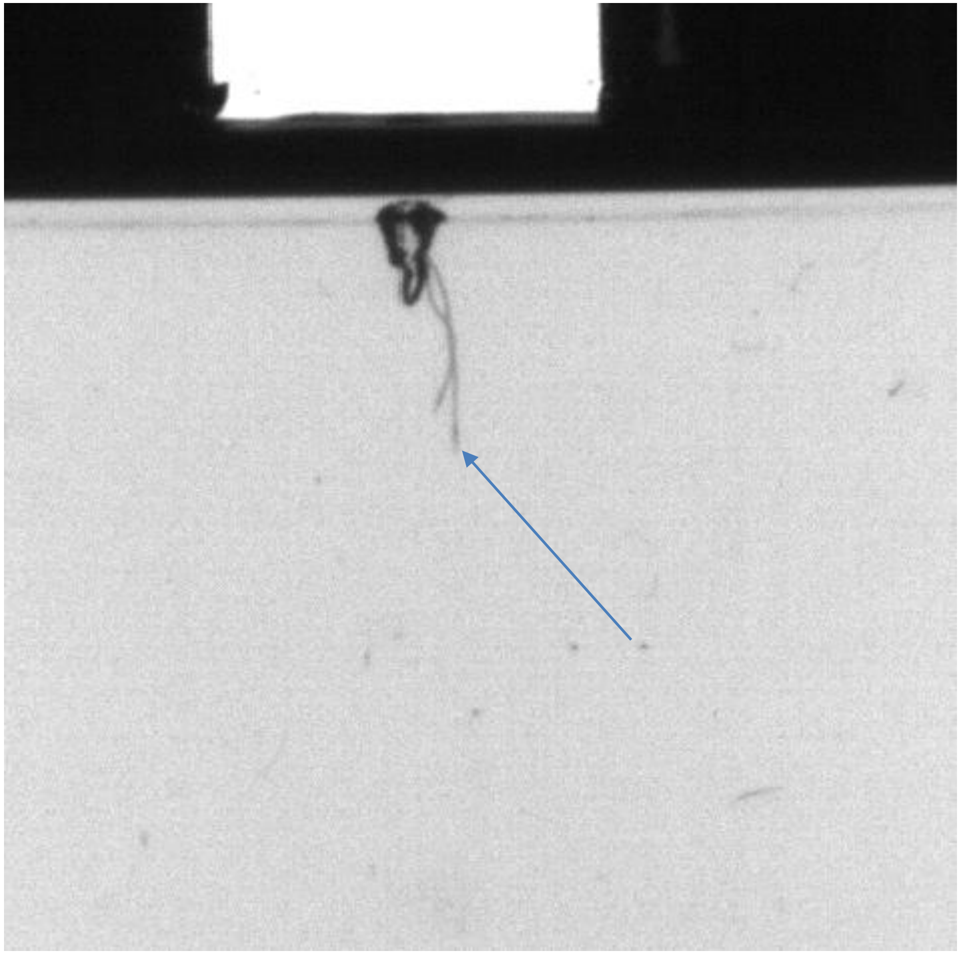

### Start 1st node at closest to PR as possible, even if on hole

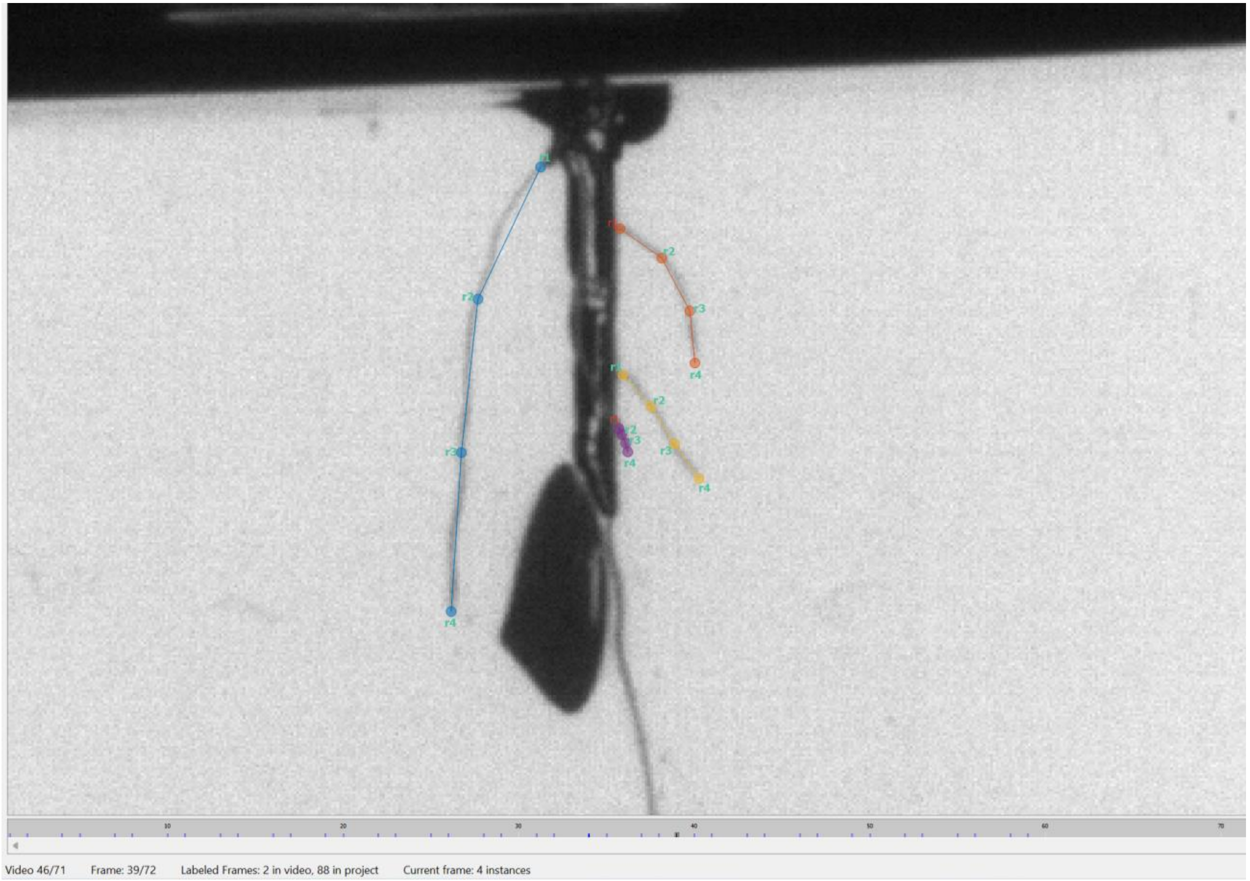

### Proof Reading LR

**Take out a timer. Time yourself per plant as you quickly go through the predictions using the right arrow. Look out for any anomalies such as:**
a. Portions of the background and cylinder detected as root
b. Important roots for the RSA not detected
c. (Note): criss-crossing is OK
d. (Note): ambiguities between primary and lateral root is OK if you cannot tell which one is which
e. (Note): Un-detected roots under 15 px is OK
f. (Note): primary root labeled as lateral root in absence of any lateral roots is OK

**If you feel that the predictions are hopeless, you need to start from scratch, and you would like these labels to be included in the training set** so that the predictions for that sort of plant are more accurate in the future, label that frame and record the frame number you labeled in the appropriate column (you can list more than one). I will add it to the labeled training set later.

**Note that labels must follow the labeling rules to be included in a training set.**
a. In this case, meaning predictions are bad enough that you would need to correct multiple predictions per frame, instead of proofreading all frames for this plant, just label 5 frames really well to include in the training set, delete all predictions from the plant (tool bar: label --> delete all predictions...) and make a note that you did this on the excel (along with adding which frames you labeled)

### Rice Main Root Labeling Rules

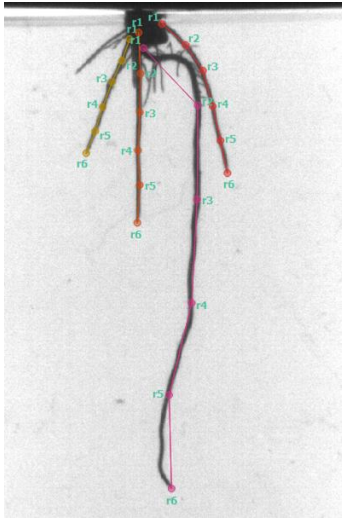

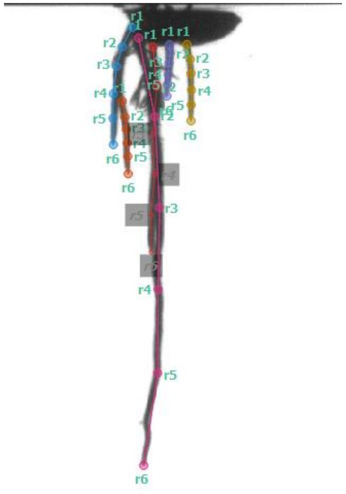

### Main Roots vs. Hairy Roots

- Main roots are the thicker and darker roots, and hairy roots come off of main roots.
- Only main roots should be labeled.

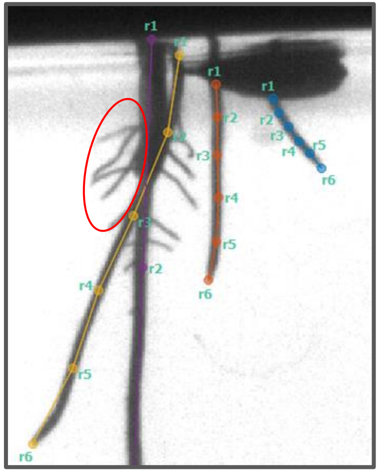

Hairy roots are circled in red—they should not be labeled

- Branching off of main roots and significantly thinner/shorter

### Skeleton of rice

```
"node_names": ["r1", "r2", "r3", "r4", "r5", "r6"],
"edge_inds": [[0, 1], [1, 2], [2, 3], [3, 4], [4, 5]]
```

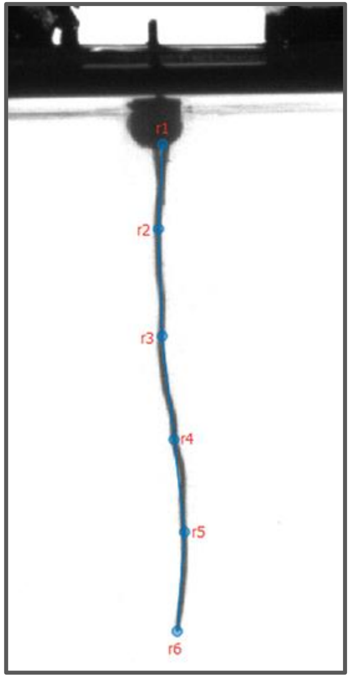


Roots are labelled using 6-node trees, which form a line.

### General Node Placement Rules

Nodes should be equally spaced along each root.

- Each 6-node instance should follow the length of one root
- [0, 1] = [1, 2]... = [5, 6]

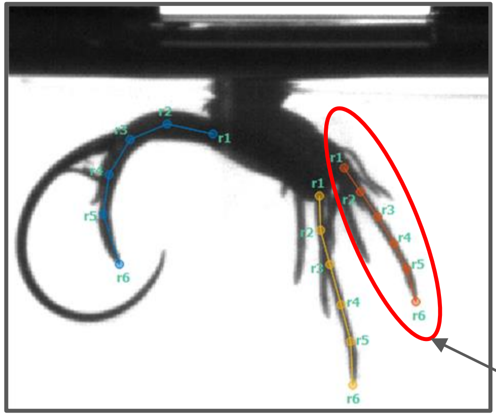

one instance/root

Nodes should be centered widthwise on the root.

- Zoom in to make sure they're centered

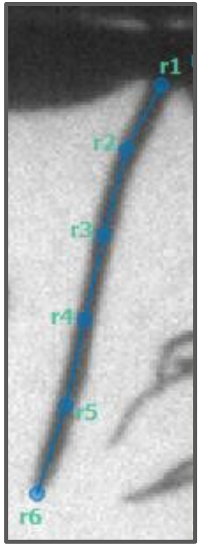

### Base Definition + Labelling Instructions

- The **base** of the root is the **first visible point** on an individual root and should be labelled as `r1`.
- If the bases of two roots overlap, `r1` on one of the roots should be moved down to the first distinguishable pint of the root that does not overlap with another root.
  - The bases of thicker and longer roots should be prioritized and kept higher.

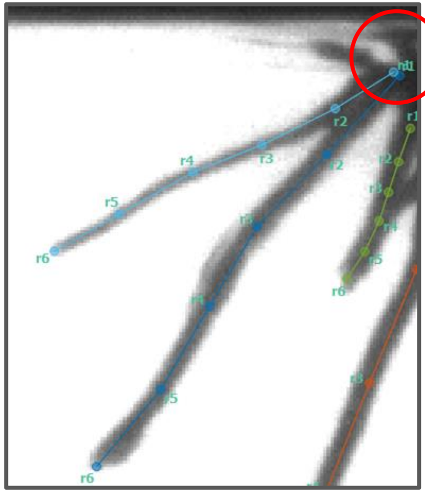

Incorrect base labelling

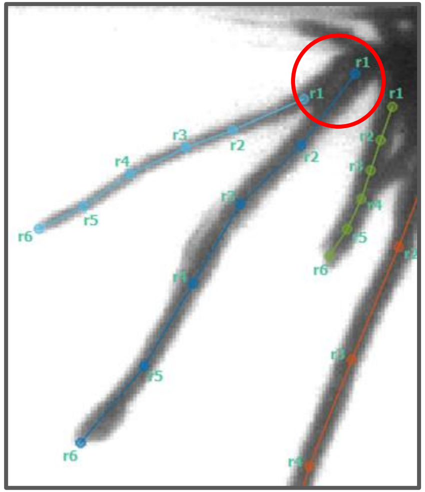

Correct base labelling

Although the bases of both leftmost roots (labeled in blue) overlap, their `r1` points should not be next to each other. The thicker root (on the right) should be prioritized and the `r1` point of the left root should move downwards to its first non-overlapping point.

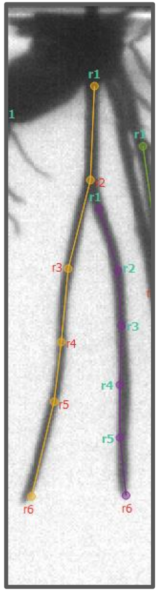

Another example of labeling `r1` lower in the case of overlap, and prioritizing the longer/thicker root.

### Tip Definition and Labelling

- The tip of the root is the last visible point, and should be labelled with `r6`.
    - `Remember to zoom in`

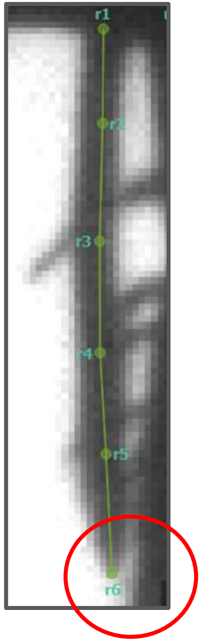

### SAVE

- Make sure to save frequently
- Changes are NOT automatically saved

### Consistency

- Consistency is key—if labels are not consistent they will have to be re-done
- If more than one person is labeling, make sure to discuss labeling preferences and special cases
    - Add new agreed-upon rules to slides to record special cases
- Go slowly and carefully rather than quickly

### Don't over label

- 10 Day-Old roots are very complicated
- Prioritize larger roots
- Prioritize more visible roots
- Prioritize roots important to overall RSA
- Minimize criss-crossing labels

### Prioritizing Roots

- If two roots have similar thickness/length, prioritize the one that is less obstructed by other roots
  - The non-prioritized one will be prioritized in a different frame

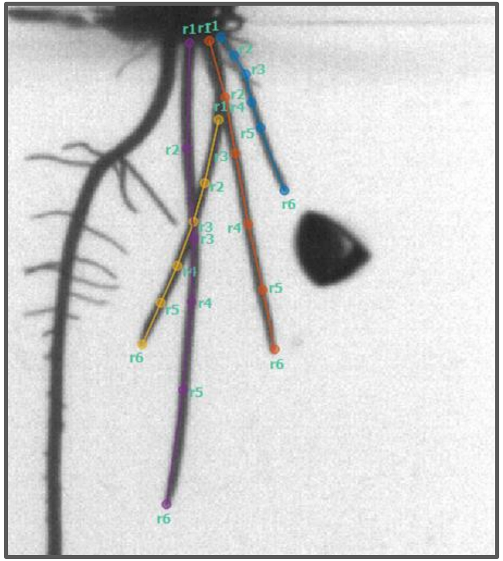

The yellow and orange roots are around the same length/thickness, but the yellow root is obstructed by the purple root, and its base is less visible than the orange base. So, the orange root should be prioritized in this frame.

### When not to label a smaller root

- If it's 75% or more occluded by another larger root

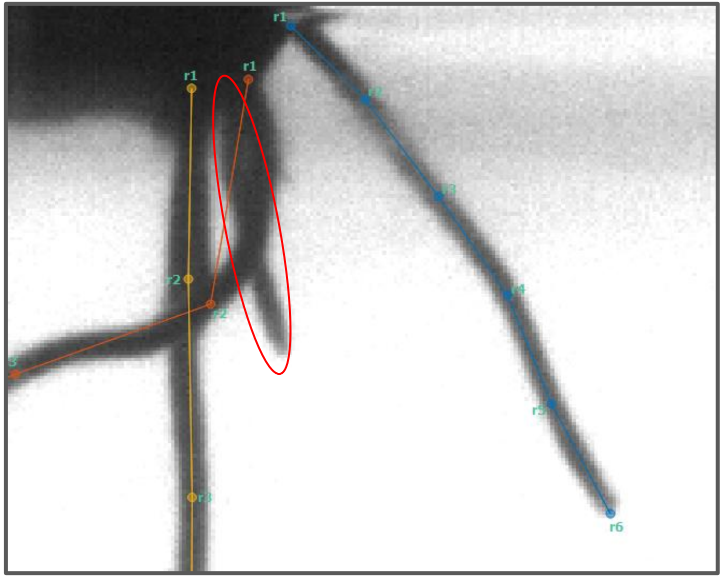


Hairy roots that shouldn't be labelled

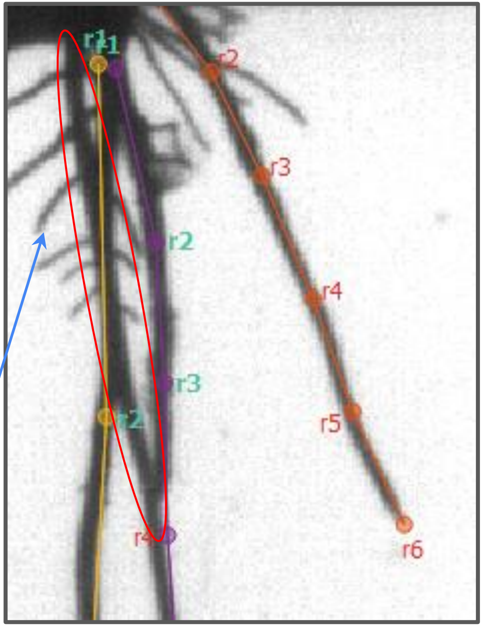


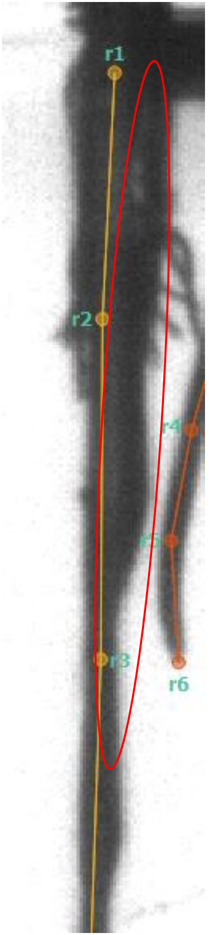

### Toggling Visibility (only when necessary) - Bases

- Toggle visibility of a node anytime it's on a part of the root that's not visible.
- If two roots' bases overlap, you can either move the base of the lower-priority root down (TRY THIS FIRST), or toggle the visibility of the nodes above its first non-overlapping point
  - For an example of moving `r1` downwards, see base definition/labelling slide

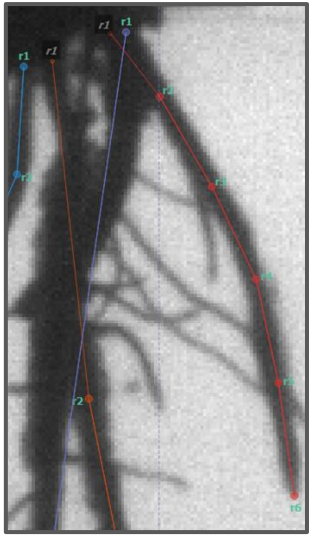


The bases of the two rightmost roots overlap. Since the purple root is both thicker and longer, it takes priority. So, the visibility of `r1` of the red root should be toggled to invisible, and `r2` should be placed at the first non-overlapping point on the red root. Use right click to toggle visibility.

### Toggling Visibility--Tips

- SLEAP uses landmark detection
  - ==> Tips are easily recognizable
- If you cannot see the tip, label the last point on the root as r5 and r6 as invisible
- Only do this if most of that root is visible (more than 75%). Otherwise, just do not label the root.

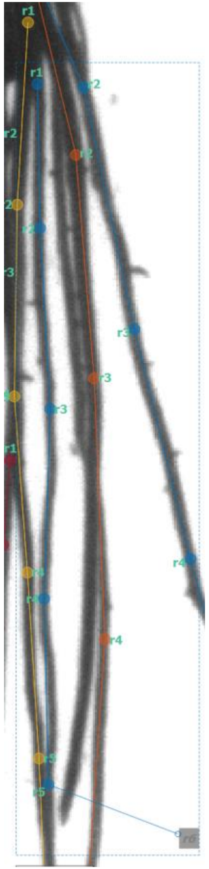

### Toggling Visibility - Holes

- If a segment in the middle of a root is not visible and 1+ nodes should be toggled invisible, the rest of the nodes until either the top or bottom (depending on which is closer or less visible) must also be toggled invisible.
    - There cannot be any holes of visibility in a root.

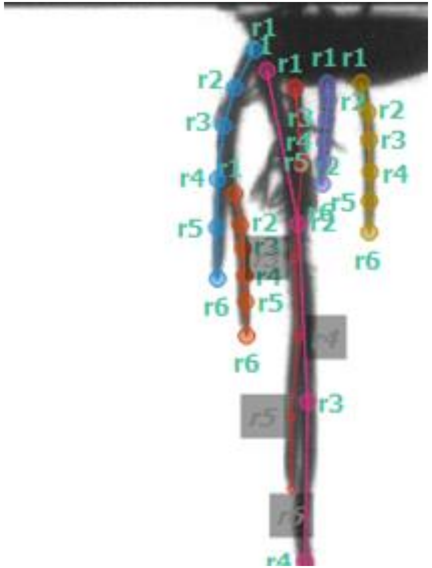

`r3` on the red root overlaps with the pink root, and in this case the pink root takes priority. So, `r3` on the red root must be toggled invisible. Since it's in the middle of the root, either `r4, r5,` and `r6` or `r1` and `r2` must also be toggled invisible to avoid a hole.

### Labeling Around Bubbles

- If a bubble is obstructing a root but you're sure the root goes under the bubble, you can place a node on top of the bubble
  - Make sure to keep nodes evenly spaced
  - If you're not sure where the root ends/if it ends behind the bubble, shift to other frames with a similar angle where the root isn't hidden behind the bubble

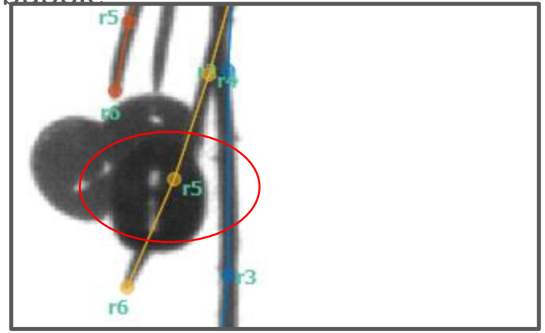


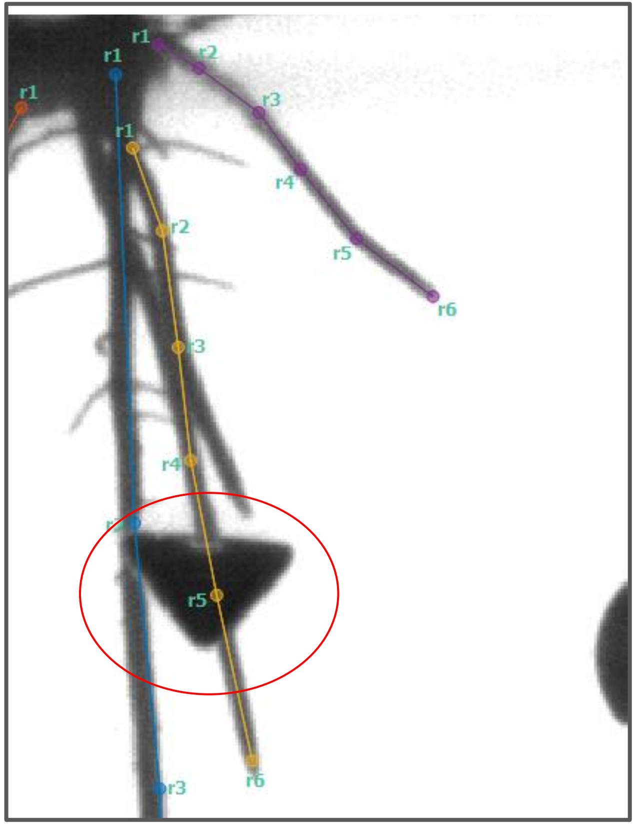

### Shoots

- This is a shoot! Do not label it.
- You can tell it is an upside-down shoot because it has sharp edges and it folds when it hits the bottom
- Label the roots only!

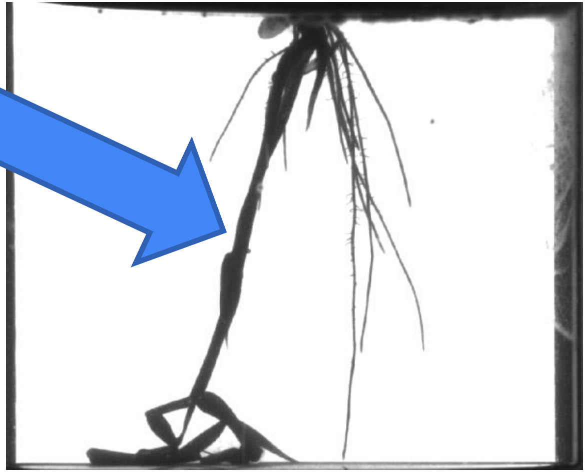

### Rice Primary Root Labeling Rules

### Primary Root

- Longest main root
- Only one root should be labeled in this model

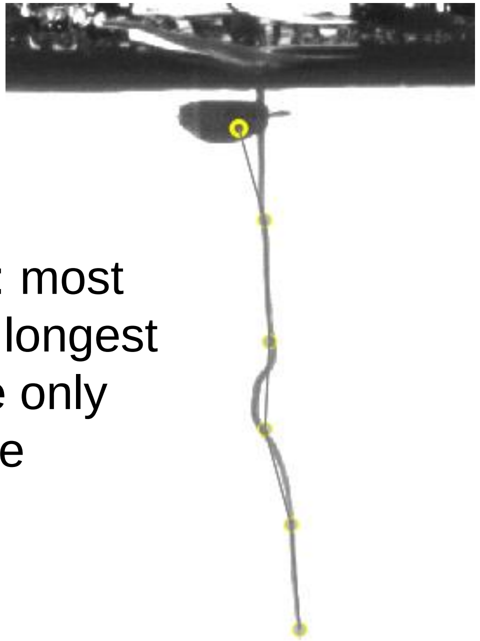

3 day-old: most times the longest root is the only root visible

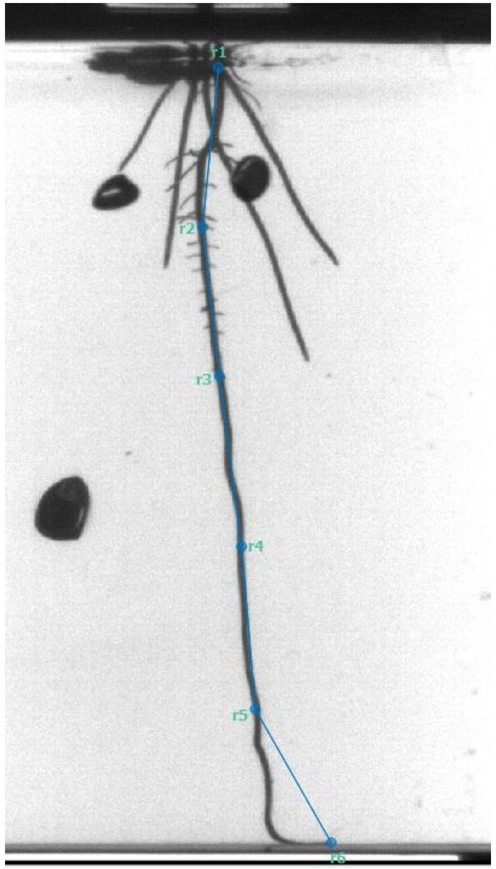


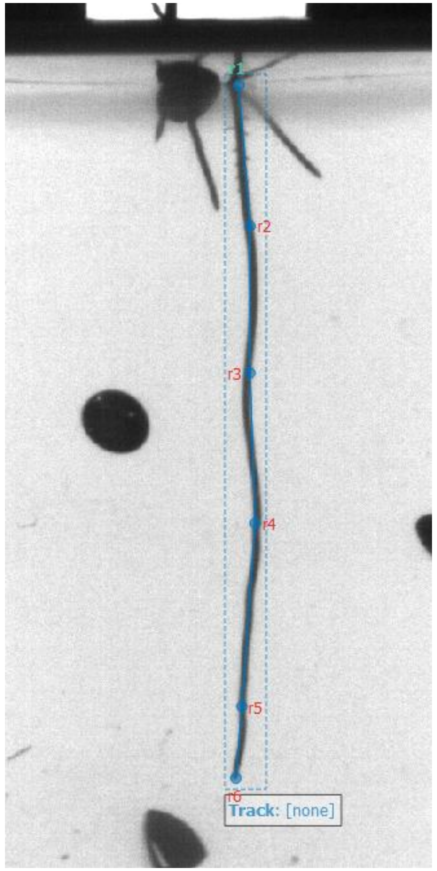

### Skeleton of rice

```
"node_names": ["r1", "r2", "r3", "r4", "r5", "r6"],
"edge_inds": [[0, 1], [1, 2], [2, 3], [3, 4], [4, 5]]
```

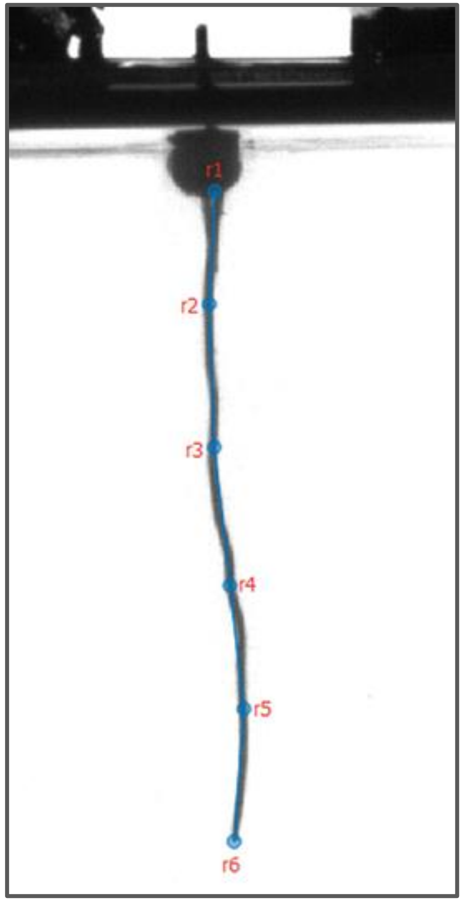


Roots are labeled using 6-node trees, which form a line.

### General Node Placement Rules

Nodes should be equally spaced along each root.

- Each 6-node instance should follow the length of one root
- `[0, 1] = [1, 2]... = [5, 6]`

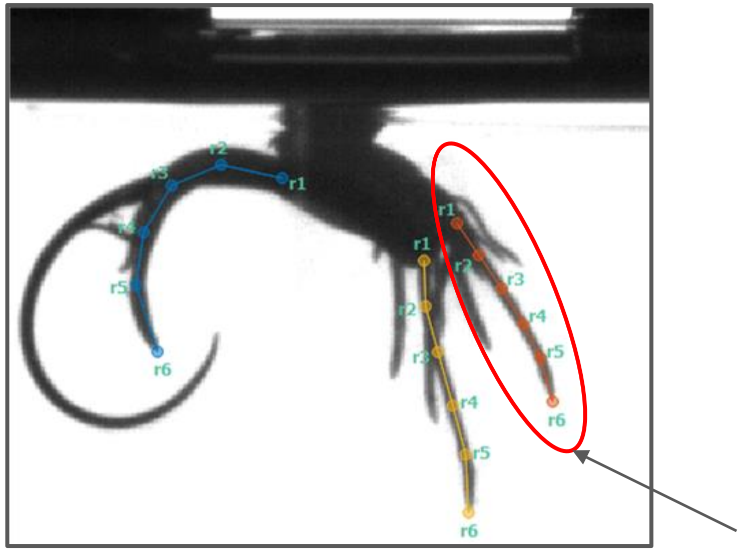


Nodes should be centered widthwise on the root.

- Zoom in to make sure they're centered

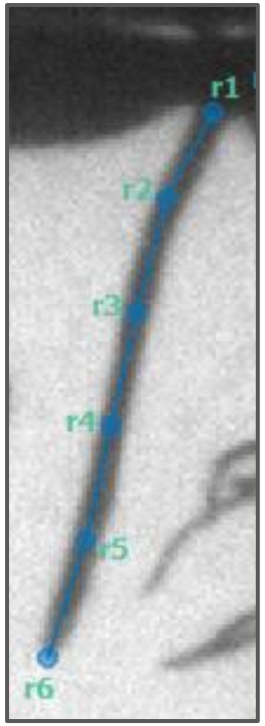

### Tip Definition and Labelling

- The tip of the root is the last visible point, and should be labelled with `r6`.
    - `Remember to zoom in`

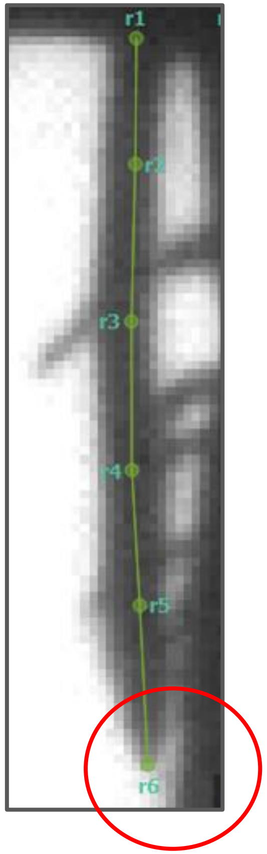

### SAVE

- Make sure to save frequently
- Changes are NOT automatically saved

### Consistency

- Consistency is key—if labels are not consistent they will have to be re-done
- If more than one person is labeling, make sure to discuss labeling preferences and special cases
  - Add new agreed-upon rules to slides to record special cases
- Go slowly and carefully rather than quickly

### Labeling Around Bubbles

- If a bubble is obstructing a root but you're sure the root goes under the bubble, you can place a node on top of the bubble
  - Make sure to keep nodes evenly spaced
  - If you're not sure where the root ends/if it ends behind the bubble, shift to other frames with a similar angle where the root isn't hidden behind the bubble

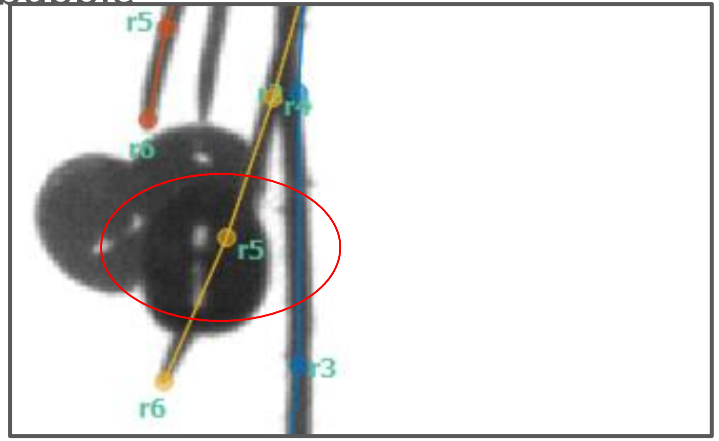


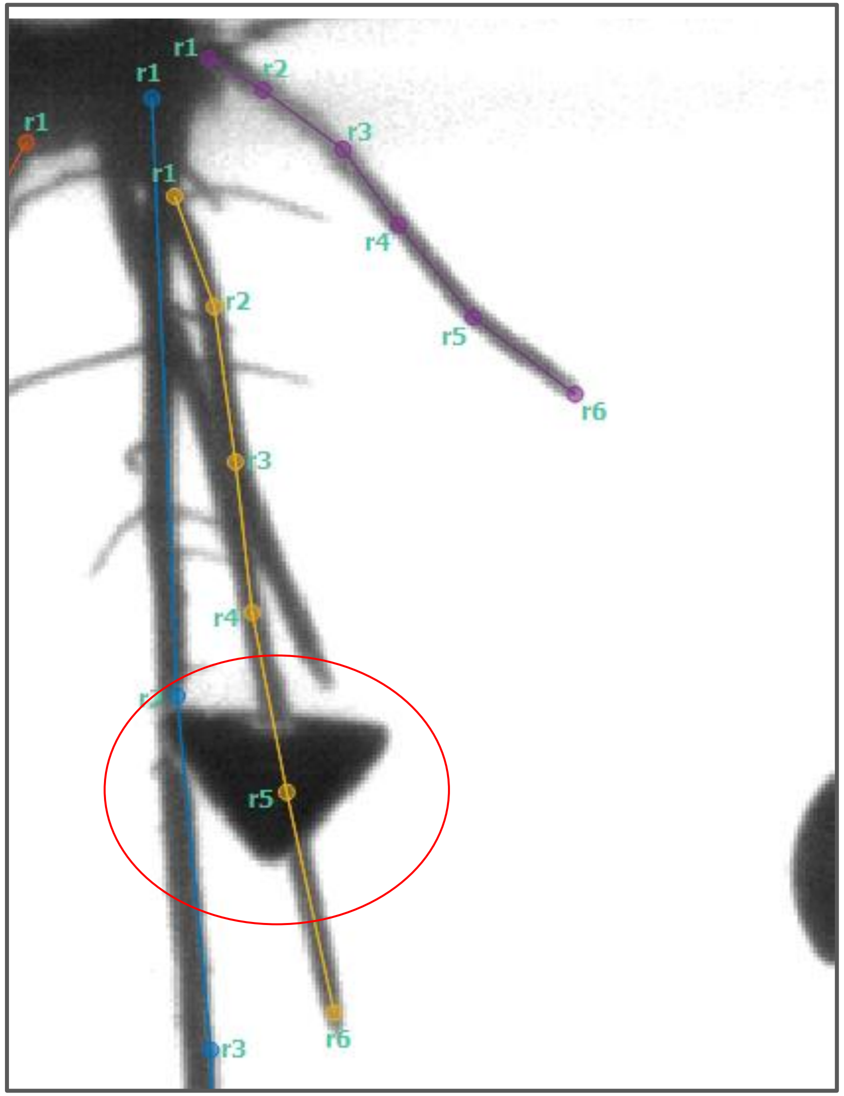

### Shoots

- This is a shoot! Do not label it.
- You can tell it is an upside-down shoot because it has sharp edges and it folds when it hits the bottom
- Label the roots only!

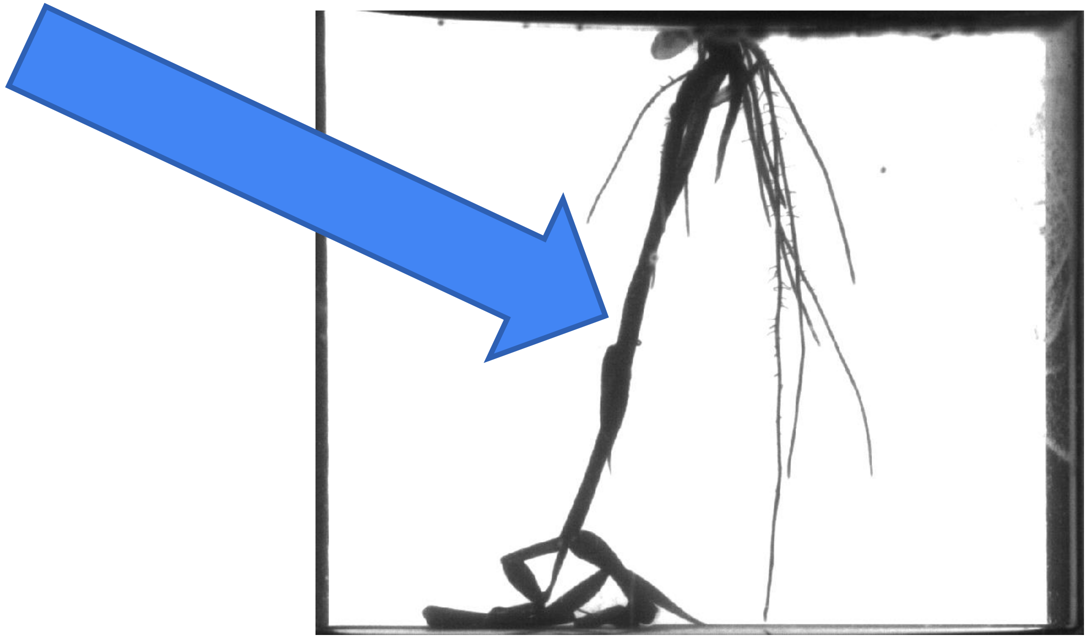

### Labeling Rules for 6 Day-Old Soy

Lateral Roots

4 nodes

### Primary Root vs. Lateral Roots

- **Primary**—usually the largest, thickest, most gravitropic root.
- **Lateral roots** grow from the primary root.

### Skeleton of soy lateral roots

```
"node_names": ["r1", "r2", "r3", "r4“]
"edge_inds": [[0, 1], [1, 2], [2, 3]]
```

Roots are labelled using 4-node trees, which form a line.

### General Node Placement Rules

Nodes should be **equally spaced** along each root.

- Each 4-node instance should follow the length of one root
- `[0, 1] = [1, 2]...`

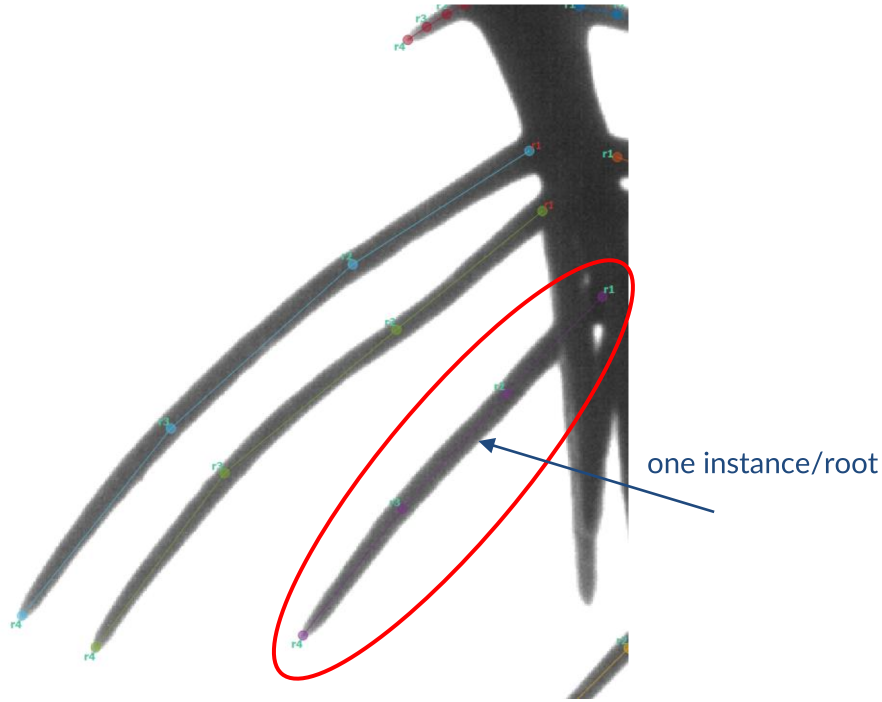


Nodes should be **centered widthwise** on the root.

- **Zoom in** to make sure they're centered

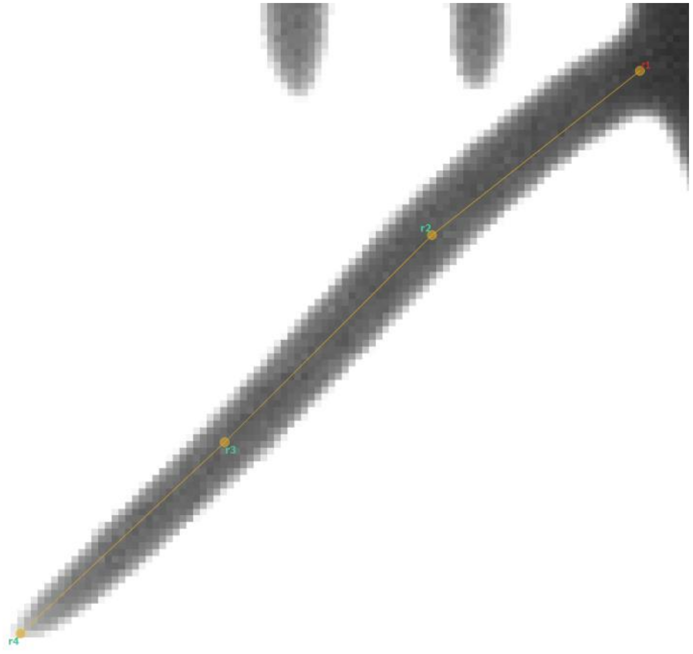

### Base Definition + Labelling Instructions

- The **base** of the root is the **first point** on a lateral root and should be labelled as `r1`.
- This is where the **lateral root** meets the **primary root**

### Tip Definition and Labelling

- The tip of the root is the last visible point, and should be labelled with `r4`.
  - `Remember to zoom in`

### SAVE

- Make sure to save frequently
- Changes are NOT automatically saved

### Consistency

- Consistency is key—if labels are not consistent they will have to be re-done
- If more than one person is labeling, make sure to discuss labeling preferences and special cases
  - Add new agreed-upon rules to slides to record special cases
- Go slowly and carefully rather than quickly

### Don't over label

- Roots can be complicated
- Prioritize larger roots
- Prioritize more visible roots
- Prioritize roots important to overall RSA
- Prioritize roots with true angle in frame
- Minimize criss-crossing labels

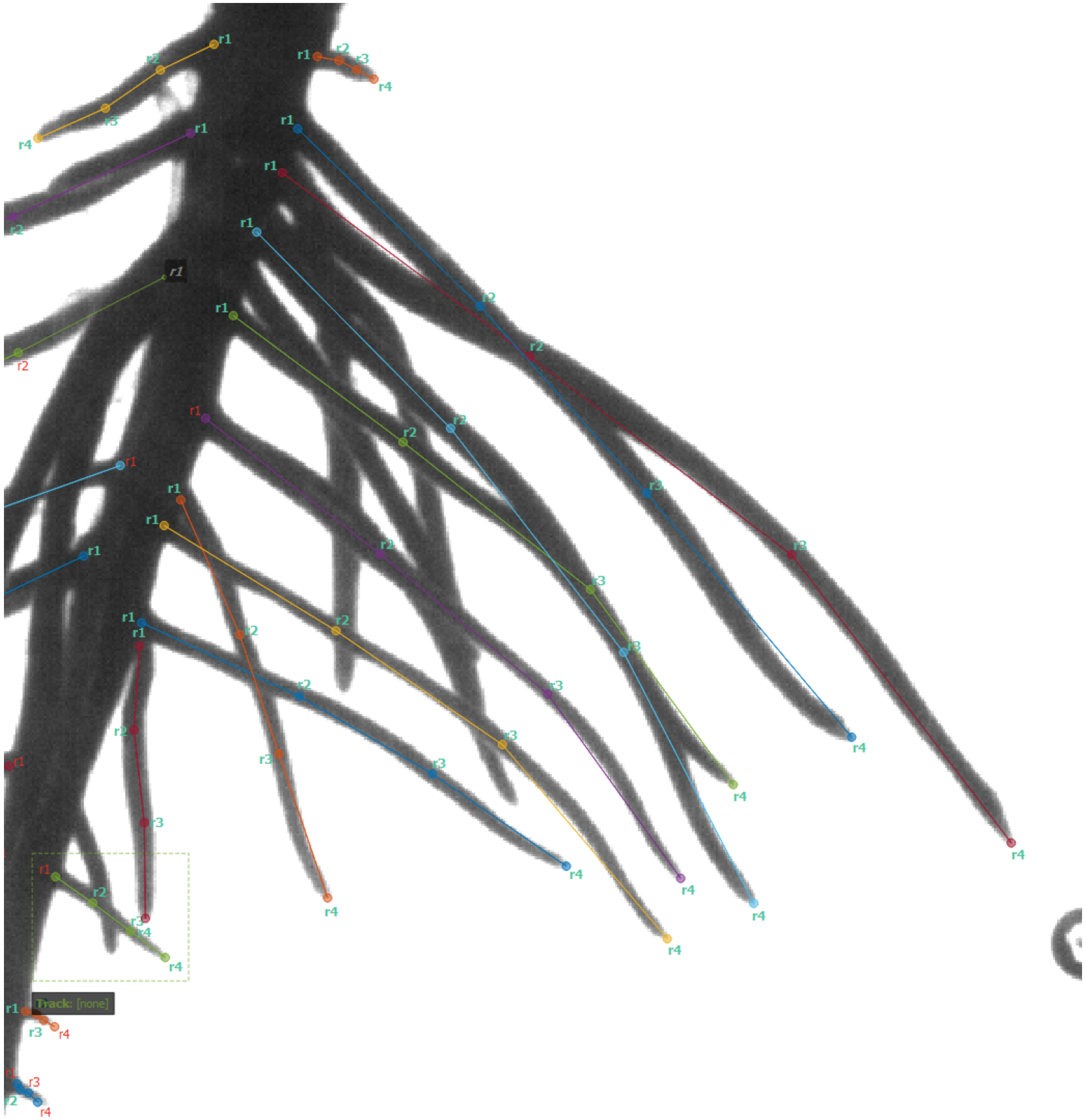
r1
r2
r3
r4
Track: [none]

### Prioritize Larger Roots

- The green root is more important than the purple root. Label the green root first and the purple root after.
- Since you cannot see the base of the purple root after the green root is labeled, mark the base of the purple root as invisible (by **right-clicking** it).

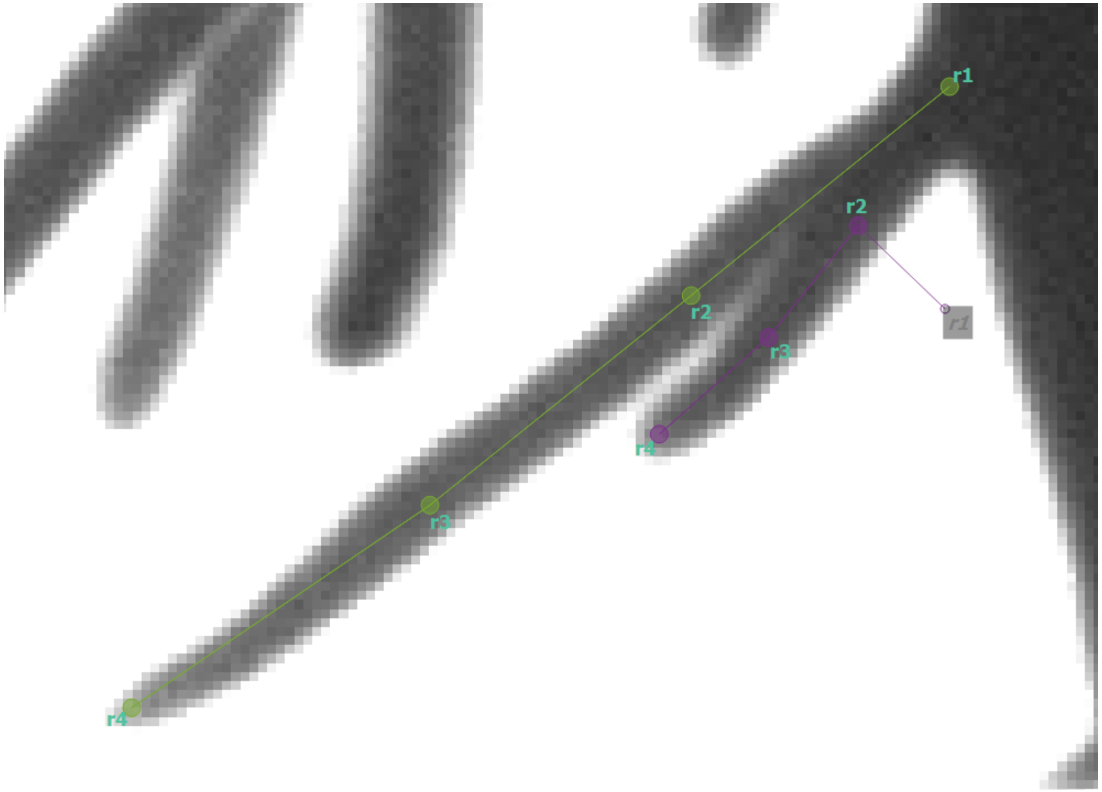

### Prioritizing Roots

- If two roots have similar thickness/length, prioritize the one that is less obstructed by other roots
  - The non-prioritized one will be prioritized in a different frame

### Prioritizing Roots--Angle

- Prioritize root with the true angle
  - Every frame shows a different angle of the plant
  - Roots viewed from the side have a smaller angle than viewed from the front
  - We want the max angle over the 72 frames

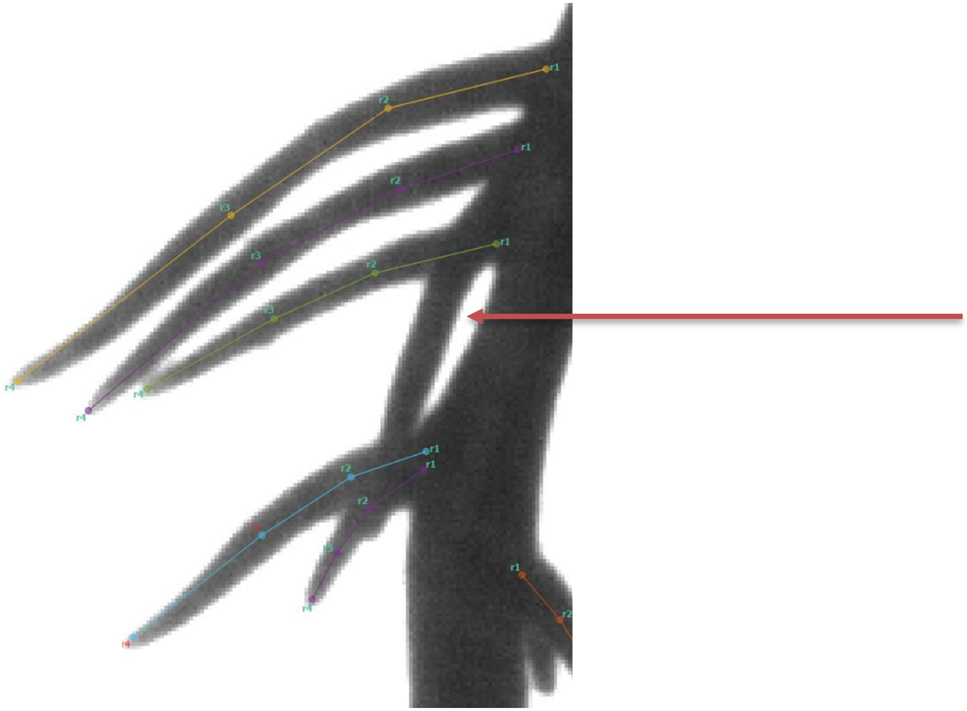

### When not to label a root

- If it's 75% or more occluded by another root

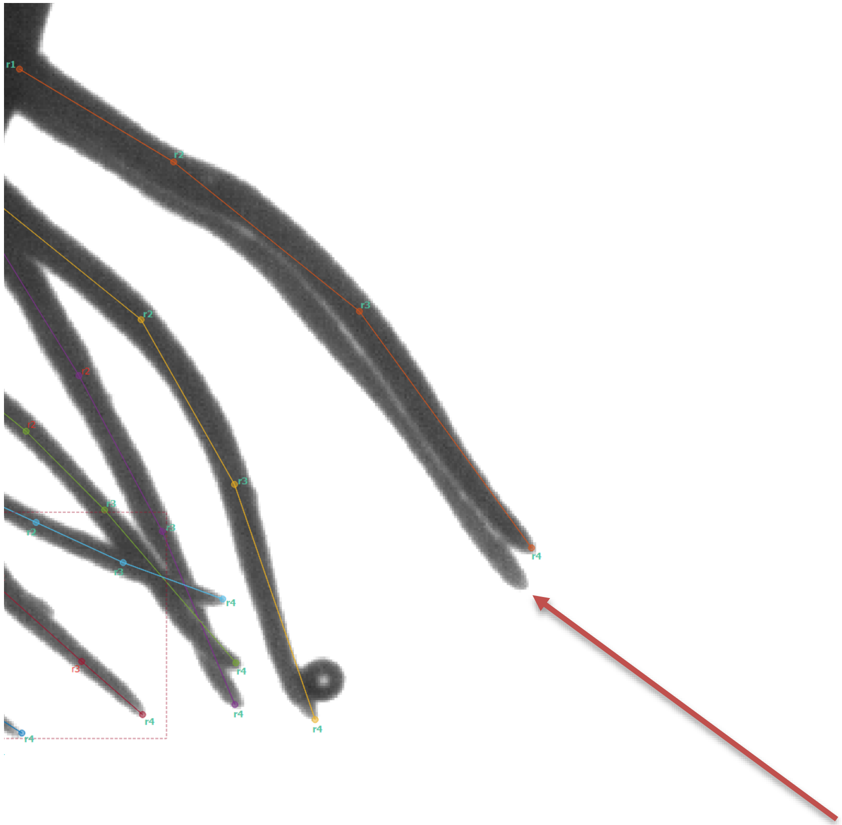

### Toggling Visibility (only when necessary) - Bases

- Toggle visibility of  a node anytime it's on a part of  the root that's not visible.

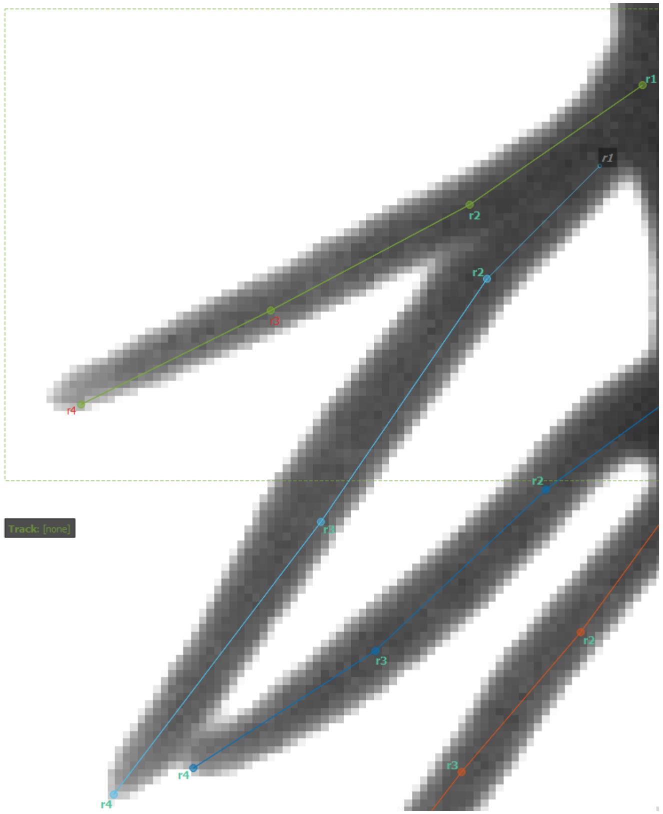

The light blue root base is behind the green root.

➔ Toggle the base of the light blue root invisible by **right – clicking it**

It will then appear grey

### Toggling Visibility – NO Holes

- If a segment in the middle of a root is not visible and 1+ nodes should be toggled invisible, the rest of the nodes until either the top or bottom (depending on which is closer or less visible) must also be toggled invisible.
  - There cannot be any holes of visibility in a root.

### Toggling Visibility--Tips

- SLEAP uses **landmark detection**
  - ==> Tips are easily recognizable
- If you cannot see the tip, label the last point on the root as r3 and r4 as invisible
- Only do this if most of that root is visible (more than 75%). Otherwise, just do not label the root.

### Labeling Around Bubbles or Contamination

- If a bubble is obstructing a root but you're **sure** the root goes under the bubble, you can place a node on top of the bubble
  - Make sure to keep nodes evenly spaced
  - If you're not sure where the root ends/if it ends behind the bubble, shift to other frames with a similar angle where the root isn't hidden behind the bubble

- You can label the base of an occluded root if you can tell where that would be
- Here the occluding root is not labeled so it's ok to label the green root's base

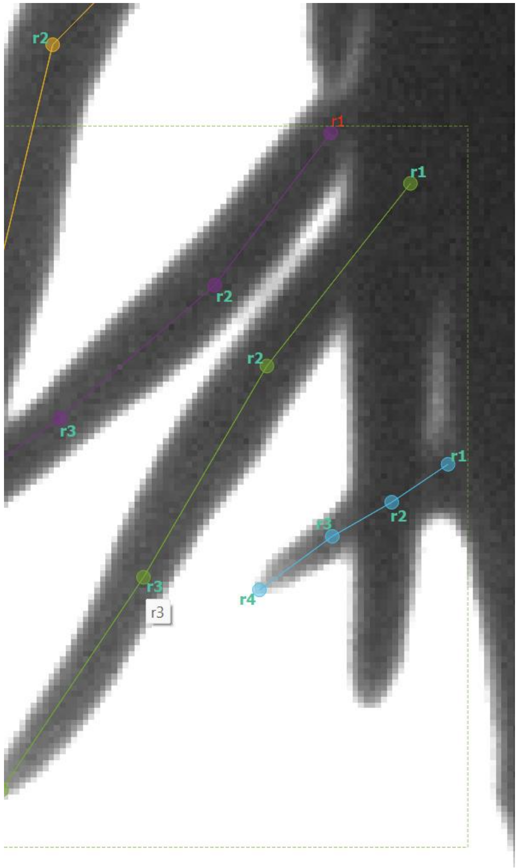

### Don't put landmarks right next to each other

- The probability fields used to find them will overlap too much.
- You can move the green base node a bit so that it is on its own root
- Or mark as invisible if too occluded to tell where the base is

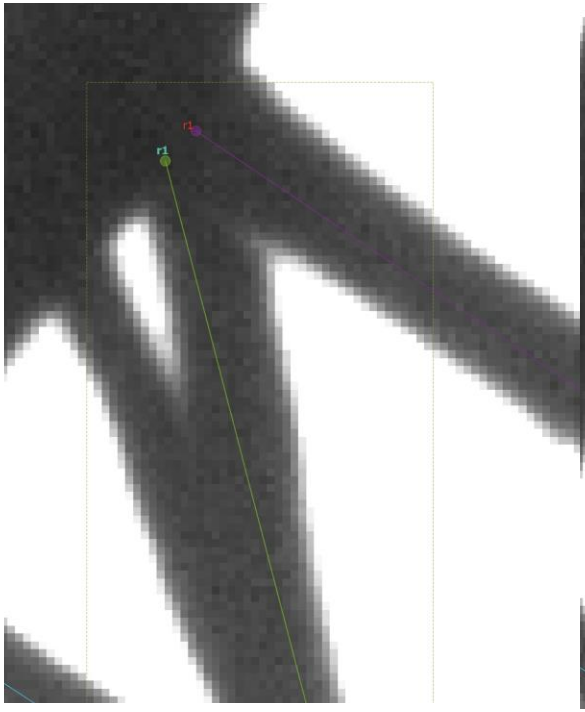

**Incorrect**

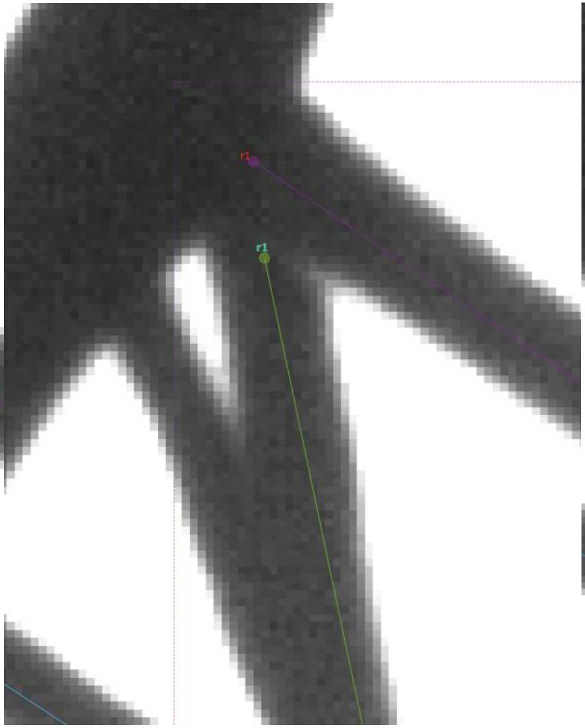

**Correct**

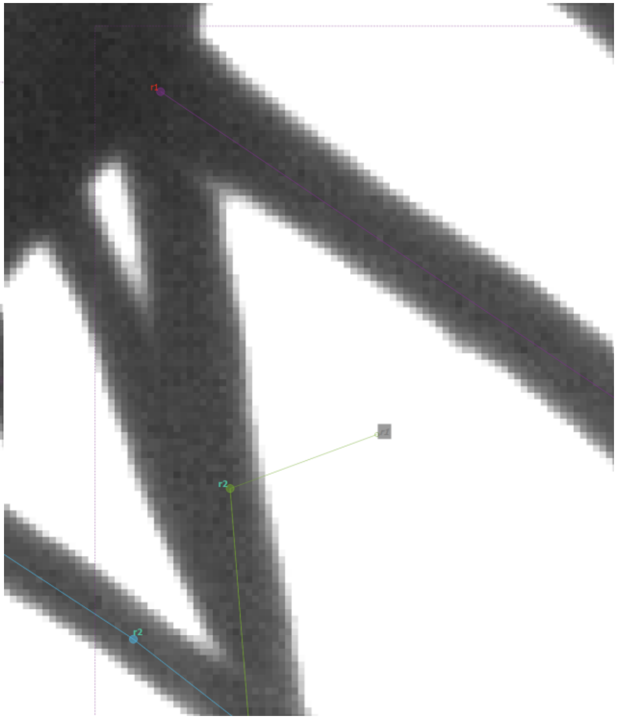

**Correct**
